## Additional File 1: supplementary materials for "Probabilistic embedding, clustering, and alignment for integrating spatial transcriptomics data with PRECAST"

[†]Equal contributions

This PDF file includes:

- Supplementary Note 1
- Supplementary Figures S1-S52

### Supplementary Note 1

### 1 PRECAST method

We first introduce the implementation details for PRECAST. Recall that the observation $\mathbf{x}_{ri} \in \mathbb{R}^p$ is an observed gene expression vector measured on tissue sample $r$ in a spatial spot $s_{ri} \in \mathbb{R}^d$, i.e., $d = 2$, and $y_{ri}$ is a latent class label, denoting cell type/spatial domain, for spot $s_{ri}$, $r = 1, \cdots, M, i = 1, \cdots, n_r$, where $M$, taking any finite positive integer, represents the number of experimental samples and $n_r$ is the number of spots for sample $r$.

We start with some notations. Let $\mathbf{y}_r = (y_{r1}, \cdots, y_{rn_r})^{\mathrm{T}}$ be the vector of latent class labels for all spots of sample $r$, $\mathbf{Z}_r = (\mathbf{z}_{r1}, \cdots, \mathbf{z}_{rn_r})^{\mathrm{T}} \in \mathbb{R}^{n_r \times q}$ be the aligned low-dimensional feature matrix of sample $r$, and $\mathbf{V}_r = (\mathbf{v}_{r1}, \cdots, \mathbf{v}_{rn_r})^{\mathrm{T}} \in \mathbb{R}^{n_r \times q}$ be the latent matrix of sample $r$ that captures the spatial dependence among neighboring spots. Let $\mathbf{0}$ represent a full-zero vector or matrix with an appropriate shape. Moreover, denote $[n_r] = \{1, \cdots, n_r\}, [n_r] \setminus i = \{j, j \neq i, j \leq n_r\}$, $n = \sum_{r=1}^{M} n_r$, and for a set $G \subset [n_r]$ and a random variable $y_{ri}$, $\mathbf{y}_{rG} = \{y_{ri}, i \in G\}$.

#### 1.1 Pseudo full/observed log-likelihood

By choosing the same number of neighbors for each spot, i.e, $m_{ri} = m$, we re-parameterize $m_{ri}^{-1}\Psi_r$ as $\tilde{\Psi}_r$. Denote $\boldsymbol{\theta} = (\mu_k, \Sigma_k, k \leq K, \mathbf{W}, \Lambda_r, \tilde{\Psi}_r, \beta_r, r \leq M)$ to be the model parameters involved. According to Models (1)–(4) in the main text, we obtain the full likelihood given by

$$P(\mathbf{X}, \mathbf{Z}, \mathbf{V}, \mathbf{y}; \boldsymbol{\theta}) = \Pi_r \left[\Pi_{i=1}^{n_r} \left\{P(\mathbf{x}_{ri}|\mathbf{z}_{ri}, \mathbf{v}_{ri}) P(\mathbf{z}_{ri}|y_{ri})\right\} P(\mathbf{y}_r) P(\mathbf{V}_r)\right], \qquad (1)$$

where $\mathbf{X} = (\mathbf{X}_1^{\mathrm{T}}, \mathbf{X}_2^{\mathrm{T}}, \cdots, \mathbf{X}_M^{\mathrm{T}})^{\mathrm{T}} \in \mathbb{R}^{n \times p}, \mathbf{Z} = (\mathbf{Z}_1^{\mathrm{T}}, \mathbf{Z}_2^{\mathrm{T}}, \cdots, \mathbf{Z}_M^{\mathrm{T}})^{\mathrm{T}} \in \mathbb{R}^{n \times q}, \mathbf{V} = (\mathbf{V}_1^{\mathrm{T}}, \mathbf{V}_2^{\mathrm{T}}, \cdots, \mathbf{V}_M^{\mathrm{T}})^{\mathrm{T}} \in \mathbb{R}^{n \times q}, \mathbf{y} = (\mathbf{y}_1^{\mathrm{T}}, \mathbf{y}_2^{\mathrm{T}}, \cdots, \mathbf{y}_M^{\mathrm{T}})^{\mathrm{T}} \in \mathbb{R}^{n \times 1}$. The full log-likelihood is given by

$$\ln P(\mathbf{X}, \mathbf{Z}, \mathbf{y}; \boldsymbol{\theta}) = \sum_r \left[\sum_i \{\ln P(\mathbf{x}_{ri}|\mathbf{z}_{ri}, \mathbf{v}_{ri}) + \ln P(\mathbf{z}_{ri}|y_{ri})\} + \ln P(\mathbf{y}_r) + \ln P(\mathbf{V}_r)\right]. \qquad (2)$$

It is very difficult to estimate the parameters due to the inter-correlation of latent variables $\mathbf{y}_r$ and $\mathbf{V}_r$, and use of conventional EM algorithms is not feasible because $E\{\ln P(\mathbf{y}_r) + \ln P(\mathbf{V}_r)|\mathbf{X}\}$ has a complex form, where the expectation is taken on $(\mathbf{y}_r, \mathbf{Z}_r, \mathbf{V}_r)$ given $\mathbf{X}$. Thus, a pseudo-likelihood technique is used to replace the joint likelihood of class labels $\mathbf{y}_r$ and latent features $\mathbf{V}_r$ with a pseudo likelihood, making the joint one separable. This technique was the key to making the computation tractable.

Following Besag [1], suppose that we have a prediction of $\mathbf{y}_r$ and $\mathbf{V}_r$, denoted by $\hat{\mathbf{y}}_r$ and $\widehat{\mathbf{V}}_r$, then the pseudo likelihoods of $\mathbf{y}_r$ and $\mathbf{V}_r$ are defined as

$$\tilde{P}(\mathbf{y}_r; \beta_r) = \Pi_i P(y_{ri}|\hat{\mathbf{y}}_{N_{ri}}), \qquad (3)$$
$$\tilde{P}(\mathbf{V}_r; \tilde{\Psi}_r) = \Pi_i P(\mathbf{v}_{ri}|\hat{\mathbf{v}}_{N_{ri}}). \qquad (4)$$

In a later subsection, we discuss how to obtain $\hat{\mathbf{y}}_r$ and $\widehat{\mathbf{V}}_r$, which are also updated iteratively.

Combining (1), (3) and (4), we obtain the pseudo full likelihood

$$\tilde{P}(\mathbf{y}, \mathbf{X}, \mathbf{Z}; \boldsymbol{\theta}) = \Pi_r \Pi_{i=1}^{n_r} P(\mathbf{x}_{ri}|\mathbf{z}_{ri}, \mathbf{v}_{ri}) P(\mathbf{z}_{ri}|y_{ri}) P(y_{ri}|\hat{y}_{N_{ri}}) P(\mathbf{v}_{ri}|\hat{\mathbf{v}}_{N_{ri}}).$$

We will show this pseudo full log likelihood is feasible. Furthermore, we obtain the pseudo observed log-likelihood given by

$$\ln \tilde{P}(\mathbf{X};\boldsymbol{\theta}) = \sum_r \sum_i \ln \sum_k P(\mathbf{x}_{ri}|y_{ri}=k;\boldsymbol{\theta})P(y_{ri}=k|\hat{y}_{N_{ri}};\boldsymbol{\theta}), \tag{5}$$

where $\mathbf{x}_{ri}|y_{ri}=k \sim N(\mathbf{W}(\mu_k+\mu_{v_{ri}}), \mathbf{W}(\tilde{\Psi}_r+\Sigma_k)W^{\mathrm{T}}+\Lambda_r)$. Based on the pseudo observed log-likelihood (5), we developed an EM-type algorithm, called ICM-EM, to estimate the parameters and achieve embedding alignment, dimension reduction and spatial clustering simultaneously. The ICM-EM is derived to maximize the lower-bound function of the pseudo observed log-likelihood.

#### 1.2 ICM-EM algorithm

##### 1.2.1 ICM step to predict y and $V$

Given all model parameters, we can obtain the predictions $\hat{\mathbf{y}}$ and $\widehat{\mathbf{V}}$ by maximizing the posterior of $\mathbf{y}$ and $\mathbf{V}$, $P(\mathbf{y},\mathbf{V}|\mathbf{X};\boldsymbol{\theta})$. However, the joint distribution $P(\mathbf{y},\mathbf{V}|\mathbf{X};\boldsymbol{\theta})$ is extremely complex, so we apply iterative conditional mode (ICM, [2]) method to alternately predict $y_{ri}$ and $\mathbf{v}_{ri}$ for each sample $r$. We call this the ICM step, which composes an important part of ICM-EM.

First, given $\mathbf{V}=\widehat{\mathbf{V}}$, we predict $\mathbf{y}$ based on ICM. Using Bayes's formula, we obtain

$$P(y_{ri}|\mathbf{X};\hat{\mathbf{y}}_{[n_r]\backslash i},\hat{\mathbf{V}}) \propto P(\mathbf{x}_{ri}|y_{ri};\hat{\mathbf{v}}_{N_{ri}})P(y_{ri}|\hat{\mathbf{y}}_{N_{ri}}).$$

Thus, we can predict $y_{ri}$ by

$$\hat{y}_{ri} = \arg\max_{y_{ri}\in\{1,\cdots,K\}} P(\mathbf{x}_{ri}|y_{ri};\hat{\mathbf{v}}_{N_{ri}})P(y_{ri}|\hat{\mathbf{y}}_{N_{ri}}), \tag{6}$$

where $(\mathbf{x}_{ri}|y_{ri}=k;\hat{\mathbf{v}}_{N_{ri}}) \sim N(\mathbf{W}(\mu_k+\hat{\mu}_{v_{ri}}), \mathbf{W}(\tilde{\Psi}_r+\Sigma_k)\mathbf{W}^{\mathrm{T}}+\Lambda_r)$.

Next, we predict $\mathbf{V}$ given $\hat{\mathbf{y}}$. Similarly, from Bayes's formula, we know

$$P(\mathbf{v}_{ri}|\mathbf{X},\hat{\mathbf{y}},\hat{\mathbf{v}}_{[n_r]\backslash i}) \propto P(\mathbf{x}_{ri}|\hat{y}_{ri},\mathbf{v}_{ri})P(\mathbf{v}_{ri}|\hat{\mathbf{v}}_{N_{ri}}).$$

Moreover, we have

$$\begin{aligned} & \ln P(\mathbf{x}_{ri}|\hat{y}_{ri},\mathbf{v}_{ri})P(\mathbf{v}_{ri}|\hat{\mathbf{v}}_{N_{ri}}) \\ = & -\frac{1}{2}(\mathbf{x}_{ri}-\mathbf{W}(\mu_k+\mathbf{v}_{ri}))^{\mathrm{T}}(\mathbf{W}\Sigma_k\mathbf{W}^{\mathrm{T}}+\Lambda_r)^{-1}(\mathbf{x}_{ri}-\mathbf{W}(\mu_k+\mathbf{v}_{ri})) \\ - & \frac{1}{2}(\mathbf{v}_{ri}-\mu_{v_{ri}})^{\mathrm{T}}\tilde{\Psi}_r^{-1}(\mathbf{v}_{ri}-\mu_{v_{ir}}), \end{aligned}$$

so we obtain

$$\begin{aligned} \hat{\mathbf{v}}_{ri} &= \arg\max_{\mathbf{v}_{ri}} \ln P(\mathbf{x}_{ri}|\hat{y}_{ri},\mathbf{v}_{ri})P(\mathbf{v}_{ri}|\hat{\mathbf{v}}_{N_{ri}}) \\ &= (\mathbf{W}^{\mathrm{T}}\mathbf{S}_{rk}\mathbf{W}+\tilde{\Psi}_r^{-1})^{-1}(\mathbf{W}^{\mathrm{T}}\mathbf{S}_{rk}(\mathbf{x}_i-\mathbf{W}\mu_k)+\tilde{\Psi}_r^{-1}\mu_{v_{ri}}), \end{aligned} \tag{7}$$

where $\mathbf{S}_{rk}=(\mathbf{W}\Sigma_k\mathbf{W}^{\mathrm{T}}+\Lambda_r)^{-1}=\Lambda_r^{-1}-\Lambda_r^{-1}\mathbf{W}\mathbf{C}_{rk}^{-1}\mathbf{W}^{\mathrm{T}}\Lambda_r^{-1}$ and $\mathbf{C}_{rk}=\mathbf{W}^{\mathrm{T}}\Lambda_r^{-1}\mathbf{W}+\Sigma_k^{-1}$. We define the total objective function with respect to $(\mathbf{V}_r,\mathbf{y}_r)$ for sample $r$ as

$$\Xi_r(\mathbf{V},\mathbf{y};\widehat{\mathbf{V}},\hat{\mathbf{y}}) = \sum_{i=1}^{n_r}\left[\ln\left\{P(\mathbf{x}_{ri}|y_{ri};\hat{\mathbf{v}}_{N_{ri}})P(y_{ri}|\hat{\mathbf{y}}_{N_{ri}})\right\}+\ln P(\mathbf{x}_{ri}|\hat{y}_{ri},\mathbf{v}_{ri})P(\mathbf{v}_{ri}|\hat{\mathbf{v}}_{N_{ri}})\right], \tag{8}$$

which will be used for the stop rule in the ICM step.

##### 1.2.2 E-step

In this subsection, we derive the lower-bound function of the pseudo observed log-likelihood, called the Q-function (with respect to $\boldsymbol{\theta}$), by taking the expectation of the pseudo full log-likelihood given the model parameters $\boldsymbol{\theta}^{(t)}$ obtained by $t$-th iteration. First, we have

$$
\begin{aligned}
\ln \tilde{P}(\mathbf{X};\boldsymbol{\theta}) &= \ln \int_{\mathbf{y}}\int_{\mathbf{V}}\int_{\mathbf{Z}} \tilde{P}(\mathbf{y},\mathbf{X},\mathbf{Z},\mathbf{V};\boldsymbol{\theta})d\mathbf{y}d\mathbf{Z}d\mathbf{V} \\
&= \sum_{r,i} \ln \int_{y_{ri}}\int_{\mathbf{z}_{ri}}\int_{\mathbf{v}_{ri}} P(\mathbf{x}_{ri}|\mathbf{z}_{ri},\mathbf{v}_{ri};\boldsymbol{\theta})P(\mathbf{z}_{ri}|y_{ri};\boldsymbol{\theta})P(y_{ri}|\hat{y}_{N_{ri}};\boldsymbol{\theta}) \\
&\qquad \times P(\mathbf{v}_{ri}|\hat{\mathbf{v}}_{N_{ri}};\boldsymbol{\theta})dy_{ri}d\mathbf{z}_{ri}d\mathbf{v}_{ri} \\
&\geq \sum_{r,i} \mathrm{E}_{\boldsymbol{\theta}^{(t)}} \left\{ \ln \frac{P(\mathbf{x}_{ri}|\mathbf{z}_{ri},\mathbf{v}_{ri};\boldsymbol{\theta})P(\mathbf{z}_{ri}|y_{ri};\boldsymbol{\theta})P(y_{ri}|\hat{y}_{N_{ri}};\boldsymbol{\theta})P(\mathbf{v}_{ri}|\hat{\mathbf{v}}_{N_{ri}};\boldsymbol{\theta})}{P(y_{ri},\mathbf{z}_{ri},\mathbf{v}_{ir}|\mathbf{x}_{ri};\hat{y}_{N_{ri}},\hat{\mathbf{v}}_{N_{ri}}\boldsymbol{\theta}^{(t)})} \right\}, \quad (9)
\end{aligned}
$$

where expectation $\mathrm{E}_{\boldsymbol{\theta}^{(t)}}$ is taken with respect to $(y_{ri},\mathbf{z}_{ri},\mathbf{v}_{ri})$ given $\mathbf{x}_{ri},\hat{y}_{N_{ri}},\hat{\mathbf{v}}_{N_{ri}}$ and $\boldsymbol{\theta}^{(t)}$, and the inequality follows from the Jensen's inequality. Next, to obtain the specific form of Q-function, we derive the explicit form of $P(y_{ri},\mathbf{z}_{ri},\mathbf{v}_{ir}|\mathbf{x}_{ri};\hat{y}_{N_{ri}},\hat{\mathbf{v}}_{N_{ri}},\boldsymbol{\theta}^{(t)})$ for evaluating the expectation in equation (9).

We define the responsibility that component $k$ takes for explaining the observation $\mathbf{x}_{ri}$

$$
\begin{aligned}
R_{rik} &= P(y_{ri}=k|\mathbf{x}_{ri};\hat{\mathbf{y}}_{[n]\backslash ri},\widehat{\mathbf{V}}) \\
&= \frac{P(\mathbf{x}_{ri}|y_{ri}=k;\hat{\mathbf{v}}_{N_{ri}},\boldsymbol{\theta})P(y_{ri}=k;\hat{\mathbf{y}}_{N_{ri}})}{\sum_{k'} P(\mathbf{x}_{ri}|y_{ri}=k';\hat{\mathbf{v}}_{N_{ri}},\boldsymbol{\theta})P(y_{ri}=k';\hat{\mathbf{y}}_{N_{ri}})}, \quad (10)
\end{aligned}
$$

where $P(y_{ri}=k;\hat{\mathbf{y}}_{N_{ri}}) = P(y_{ri}=k|\mathbf{y}_{N_{ri}}=\hat{\mathbf{y}}_{N_{ri}})$. Note that $R_{rik}$ is also the pseudo posterior probability of $y_{ri}$. Then, we have

$$
\begin{aligned}
& P(y_{ri}=k,\mathbf{z}_{ri},\mathbf{v}_{ri}|\mathbf{x}_{ri};\hat{\mathbf{y}}_{N_{ri}},\hat{\mathbf{v}}_{N_{ri}},\boldsymbol{\theta}) \\
=\ & P(y_{ri}=k|\mathbf{x}_{ri};\hat{\mathbf{y}}_{N_{ri}},\hat{\mathbf{v}}_{N_{ri}})P(\mathbf{z}_{ri},\mathbf{v}_{ri}|\mathbf{x}_{ri},y_{ri}=k;\hat{\mathbf{y}}_{N_{ri}},\hat{\mathbf{v}}_{N_{ri}}) \\
=\ & R_{rik}P(\mathbf{z}_{ri},\mathbf{v}_{ri}|\mathbf{x}_{ri},y_{ri}=k;\hat{\mathbf{y}}_{N_{ri}},\hat{\mathbf{v}}_{N_{ri}}),
\end{aligned}
$$

thus, we only require to derive the conditional distribution $(\mathbf{z}_{ri},\mathbf{v}_{ri}|\mathbf{x}_{ri},y_{ri}=k)$ given the predictions $(\hat{\mathbf{y}}_{N_{ri}},\hat{\mathbf{v}}_{N_{ri}})$.

Denote $\bar{\mathbf{S}}_{rk} = \left\{\mathbf{W}(\tilde{\Psi}_r+\Sigma_k)\mathbf{W}^{\mathrm{T}}+\Lambda_r\right\}^{-1} = \Lambda_r^{-1}-\Lambda_r^{-1}\mathbf{W}\bar{\mathbf{C}}_{rk}^{-1}\mathbf{W}^{\mathrm{T}}\Lambda_r^{-1}$ with $\bar{\mathbf{C}}_{rk} = \mathbf{W}^{\mathrm{T}}\Lambda_r^{-1}\mathbf{W}+(\tilde{\Psi}_r+\Sigma_k)^{-1}$. Since $(\mathbf{z}_{ri},\mathbf{v}_{ri},\mathbf{x}_{ri}|y_{ri}=k)$ is a multivariate normal distribution, we use the formula for the conditional expectation and covariance of multivariate normal distribution. Conditional on $y_{ri}=k$, we have

$$
\begin{aligned}
\mathrm{E}_k\{(\mathbf{z}_{ri},\mathbf{v}_{ri})|\mathbf{x}_{ri}\} &= (\mu_k;\mu_{v_{ri}}) + (\Sigma_k;\tilde{\Psi}_r)\mathbf{W}^{\mathrm{T}}\bar{\mathbf{S}}_{rk}\{\mathbf{x}_{ri}-\mathbf{W}(\mu_k+\mu_{v_{ri}})\}, \\
\mathrm{var}_k\{(\mathbf{z}_{ri},\mathbf{v}_{ri})|\mathbf{x}_{ri}\} &= \mathrm{diag}(\Sigma_k,\tilde{\Psi}_r) - \Sigma_{(z,v)x,rk}\bar{\mathbf{S}}_{rk}\Sigma_{(z,v)x,rk}^{\mathrm{T}},
\end{aligned}
$$

where $\mathrm{E}_k(\mathbf{z}_{ri}) \equiv E(\mathbf{z}_{ri}|y_{ri}=k), \mathrm{var}_k(\mathbf{z}_{ri}) \equiv var(\mathbf{z}_{ri}|y_{ri}=k)$ and $\Sigma_{(z,v)x,rk} = (\Sigma_k;\tilde{\Psi}_r)\mathbf{W}^{\mathrm{T}} \in \mathbb{R}^{2q\times p}$.

Based on the conditional joint distribution of $(\mathbf{z}_{ri}, \mathbf{v}_{ri})$ given $(\mathbf{x}_{ri}, y_{ri} = k)$, we derive some quantities used for the M-step. First, we have

$$\begin{aligned}
\mathrm{E}_k(\mathbf{z}_{ri}|\mathbf{x}_{ri}) &= \mu_k + \Sigma_k \mathbf{W}^{\mathrm{T}} \bar{\mathbf{S}}_{rk}(\mathbf{x}_{ri} - \mathbf{W}(\mu_k + \mu_{v_{ri}})) \\
\mathrm{var}_k(\mathbf{z}_{ri}|\mathbf{x}_{ri}) &= \Sigma_k - \Sigma_k \mathbf{W}^{\mathrm{T}} \bar{\mathbf{S}}_{rk} \mathbf{W} \Sigma_k, \\
\mathrm{E}_k(\mathbf{v}_{ri}|\mathbf{x}_{ri}) &= \mu_{v_{ri}} + \tilde{\Psi}_r \mathbf{W}^{\mathrm{T}} \bar{\mathbf{S}}_{rk}(\mathbf{x}_{ri} - \mathbf{W}(\mu_k + \mu_{v_{ri}})) \\
\mathrm{var}_k(\mathbf{v}_{ri}|\mathbf{x}_{ri}) &= \tilde{\Psi}_r - \tilde{\Psi}_r \mathbf{W}^{\mathrm{T}} \bar{\mathbf{S}}_{rk} \mathbf{W} \tilde{\Psi}_r.
\end{aligned}$$

Moreover,

$$\mathbf{z}_{ri} + \mathbf{v}_{ri}|\mathbf{x}_{ri}, y_{ri} = k \sim N(\mu_{zv,rk}, \Sigma_{zv,rk}),$$

where

$$\mu_{zv,rk} = \mu_k + \Sigma_k \mathbf{W}^{\mathrm{T}} \bar{\mathbf{S}}_{rk}(\mathbf{x}_{ri} - \mathbf{W}(\mu_k + \mu_{v_{ri}})) + \mu_{v_{ri}} + \tilde{\Psi}_r \mathbf{W}^{\mathrm{T}} \bar{\mathbf{S}}_{rk}(\mathbf{x}_{ri} - \mathbf{W}(\mu_k + \mu_{v_{ri}})),$$

and

$$\Sigma_{zv,rk} = \Sigma_k - \Sigma_k \mathbf{W}^{\mathrm{T}} \bar{\mathbf{S}}_{rk} \mathbf{W} \Sigma_k + \tilde{\Psi}_r - \tilde{\Psi}_r \mathbf{W}^{\mathrm{T}} \mathbf{S}_{rk} \mathbf{W} \tilde{\Psi}_r - \tilde{\Psi}_r \mathbf{W}^{\mathrm{T}} \bar{\mathbf{S}}_{rk} W \Sigma_k - \Sigma_k \mathbf{W}^{\mathrm{T}} \bar{\mathbf{S}}_{rk} \mathbf{W} \tilde{\Psi}_r.$$

Next, we provide some notations for the posterior expectation and covariance of latent random variables. Denote

$$\begin{aligned}
\langle \mathbf{z}_{ri} \rangle_k^{(t)} &= \mathrm{E}_k(\mathbf{z}_{ri}|\mathbf{x}_{ri}; \boldsymbol{\theta}^{(t)}), \\
\langle \mathbf{z}_{ri} \mathbf{z}_{ri}^{\mathrm{T}} \rangle_k^{(t)} &= \mathrm{var}_k(\mathbf{z}_{ri}|\mathbf{x}_{ri}; \boldsymbol{\theta}^{(t)}) + \mathrm{E}_k(\mathbf{z}_{ri}|\mathbf{x}_{ri}; \boldsymbol{\theta}^{(t)}) \mathrm{E}_k(\mathbf{z}_{ri}|\mathbf{x}_{ri} \boldsymbol{\theta}^{(t)})^{\mathrm{T}}, \\
\langle \mathbf{v}_{ri} \rangle_k^{(t)} &= \mathrm{E}_k(\mathbf{v}_{ri}|\mathbf{x}_{ri}; \boldsymbol{\theta}^{(t)}), \\
\langle \mathbf{v}_{ri} \mathbf{v}_{ri}^{\mathrm{T}} \rangle_k^{(t)} &= \mathrm{var}_k(\mathbf{v}_{ri}|\mathbf{x}_{ri}; \boldsymbol{\theta}^{(t)}) + \mathrm{E}_k(\mathbf{v}_{ri}|\mathbf{x}_{ri}; \boldsymbol{\theta}^{(t)}) \mathrm{E}_k(\mathbf{v}_{ri}|\mathbf{x}_{ri}; \boldsymbol{\theta}^{(t)})^{\mathrm{T}}, \\
\langle \mathbf{v}_{ri} + \mathbf{z}_{ri} \rangle_k^{(t)} &= \mu_{zv,rk}, \\
\langle (\mathbf{v}_{ri} + \mathbf{z}_{ri})(\mathbf{v}_{ri} + \mathbf{z}_{ri})^{\mathrm{T}} \rangle_k^{(t)} &= \Sigma_{zv,rk} + \mu_{zv,rk} \mu_{zv,rk}^{\mathrm{T}}.
\end{aligned}$$

Furthermore, on the basis of the above derivation, we give the explicit form of Q-function by

$$\begin{aligned}
Q(\boldsymbol{\theta}; \boldsymbol{\theta}^{(t)}, \hat{\mathbf{y}}, \widehat{V}) &= \sum_{r,i,k} R_{rik}^{(t)} \mathrm{E}_{(\mathbf{z}_{ri}, \mathbf{v}_{ri})|\mathbf{x}_{ri}, y_{ri}=k; \boldsymbol{\theta}^{(t)}} \ln P(\mathbf{x}_{ri}|\mathbf{z}_{ri}, \mathbf{v}_{ri}; \boldsymbol{\theta}) \\
&+ \sum_{r,i,k} R_{rik}^{(t)} \mathrm{E}_{\mathbf{z}_{ri}|\mathbf{x}_{ri}, y_{ri}=k; \boldsymbol{\theta}^{(t)}} \ln P(\mathbf{z}_{ri}|y_{ri} = k; \boldsymbol{\theta}) \\
&+ \sum_{r,i,k} R_{rik}^{(t)} \mathrm{E}_{\mathbf{v}_{ri}|\mathbf{x}_{ri}, y_{ri}=k; \boldsymbol{\theta}^{(t)}} \ln P(\mathbf{v}_{ri}|\hat{\mathbf{v}}_{N_{ri}}; \tilde{\Psi}_r) \\
&+ \sum_{r,i,k} R_{rik}^{(t)} \ln P(y_{ri} = k|\hat{\mathbf{y}}_{N_{ri}}; \beta_r) \\
&= \sum_{r,i} (I_{ri1} + I_{ri2} + I_{ri3} + I_{ri4}) + const,
\end{aligned}$$

where $R_{rik}^{(t)}$ is the value of $R_{rik}$ when $\boldsymbol{\theta} = \boldsymbol{\theta}^{(t)}$, *const* is a constant term independent of parameters, and

$$\begin{aligned}
I_{ri1} &= \sum_k R_{rik}^{(t)} \left\{ -\frac{1}{2} \ln |\Lambda_r| - \frac{1}{2} \left( \mathbf{x}_{ri}^{\mathrm{T}} \Lambda_r^{-1} \mathbf{x}_{ri} + tr(\mathbf{W}^{\mathrm{T}} \Lambda_r^{-1} \mathbf{W} (\langle (\mathbf{z}_{ri} + \mathbf{v}_{ri})(\mathbf{z}_{ri} + \mathbf{v}_{ri})^{\mathrm{T}} \rangle_k^{(t)})) \right.\right. \\
&- \left.\left. 2 \mathbf{x}_{ri}^{\mathrm{T}} \Lambda_r^{-1} \mathbf{W} \langle \mathbf{z}_{ri} + \mathbf{v}_{ri} \rangle_k^{(t)} \right) \right\},
\end{aligned}$$

$$I_{ri2} = \sum_k R_{rik}^{(t)} \left\{ -\frac{1}{2}\ln|\Sigma_k| - \frac{1}{2}tr(\Sigma_k^{-1}\langle \mathbf{z}_{ri}\mathbf{z}_{ri}^{\mathrm{T}}\rangle_k^{(t)}) + (\mu_k)^{\mathrm{T}}\Sigma_k^{-1}\langle \mathbf{z}_{ri}\rangle_k^{(t)} - \frac{1}{2}\mu_k^{\mathrm{T}}\Sigma_k^{-1}\mu_k \right\},$$

$$I_{ri3} = -\frac{1}{2}\sum_{k=1}^{K} R_{rik}^{(t)} \left\{ \ln|\tilde{\Psi}_r| + \mu_{v_{ri}}^{\mathrm{T}}\tilde{\Psi}_r^{-1}\mu_{v_{ri}} + tr(\tilde{\Psi}_r^{-1}\langle \mathbf{v}_{ri}\mathbf{v}_{ri}^{\mathrm{T}}\rangle_k^{(t)}) - 2\mu_{v_{ri}}^{\mathrm{T}}\tilde{\Psi}_r^{-1}\langle \mathbf{v}_{ri}\rangle_k^{(t)} \right\},$$

and

$$I_{ri4} = -\ln C_{ri}(\beta_r, \hat{\mathbf{y}}_{N_{ri}}) - \beta_r \sum_k R_{rik}^{(t)} \sum_{i' \in N_{ri}} \{1 - \delta(k, \hat{y}_{ri'})\}.$$

At this point, we have obtained the explicit Q-function; next, we solve the iterative solution for the model parameters, called the M-step, based on the Q-function.

##### 1.2.3 M-step

Taking derivatives of $Q$ function with respect to each parameter in $\boldsymbol{\theta}$, we obtain the iterative solution of model parameters in the $(t+1)$-th iteration, given by

$$\mu_k = \left\{ \sum_r \sum_{i=1}^{n_r} R_{rik}^{(t)} \right\}^{-1} \left\{ \sum_r \sum_{i=1}^{n_r} R_{rik}^{(t)} \langle \mathbf{z}_{ri}\rangle_k^{(t)} \right\}, \tag{11}$$

$$\Sigma_k = \frac{\sum_r \sum_i R_{rik}^{(t)} \left\{ \langle \mathbf{z}_{ri}\mathbf{z}_{ri}^{\mathrm{T}}\rangle_k^{(t)} - 2\langle \mathbf{z}_{ri}\rangle_k^{(t)}(\mu_k)^{\mathrm{T}} + \mu_k\mu_k^{\mathrm{T}} \right\}}{\sum_r \sum_i R_{rik}(\boldsymbol{\theta}^{(t)})}, \tag{12}$$

$$\tilde{\Psi}_r = \frac{\sum_i \sum_k R_{rik}^{(t)}(\mu_{v_{ri}}\mu_{v_{ri}}^{\mathrm{T}} + \langle \mathbf{v}_{ri}\mathbf{v}_{ri}^{\mathrm{T}}\rangle_k^{(t)} - 2\langle \mathbf{v}_{ri}\rangle_k^{(t)}\mu_{v_{ri}}^{\mathrm{T}})}{n_r}, \tag{13}$$

$$vec(\mathbf{W}^{\mathrm{T}}) = \mathbf{B}_w^{-1} vec(\mathbf{A}_w^{\mathrm{T}}), \tag{14}$$

$$\text{where } \mathbf{A}_w = \sum_r \sum_i \sum_k R_{rik}^{(t)} \Lambda_r^{-1} \mathbf{x}_{ri} \langle \mathbf{z}_{ri} + \mathbf{v}_{ri}\rangle_k^{(t),T},$$

$$\mathbf{B}_w = \sum_r \sum_i \sum_k R_{rik}^{(t)} \Lambda_r^{-1} \otimes \langle (\mathbf{z}_{ri} + \mathbf{v}_{ri})(\mathbf{z}_{ri} + \mathbf{v}_{ri})^{\mathrm{T}} \rangle_k^{(t)},$$

$$\lambda_{rj} = \frac{1}{n_r} \sum_i \sum_k R_{rik}^{(t)} \left( s_{1rijk}^2 + s_{2rjk}^2 \right), \tag{15}$$

$$\beta_r = \arg\max_{\beta_r} \sum_i I_{ri4}, \tag{16}$$

where the $vec$ operator is an operator that transforms a matrix into a column vector by vertically stacking the columns of the matrix, $s_{1rijk}^2 = (x_{rij} - \mathbf{w}_j^{\mathrm{T}} \langle \mathbf{z}_{ri} + \mathbf{v}_{ri}\rangle_k^{(t)})^2$, $s_{2rjk}^2 = \mathbf{w}_j^{\mathrm{T}} \Sigma_{zv,rk}^{(t)} \mathbf{w}_j$ and $\mathbf{w}_j$ is the $j$-th column of $\mathbf{W}^{\mathrm{T}}$.

To extract batch-corrected and cell-type/domain-relevant low dimensional representations of $\mathbf{X}_r$ for each sample, we estimate $\mathbf{z}_{ri}$ by the posterior expectation

$$E(\mathbf{z}_{ri}|\mathbf{x}_{ri}) = \sum_{k=1}^{K} R_{rik} \mathrm{E}_k(\mathbf{z}_{ri}|\mathbf{x}_{ri}), \tag{17}$$

since $E(\mathbf{z}_{ri}|\mathbf{x}_{ri}) = \int \mathbf{z}_{ri} P(\mathbf{z}_{ri}|\mathbf{x}_{ri}; \boldsymbol{\theta}) d\mathbf{z}_{ri}$ with $P(\mathbf{z}_{ri}|\mathbf{x}_{ri}; \boldsymbol{\theta}) = \sum_{k=1}^{K} P(y_{ri} = k|\mathbf{x}_{ri}) P(\mathbf{z}_{ri}|y_{ri} = k, \mathbf{x}_{ri}) = \sum_{k=1}^{K} R_{ik} P(\mathbf{z}_{ri}|y_{ri} = k, \mathbf{x}_{ri})$.

#### 1.3 Computational challenge

Since the derivative of $Q$ function with respect to $\beta_r$ is complicated due to the complexity of the partition function $C_{ri}(\beta_r, \hat{\mathbf{y}}_{N_{ri}})$, we use a grid search strategy to update $\beta_r$ using the fact $\beta_r = \arg\max_{\beta_r} \sum_i I_{ri4}$. Given a grid of $\beta_r$, such as $\{\beta_{r1}, \cdots, \beta_{rS}\}$, we evaluate the value of $\sum_i I_{ri4}$ for each $\beta_{rs}, s \leq S$, then we choose the value that maximizes $\sum_i I_{ri4}$.

To update $\mathbf{W}$, we are required to evaluate a $pq \times pq$ matrix inverse $\mathbf{B}_w^{-1}$, requiring the computational complexity $O(q^3 p^3)$ in each iteration. However, it is not feasible to compute this directly, as the dimension of genes $p$ is very large. Therefore, we derive a simple expression by the fact that $\mathbf{B}_w$ is a block diagonal matrix, i.e.,

$$\begin{aligned}
\mathbf{B}_w &= \sum_r \sum_i \sum_k R_{rik}^{(t)} \Lambda_r^{-1} \otimes \langle \mathbf{u}_{ri} \mathbf{u}_{ri}^{\mathrm{T}} \rangle_k^{(t)} \\
&= \begin{pmatrix} \sum_r \sum_i \sum_k R_{rik}^{(t)} \lambda_{r1}^{-1} \langle \mathbf{u}_{ri} \mathbf{u}_{ri}^{\mathrm{T}} \rangle_k^{(t)} & \dots & \mathbf{0} \\ \dots & \dots & \dots \\ \mathbf{0} & \dots & \sum_r \sum_i \sum_k R_{rik}^{(t)} \lambda_{rp}^{-1} \langle \mathbf{u}_{ri} \mathbf{u}_{ri}^{\mathrm{T}} \rangle_k^{(t)} \end{pmatrix} \\
&= \mathrm{diag}(\mathbf{B}_{w1}, \cdots, \mathbf{B}_{wp}), \mathbf{B}_{wj} \in \mathbb{R}^{q \times q}, j \leq p,
\end{aligned}$$

where $\mathbf{u}_{ri} = \mathbf{z}_{ri} + \mathbf{v}_{ri}$. Thus, using the inverse formula of the block diagonal matrix, we have

$$\mathbf{B}_w^{-1} = \mathrm{diag}(\mathbf{B}_{w1}^{-1}, \cdots, \mathbf{B}_{wp}^{-1}).$$

Denote $\mathbf{A}_w = (\mathbf{a}_{1,w}, \cdots, \mathbf{a}_{p,w})^{\mathrm{T}}$, where $\mathbf{a}_{j,w} = \sum_r \sum_i \sum_k R_{rik}^{(t)} \lambda_{rj}^{-1} \mathbf{x}_{rij} \langle \mathbf{z}_{ri} + \mathbf{v}_{ri} \rangle_k^{(t),\mathrm{T}}$, then we have

$$\mathbf{w}_j = \mathbf{B}_{wj}^{-1} \mathbf{a}_{j,w},$$

where $\mathbf{B}_{wj} = \sum_r \lambda_{rj}^{-1} \left\{ \sum_i \sum_k R_{rik}^{(t)} \langle (\mathbf{z}_{ri} + \mathbf{v}_{ri})(\mathbf{z}_{ri} + \mathbf{v}_{ri})^{\mathrm{T}} \rangle_k^{(t)} \right\}$. Thus, we reduce the computational complexity from $O(q^3 p^3)$ to $O(q^3 p)$, which makes the ICM-EM of PRECAST computationally efficient.

#### 1.4 Algorithm implementation of PRECAST

We summarize the proposed ICM-EM algorithm as algorithms 1 and 2. Algorithm 1 describes the detailed implementation of ICM algorithm used in the algorithm 2 for predicting $\mathbf{y}$ and $\mathbf{V}$. Algorithm 2 presents details of the ICM-EM algorithm. Following the suggestion of Besag [2], we set the initial value of $\beta_r$ to 1.5. To get the initial values of other parameters, we perform PCA analysis on $\mathbf{X}$ with the number of PCs $q$, then obtain loading matrix $L_1$ and score matrix $L_2$, and perform Gaussian mixture model estimation on the score matrix $L_2$, then obtain mean component $\tilde{\mu}_k$, equal covariance component $\tilde{\Sigma}_k$ and cluster labels $\tilde{\mathbf{y}}$. Finally, we initialize $\hat{\mathbf{y}}^{(0)} = \tilde{\mathbf{y}}, \widehat{\mathbf{V}}^{(0)} = \mathbf{0}, \mathbf{W}^{(0)} = L_1, \mu_k^{(0)} = \tilde{\mu}_k, k \leq K, \Sigma_k^{(0)} = \tilde{\Sigma}_k, \tilde{\Psi}_r^{(0)} = cov(L_2)$ and $\lambda_{rj}^{(0)} = \frac{1}{n_r} \sum_{i=1}^{n_r} (x_{rij} - L_{2,ri.} L_{1,j.}^{\mathrm{T}})^2, r \leq M$, where $L_{2,ri.}$ is the row of $L_2$ corresponding to $i$-th spot in sample $r$ and $L_{1,j.}$ is the $j$-th row of $L_1$.

#### 1.5 Determining the number of clusters

Modified BIC criteria [3, 4] were used to determine the number of clusters $K$,

$$MBIC(K) = -2 \ln P(\mathbf{X}; \widehat{\boldsymbol{\theta}}(K)) + C_n df(K) \ln n, \tag{18}$$

**Algorithm 1** ICM algorithm used in ICM-EM algorithm

**Input:** $\mathbf{X}$, $\hat{\mathbf{y}}^{(t-1)}, \boldsymbol{\theta}^{(t-1)}$, $\mathcal{S} = \{s_{ri}\}$, maximum iterations of ICM $maxIter\_ICM$, relative tolerance of difference of total objective function $eps\_ICM$.

**Output:** $\hat{\mathbf{y}}^{(t)}, \widehat{\mathbf{V}}^{(t)}$.

**for** each $r \in 1, \cdots, M$ **do**
  **for** each $l \in 1, \cdots, maxIter\_ICM$ **do**
    Update $\mathbf{y}_r^{(t-1),l}$ based on equation (6);
    Update $\mathbf{V}_r^{(t-1),l}$ based on equation (7);
    Evaluate the total objective function, $\Xi_r(l)$, by (8).
    **if** $|\Xi_r(l) - \Xi_r(l-1)|/|\Xi_r(l-1)| < eps\_ICM$ **then**
      break;
    **end if**
  **end for**
**end for**
$\hat{\mathbf{y}}^{(t)} = \hat{\mathbf{y}}^{(t-1),l}, \widehat{V}^{(t)} = \widehat{\mathbf{V}}^{(t-1),l}$
**return** $\hat{\mathbf{y}}^{(t)}, \widehat{\mathbf{V}}^{(t)}$

**Algorithm 2** The proposed ICM-EM algorithm for PRECAST

**Input:** $\mathbf{X}_r$, $\mathcal{S} = \{s_{ri}\}, r \leq M, i \leq n_r$, $q$, $K$, grid points of $\beta_r$, $beta\_grid$, maximum iterations of EM $maxIter$, relative tolerance of pseudo loglikelihood $epsLogLike$, maximum iterations of ICM $maxIter\_ICM$, relative tolerance of total objective function in ICM $eps\_ICM$.

**Output:** $\hat{\mathbf{y}}, \widehat{\mathbf{V}}, \widehat{\mathbf{Z}}$ and $\widehat{\boldsymbol{\theta}}$

Initialize $\hat{\mathbf{y}}^{(0)}, \widehat{\mathbf{V}}^{(0)}$ and $\boldsymbol{\theta}^{(0)}$
**for** each $t \in 1, \cdots, maxIter$ **do**
  Update $(\hat{\mathbf{y}}^{(t)}, \widehat{\mathbf{V}}^{(t)})$ based on function $ICM(X, \hat{\mathbf{y}}^{(t-1)}, \widehat{\mathbf{V}}^{(t-1)}, \boldsymbol{\theta}^{(t-1)}, \mathcal{S}, maxIter\_ICM, eps\_ICM)$;
  Update $\{\beta_r^{(t)}, r \leq M\}$ based on grid search on $beta\_grid$;
  Update $\{\mu_k^{(t)}, k \leq K\}$ based on Equation (11);
  Update $\{\Sigma_k^{(t)}, k \leq K\}$ based on Equation (12);
  Update $\{\tilde{\Psi}_r^{(t)}, r \leq M\}$ based on Equation (13);
  Update $\mathbf{W}^{(t)}$ based on Equation (14);
  Update $\{\Lambda_r^{(t)}, r \leq M\}$ based on Equation (15);
  Evaluate the pseudo observational loglikelihood, $LogLike(t)$, by (5).
  **if** $|LogLike(t) - LogLike(t-1)|/|LogLike(t-1)| < epsLogLike$ **then**
    break;
  **end if**
**end for**
Evaluate $\widehat{\mathbf{Z}}$ based on Equation (17) by replacing $R_{rik}$ and $\langle \mathbf{z}_{ri} \rangle_k$ with $R_{rik}^{(t)}$ and $\langle \mathbf{z}_{ri} \rangle_k^{(t)}$, respectively.
**return** $\hat{\mathbf{y}}, \widehat{\mathbf{V}}, \widehat{\mathbf{Z}}$ and $\widehat{\boldsymbol{\theta}}$

where $df(K) = 2qK + pq + M(p + \frac{q(q+1)}{2} + 1)$ and $C_n$ is a positive constant that can depend on $n$ and $p$. When $C_n = 1$, the modified BIC reduces to the traditional BIC [5]. Following the lead of Ma et. al [4], we take the same strategy and let $C_n = c\ln(\ln(p+n))$, where $c$ is the positive constant default 1. This modified BIC is proposed for high-dimensional data settings to select tuning parameters, which is more suitable than conventional BIC criteria for our proposed model involving high-dimensional gene expressions. Since the observed log-likelihood $\ln P(\mathbf{X}; \widehat{\boldsymbol{\theta}}(K))$ is intractable in Enq. (18), it is approximated by the pseudo observed log-likelihood $\ln \tilde{P}(\mathbf{X}; \widehat{\boldsymbol{\theta}}(K))$ in Enq. (5).

#### 1.6 Definition of neighbors

We briefly introduce the precise definition of the neighborhood system in this section. We set the number of neighbors for each spot to $m$, i.e., $m_{ri} = m$. We define the neighbors of a spot using the Euclidean distance between spatial coordinates for two different spots. For spot $s_{ri}$, the $m$ nearest spots, in the sense of Euclidean distance between them and $s_{ri}$, are defined as the neighbors of spot $s_{ri}$. In this paper, we consider $m = 4$ for data from ST platform due to the rectangle lattice structure, and $m = 6$ for data from 10X Visium platform due to the hexagonal lattice structure. Because the low-resolution data obtained by Slide-seqV2 platform has close resolution with the data from 10X Visium, we also use $m = 6$ for the resolution-reduced data. In contrast, we use $m = 24$ for the raw-resolution data measured on the Slide-seqV2 platform, whose resolution is four-fold that of 10X Visium.

#### 1.7 Details of compared methods

We provided more details about the compared integration methods. Seurat V3 method is based on finding the mutual nearest neighbors. First, anchor pairs between different batches were obtained using the function `FindIntegrationAnchors` in the *Seurat* R package. Then, the `IntegrateData` function in *Seurat* was used to integrate the datasets based on the anchor pairs and their confidence scores. Finally, we obtained the aligned 15-dimensional embeddings using the function `RunPCA` in *Seurat*. Compared with the original MNN method, fastMNN is a faster version based on PCA and is used to find the mutual nearest neighbors, greatly increasing the computational efficiency. We used `fastMNN` in the R package *batchelor* with the top 15 PCs and other default parameters and obtained all aligned 15-dimensional embeddings. Another PCA-based method, Harmony corrects batch-contaminated PCs by iteratively performing maximum diversity clustering and linear mixture model correction. For the analysis, we first obtained the 15-dimensional PCs by performing PCA on the combined normalized expression matrix, then took the obtained PCs and batch information as input for the function `HarmonyMatrix` in the R package *harmony*, and lastly, obtained the aligned 15-dimensional embeddings as output. Scanorama is a generalization of mutual nearest-neighbor matching used to find similar elements among many datasets. Instead of searching for similar elements in high-dimensional gene space, Scanorama applies randomized singular value decomposition to extract the aligned embeddings from the combined cell-by-gene expression matrix. We conducted the analysis following the author's pipeline `https://scanpy-tutorials.readthedocs.io/en/latest/spatial/integration-scanorama.html` and set the parameter *dimred=15* in function `correct_scanpy` to extract aligned 15-dimensional embeddings. The pipelines for scGen and scVI, which are two deep-learning-based methods, were followed using the two respective links, `https://scgen.readthedocs.io/en/latest/tutorials/scgen_batch_removal.html` and `https://docs.scvi-tools.org/en/stable/tutorials/notebooks/tabula_muris.html`, and we set the number of latent embeddings to 15 for both methods in the analysis. MEFISTO is a factor-model-based integration method that integrates multiple datasets by simultaneously identifying and aligning the underlying patterns of variation. To implement this, we followed the author's pipeline provided at `https://github.com/bioFAM/MEFISTO_tutorials/blob/master/MEFISTO_microbiome.ipynb` and `https://github.com/bioFAM/MEFISTO_tutorials/blob/master/MEFISTO_ST.ipynb`, and set the number of factors to 15. PASTE is a pairwise alignment method that integrates adjacent slices based on transcriptional and spatial similarity using optimal transport and nonnegative matrix factorization; we followed the author's pipeline `https://github.com/raphael-group/paste/blob/main/docs/source/notebooks/getting-started.ipynb` for implementation and set the number of low-dimensional representation of center slice to 15.

More details were also presented for the compared clustering methods. SC-MEB and BayesSpace were recently developed to perform spatial clustering based on a discrete Markov random field [6, 7], Louvain is a conventional non-spatial clustering algorithm based on community detection in large networks [8], and BASS is a newly developed clustering method for multiple SRT data based on the aligned embeddings from Harmony. The aligned embeddings and spatial coordinates were used as input for SC-MEB and BayesSpace, only the aligned embeddings were the input for Louvain, and the count matrices and spatial coordinates were the input for BASS. Similar to PRECAST, SC-MEB uses MBIC [3] to determine the number of clusters in a data-driven manner, BayesSpace adopts the average loglikelihood-maximization-based method in early iterations, Louvain uses a community-modularity maximizing rule [8], while BASS requires users to specify the number of clusters [9]. For fair comparison, BASS uses the number of clusters selected by PRECAST.

#### 1.8 Details of evaluation metrics

In the simulation, the F1 score of average silhouette coefficients and cLISI/iLISI were used to assess performance in batch correction. ARI and NMI were adopted to measure the similarity between the estimated clusters and the true one. CCor was used to measure the similarity of the batch corrected low-dimensional embeddings and the true one. Concor was used to measure remained information between gene expressions and spatial domains by excluding the effect of the extracted latent features. In the real data analysis, manual annotations based on additional experiments were available. ARI and NMI were used to measure the similarity between labels from the estimated partition and the manually annotated clusters, and the F1 score of average silhouette coefficients and cLISI/iLISI were also evaluated based on the manual annotations.

**Local inverse Simpson's index**. To assess the performance of the batch correction, cell-type/integration local inverse Simpson's index (cLISI/iLISI) [10] was used to evaluate the quality of merging the shared cell populations among batches and mixing spots from $M$ tissue slides, respectively. Korsunsky et al. [10] pointed out that LISI is more sensitive to local distances and more interpretable than conventional metrics such as entropy score [11] and kBET [12]. LISI assigns each spot a diversity score which is the effective number of cell types/data batches in that cell's/batch's neighborhood. For $M$ SRT batches with $K$ cell types, accurate integration should maintain a cLISI value close to 1, reflecting the purity of unique cell type in the neighbors of each spot defined by the low-dimensional embeddings, and an iLISI value close to $M$, means the sufficient mixing of $M$ data batches. An erroneous embedding would include neighborhoods with a cLISI more than 1 and an iLISI less than $M$, where an extremely worse case is cLISI close to $K$ and iLISI close to 1, indicating that neighbors have $K$ different types of cells but one data batch.

**Silhouette coefficient**. To simultaneously evaluate the separation of each cell/domain cluster and mixing of multiple datasets, we also consider a metric called the F1 score of average silhouette coefficients based on two groupings. Specifically, we calculated the silhouette coefficient [11] with clusters defined by cell types/domains and batches, respectively. For a spot $i$, let $a(i)$ be the average distance between $i$ and all the other spots within the same cluster, and denote $b(i)$ to be the smallest average distance between spot $i$ and all the spots in any other cluster, then the silhouette coefficient of spot $i$ is defined as

$$\mathrm{S}(i) = \frac{a(i) - b(i)}{\max\{a(i), b(i)\}} \in [-1, 1].$$

After batch effect removal, the average silhouette coefficient of all spots from different batches is calculated by

$$\mathrm{silh} = \frac{1}{\sum_{r=1}^{M} n_r} \sum_{r=1}^{M} \sum_{i \in \mathcal{S}_r} \mathrm{S}(i),$$

where $\mathcal{S}_r$ represents the spot index of sample $r$. We evaluated the average silhouette coefficient of SRT expression data using two different clusters: (i) clusters defined by known cell types/domains as the cell type/domain silhouette coefficient ($\mathrm{silh}_{cluster}$); and (2) clusters defined by batches as the batch silhouette coefficient ($\mathrm{silh}_{batch}$). Ideally, the aligned low-dimensional embeddings matrix has a large $\mathrm{silh}_{cluster}$, which indicates the preservation of biological signals, and a small $\mathrm{silh}_{batch}$, which suggests the spots are not grouped by batch. To jointly consider these two evaluation measurements, we calculated the harmonic mean of these two average silhouette coefficients following transformation, which is called the F1 score, ranges from 0 to 1, and is given by

$$\mathrm{F1\_score\_silh} = \frac{2(1 - \mathrm{silh}'_{\mathrm{batch}})\mathrm{silh}'_{cluster}}{\mathrm{silh}'_{cluster} + (1 - \mathrm{silh}'_{\mathrm{batch}})} \in [0, 1],$$

where $\mathrm{silh}'_{\mathrm{batch}} = \frac{1+\mathrm{silh}_{\mathrm{batch}}}{2}$ and $\mathrm{silh}'_{\mathrm{cluster}} = \frac{1+\mathrm{silh}_{\mathrm{cluster}}}{2}$. A larger value of F1_score_silh indicates that the cell type/domain assignment in the aligned dataset is more appropriate, where a spot is close to spots of the same type and distant from spots of different types. $\mathrm{S}(i)$ is calculated using the Euclidean distance on the batch-corrected low-dimensional embeddings.

**Canonical correlation coefficients & conditional correlation**. For dimension reduction, we consider two measurements to assess the performance of the recovery of true latent features. The first one is the mean canonical correlation between the estimated features and the true one defined as

$$\mathrm{CCor} = \frac{1}{q} \sum_{l=1}^{q} \zeta_l(\mathbf{z}_i, \hat{\mathbf{z}}_i),$$

where $\zeta_l$ is the $l$-th canonical correlation coefficients. The mean canonical correlation coefficient measures the similarity of two sets of random variables, and a larger value means a better estimation of true latent features. The second one is the mean conditional correlation between gene expression $\mathbf{x}_i$ and cell-type/domain label $y_i$ given the estimated latent features $\hat{\mathbf{z}}_i$ defined as

$$\text{ConCor} = \frac{1}{p}\sum_{j=1}^{p} corr(y_i, resid_{ij}),$$

where $resid_i$ is the residual of $x_{ij}$ regressing on $\hat{\mathbf{z}}_i$ and $corr(y_i, resid_{ij})$ is the Pearson correlation coefficient between $y_i$ and $resid_{ij}$. Ideally, we want to obtain the estimated aligned low-dimensional features that contain all information on cell types/domains, in other words, $y_i \perp \mathbf{x}_i | \hat{\mathbf{z}}_i$. Thus, a smaller conditional correlation suggests a better performance. In the simulation studies, the true latent features are known, thus, both CCor and ConCor are evaluated. In the real data analysis, only ConCor is evaluated since true latent features are infeasible.

**Adjusted Rand index**. For evaluating clustering performance, we consider the adjusted Rand index (ARI) [13] and normalized mutual information (NMI) [14]. ARI [13] is the corrected version of the Rand index (RI) [15] that avoids some drawbacks of RI [13] and is defined as

$$\text{ARI} = \frac{RI - E(RI)}{\max(RI) - E(RI)},$$

where $E(RI)$ and $\max(RI)$ are the expected value and maximum value of $RI$, respectively. Suppose there are $n$ spots in SRT dataset. Let $U = (u_1, \cdots, u_i, \cdots, u_K)$ and $\bar{U} = (\bar{u}_1, \cdots, \bar{u}_j, \cdots, \bar{u}_L)$ denote two clustering labels for $n$ spots of all combined samples from two different clustering methods, where $K$ and $L$ corresponds to the numbers of clusters from these two methods. Let $n_{ij}$ be the number of spots belonging to both classes $u_i$ and $\bar{u}_j$, and $a_i$ and $b_j$ be the number of spots in classes $u_i$ and $\bar{u}_j$, respectively; then the specific formula of ARI is given by

$$\text{ARI} = \frac{\sum_{ij}\binom{n_{ij}}{2} - [\sum_i \binom{a_i}{2}\sum_j \binom{b_j}{2}]/\binom{n}{2}}{\frac{1}{2}[\sum_i \binom{a_i}{2} + \sum_j \binom{b_j}{2}] - [\sum_i \binom{a_i}{2}\sum_j \binom{b_j}{2}]/\binom{n}{2}}.$$

ARI measures the similarity between two different partitions and ranges from $-1$ to 1. A larger value of ARI means a higher similarity between two partitions. ARI takes a value of 1 when the two partitions are equal up to a permutation.

**Normalized mutual information**. NMI is a revised version of mutual information (MI) that makes the value of MI range from zero to one. MI originates from probability theory and information theory, and measures the mutual dependence of two random variables. More specifically, it quantifies the "amount of information" in units such as Shannons (bits) obtained for one random variable by observing the other random variable. Let $x$ and $y$ be two discrete random variables, i.e. the random variables taking the values of class labels on two different partitions, then their MI can be defined as

$$\text{MI}(x, y) = \sum_x \sum_y P(x, y) \ln \frac{P(x, y)}{P(x)P(y)} = H(x) + H(y) - H(x, y), \tag{19}$$

where $P(x, y)$ is the joint distribution of $(x, y)$, $P(x)$ and $P(y)$ are the marginal distributions of $x$ and $y$, respectively, and $H(x)$, $H(y)$ and $H(x, y)$ are the marginal entropies of $x$, $y$ and the joint entropy of $(x, y)$, respectively. Intuitively, mutual information measures the information that $x$ and $y$ share. If $x$ and $y$ do not share information and are mutually independent, then $\text{MI}(x, y) = 0$. At the other extreme, if $y = x$, then $\text{MI}(x, y) = H(x)$. This indicates MI does not take values between zero and one. Therefore, some normalized versions have been proposed and we used one version of them, defined as

$$\text{NMI}(x, y) = \frac{\text{MI}(x, y)}{\max(H(x), H(y))}.$$

From the above formula, we know when the two partitions are equal up to a permutation, the NMI takes a value of 1.

#### 1.9 Additional simulations

**Scenario 4. Normalized gene expression data: domain labels and spatial coordinates from Potts models.** For this scenario, we generated log-normalized gene expression data for three samples and the spatial coordinates based on Potts models with four neighborhoods. The class label $y_{ri}$, loading matrices $\mathbf{W}, \mathbf{W}_r$, latent features $\boldsymbol{\nu}_{ri}$ and $\mathbf{z}_{ri}$ were generated in the same way as described for scenario 1, except that $\mu_k$ had a different value (Supplementary Data 8), and we only generated normalized gene expression $\mathbf{x}_{ri}$ using $\mathbf{x}_{ri} = \mathbf{W}(\mathbf{z}_{ri} + \boldsymbol{\nu}_{ri}) + \mathbf{W}_r\boldsymbol{\zeta}_{ri} + \boldsymbol{\varepsilon}_{ri}$ and $\boldsymbol{\varepsilon}_{ri} \sim N(\mathbf{0}, \Lambda_r)$, where $\Lambda_r = \text{diag}(\lambda_{rj}), j = 1, \ldots, p$, $\lambda_{1j} = 2(1 + 0.5|z_{1j}|)$ with $z_{1j} \overset{i.i.d.}{\sim} N(0, 3)$; $\lambda_{2j} = 2(1 + 0.2z_{2j})$ with $z_{2j} \overset{i.i.d.}{\sim} U[0, 1]$; and $\lambda_{3j} = 2(1 + z_{3j})$ with $z_{3j} \overset{i.i.d.}{\sim} U[0, 1]$.

**Scenario 5. Normalized gene expression data: domain labels and spatial coordinates from DLPFC data.** In this scenario, we generated log-normalized gene expression data based on three DLFPC datasets (ID: 151507, 151669 and 151673) from three donors (Visium platform). The class label $y_{ri}$, loading matrices $\mathbf{W}, \mathbf{W}_r$, latent features $\boldsymbol{\nu}_{ri}$ and $\mathbf{z}_{ri}$ were generated as described for scenario 2. The only difference was that normalized gene expression data $\mathbf{x}_{ri}$ was generated using $\mathbf{x}_{ri} = \mathbf{W}(\mathbf{z}_{ri} + \boldsymbol{\nu}_{ri}) + \mathbf{W}_r\boldsymbol{\zeta}_{ri} + \boldsymbol{\varepsilon}_{ri}, \tau_{rj} \sim N(0, 4)$ and $\boldsymbol{\varepsilon}_{ri} \sim N(\mathbf{0}, \Lambda_r)$, where $\Lambda_r = \text{diag}(\lambda_{rj}), j = 1, \ldots, p$, $\lambda_{1j} = 2(1 + 0.5|z_{1j}|)$ with $z_{1j} \overset{i.i.d.}{\sim} N(0, 3)$; $\lambda_{2j} = 2(1 + 0.2z_{2j})$ with $z_{2j} \overset{i.i.d.}{\sim} U[0, 1]$; and $\lambda_{3j} = 2(1 + z_{3j})$ with $z_{3j} \overset{i.i.d.}{\sim} U[0, 1]$.

### Supplementary Figures

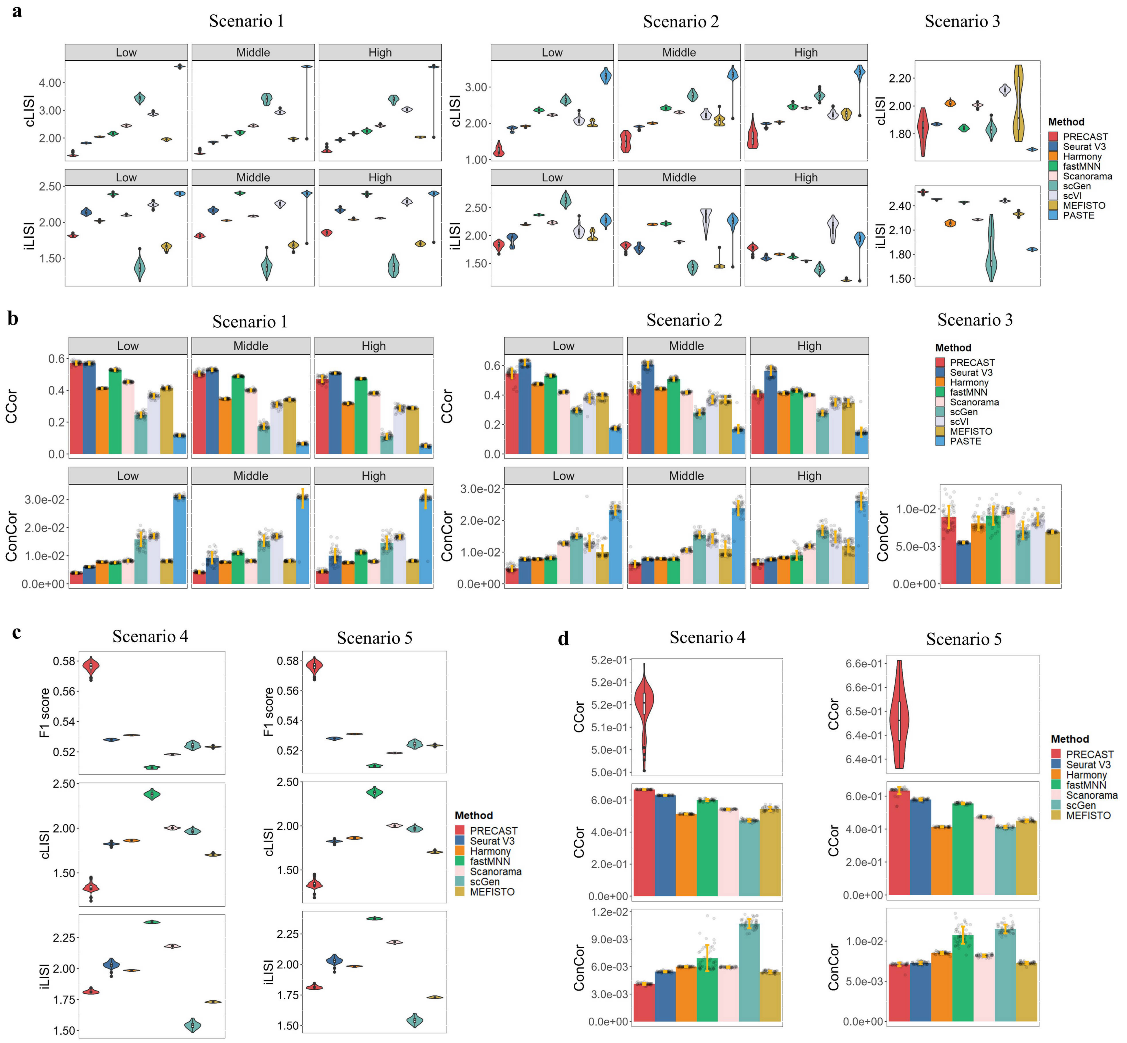


Figure S1: Batch correction and dimension reduction performance for simulated data ($n = 11,425$ spots over 50 independent experiments). a. Violin plot of cLISI/iLISI based on the batch-corrected 15-dimensional embeddings from PRECAST and eight other compared methods in scenarios 1-3. b. Bar plot of canonical/conditional correlations based on the batch-corrected 15-dimensional embeddings from PRECAST and eight other methods in scenarios 1-3. In scenario 3, the true low-dimensional embeddings are unknown, so we could not evaluate the canonical correlation (CCor). c. Violin plot of cLISI/iLISI/F1 score based on the batch-corrected 15-dimensional embeddings from PRECAST and five other compared methods in scenarios 4 and 5. d. Violin plot showing the canonical correlations between estimated slide-specific embeddings due to neighboring microenvironments from PRECAST and the underly truth; Bar plot of canonical/conditional correlations based on the batch-corrected 15-dimensional embeddings from PRECAST and five other methods in scenarios 4 and 5. Note that scVI and PASTE are only applicable to the scenarios 1-3 with count matrices. In the bar plot, the error bands represent the mean value ± standard deviation. In the boxplot, the center line, box lines and whiskers represent the median, upper, and lower quartiles, and 1.5 times interquartile range, respectively.

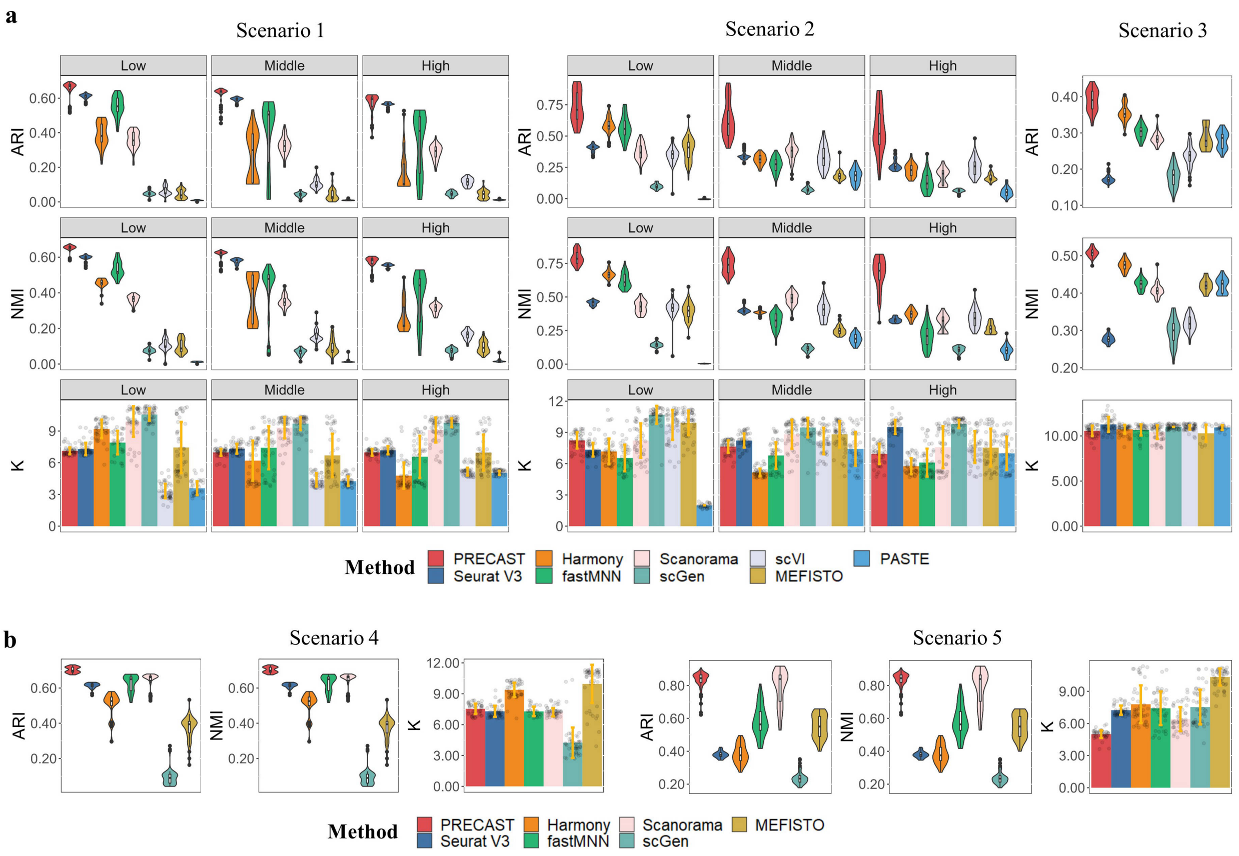


Figure S2: Clustering analysis of simulated data from 50 repeatitions. a. Domain clustering performance of PRECAST and eight other integration methods based on SC-MEB clustering in scenarios 1-3. Upper panel: Violin plot of ARIs from SC-MEB clustering based on the low-dimensional embeddings of Harmony, fastMNN, Scanorama, scGen, scVI, MEFISTO and PASTE. Middle panel: Violin plot of NMIs from PRECAST and other compared methods. Bottom panel: Bar plot showing the number of clusters selected by PRECAST and other compared methods, where the true number of clusters is $K = 7$. b. Domain clustering performance of PRECAST and eight other integration methods based on SC-MEB clustering in scenarios 4 and 5. scVI and PASTE are only applicable to the scenarios 1-3 with count matrices. In the bar plot, the error bands represent the mean value $\pm$ standard deviation. In the boxplot, the center line, box lines and whiskers represent the median, upper, and lower quartiles, and 1.5 times interquartile range, respectively.

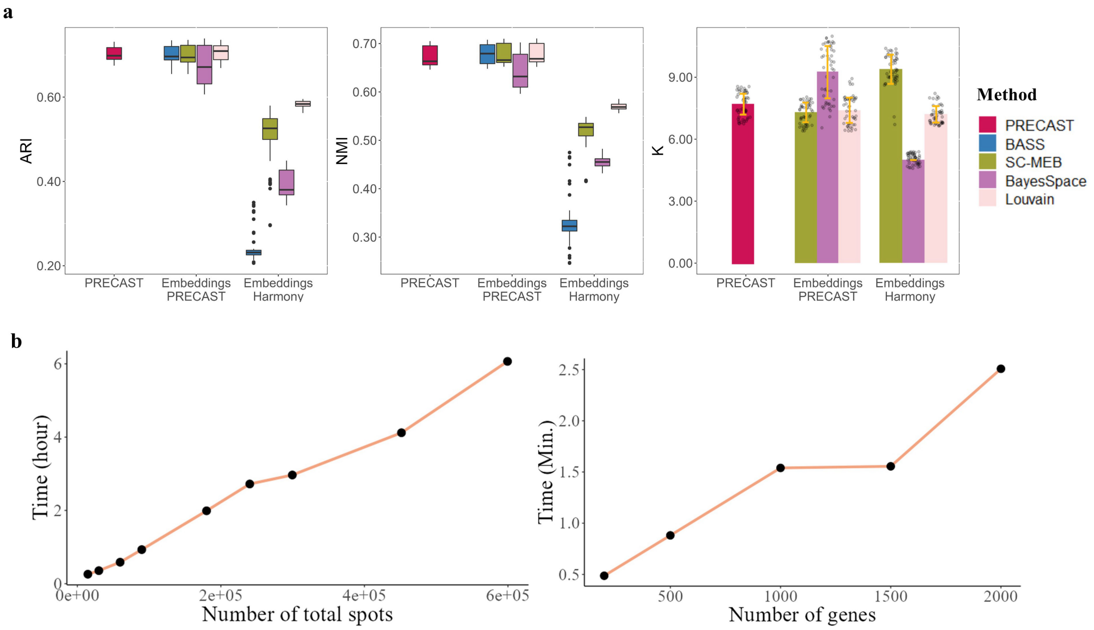


Figure S3: Clustering analysis based on embeddings from PRECAST and Harmony in scenario 4 and scalability analysis ($n = 11,425$ spots over 50 independent experiments). a. Left panel: Box plot of ARIs from PRECAST, and BASS, SC-MEB, BayesSpace and Louvain based on the low-dimensional embeddings of PRECAST and Harmony. Middle panel: Box plot of NMIs from these methods. In the boxplot, the center line, box lines and whiskers represent the median, upper, and lower quartiles, and 1.5 times interquartile range, respectively. Right panel: Bar plot of the number of clusters selected by PRECAST, SC-MEB, BayesSpace and Louvain. In the bar plot, the error bands represent the mean value ± standard deviation. Note that BASS cannot choose the number of clusters automatically, so we used the number of clusters selected by PRECAST. b. Linear computational complexity of PRECAST with regard to the number of spots/genes. Left panel: Line plot of running time and number of spots (given 2000 genes) when running 30 iterations of three datasets on a linux server with 2.10GHz Intel(R) Xeon(R) Gold 6230 CPU and 50G memory. Right panel: Line plot of running time and number of genes (given 15,000 spots) in total when running 30 iterations of three datasets on the same machine.

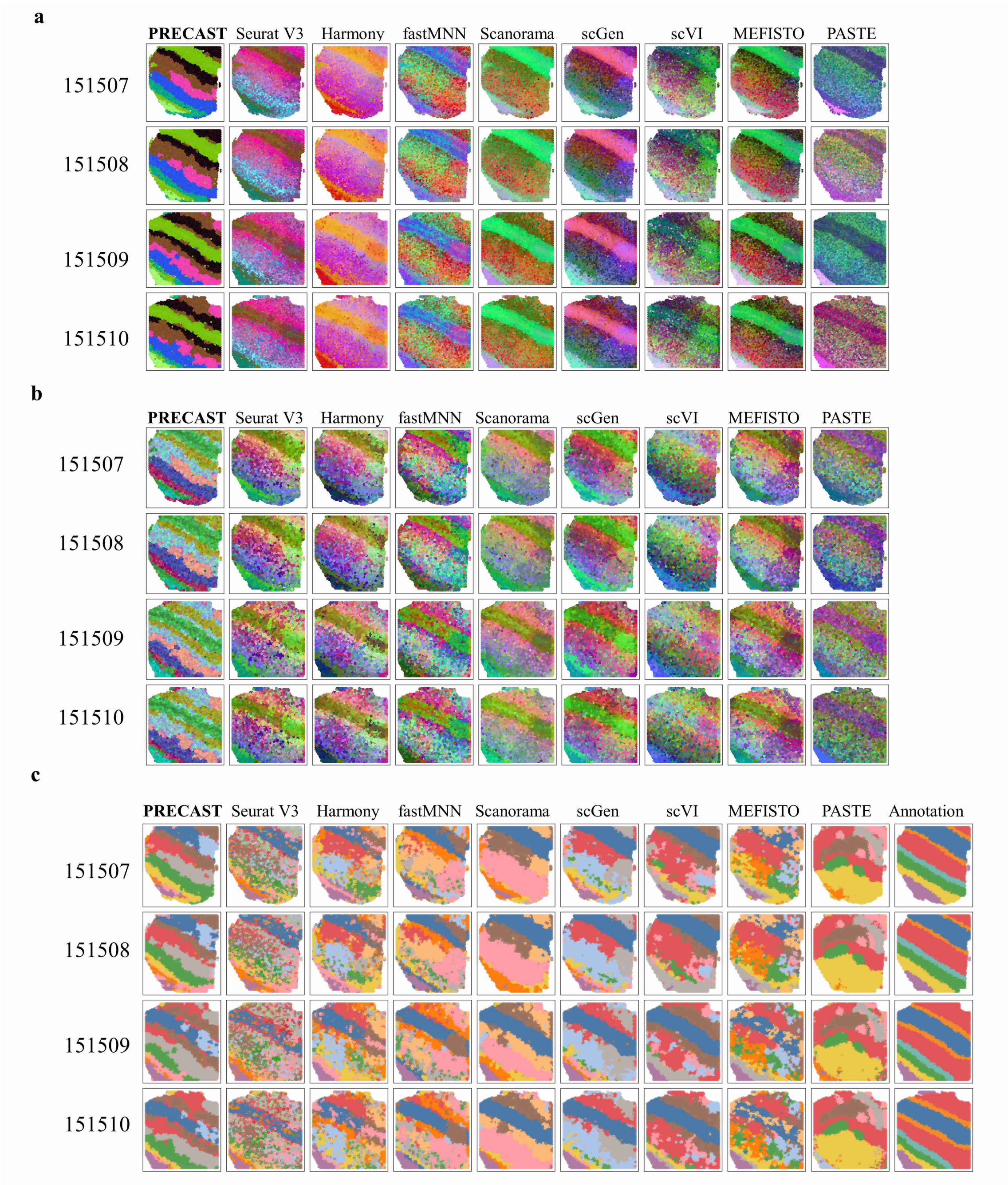


Figure S4: Dimension reduction with batch correction and clustering analysis for the 12 dorsolateral prefrontal cortex Visium sections. a. UMAP RGB plots for samples 1-4 with sample ID 151507-151510 of Donor 1 based on the low-dimensional embeddings from PRECAST and eight other methods. b. tSNE RGB plots for these methods. c. Domain clustering plots for these methods and manual annotations, where SC-MEB is used in the other methods for clustering based on the low-dimensional embeddings.

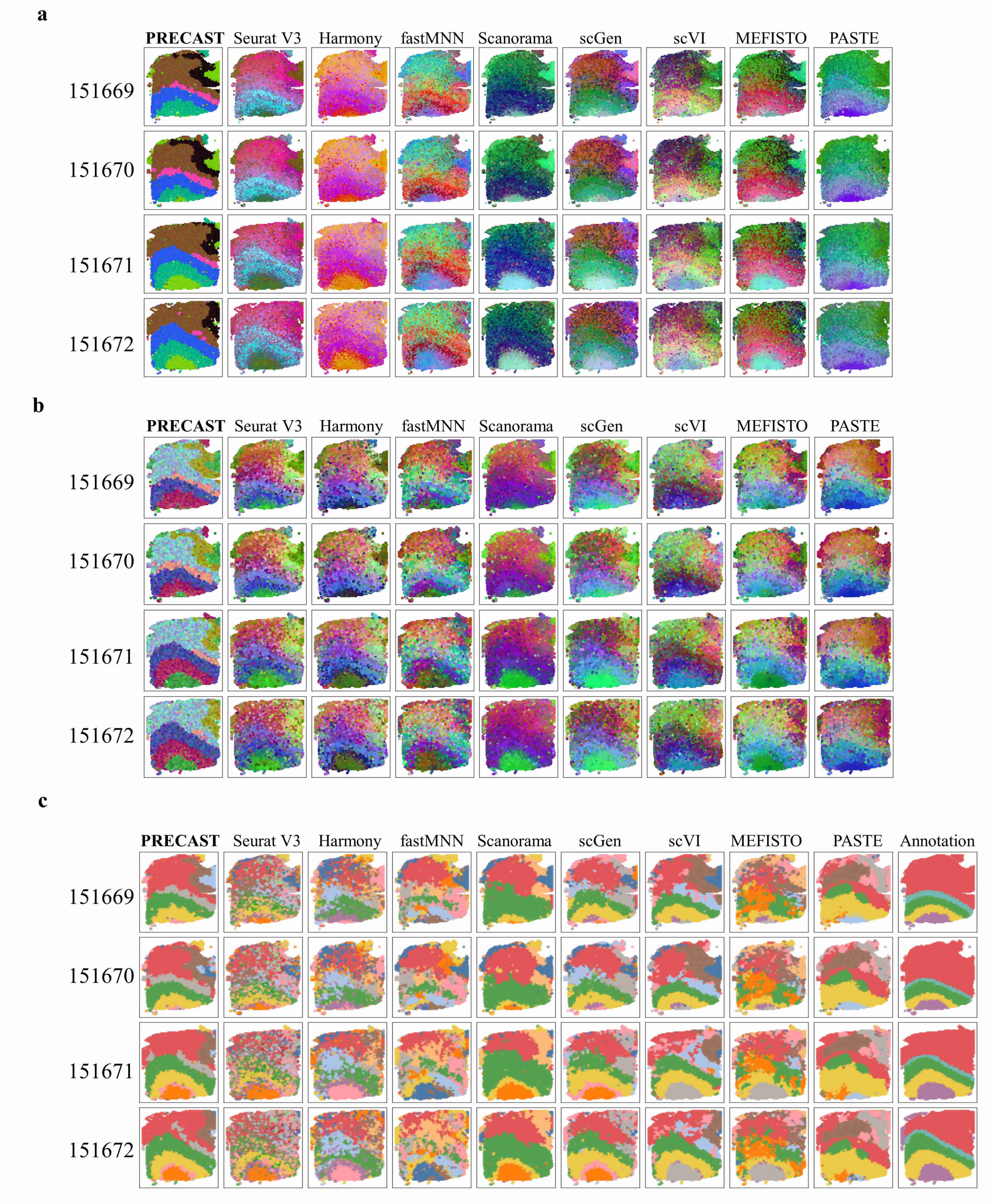


Figure S5: Dimension reduction with batch correction and clustering analysis for the 12 dorsolateral prefrontal cortex Visium sections. a. UMAP RGB plots for samples 5-8 with sample ID 151669-151672 of Donor 2 based on the low-dimensional embeddings from PRECAST and eight other methods. b. tSNE RGB plots for these methods. c. Domain clustering plots for these methods and manual annotations, where SC-MEB is used in the other methods for clustering based on the low-dimensional embeddings.

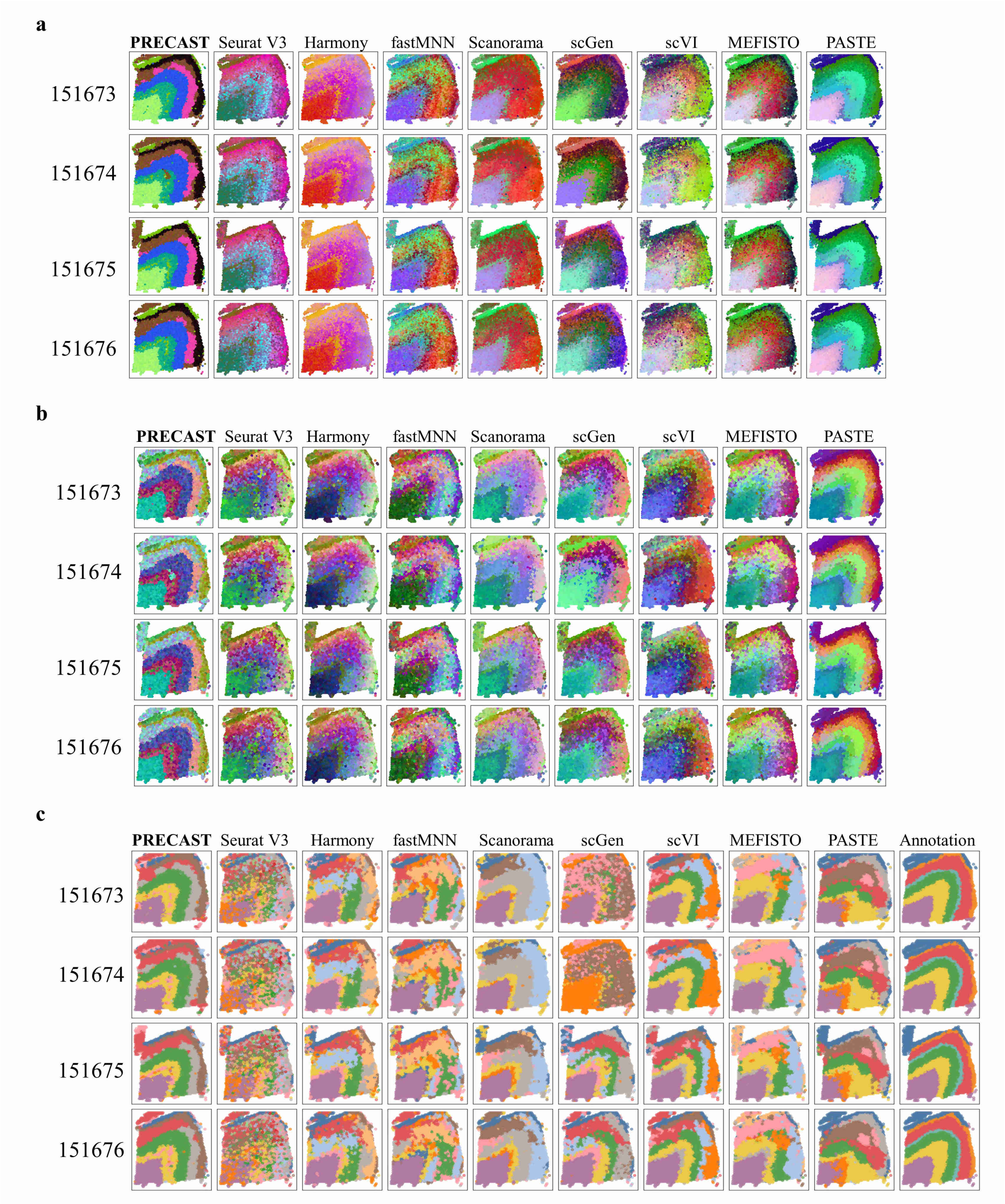


Figure S6: Dimension reduction with batch correction and clustering analysis for the 12 dorsolateral prefrontal cortex Visium sections. a. UMAP RGB plots for samples 9-12 with sample ID 151673-151676 of Donor 3 based on the low-dimensional embeddings from PRECAST and eight other methods. b. tSNE RGB plots for these methods. c. Domain clustering plots for these methods and manual annotations, where SC-MEB is used in the other methods for clustering based on their low-dimensional embeddings.

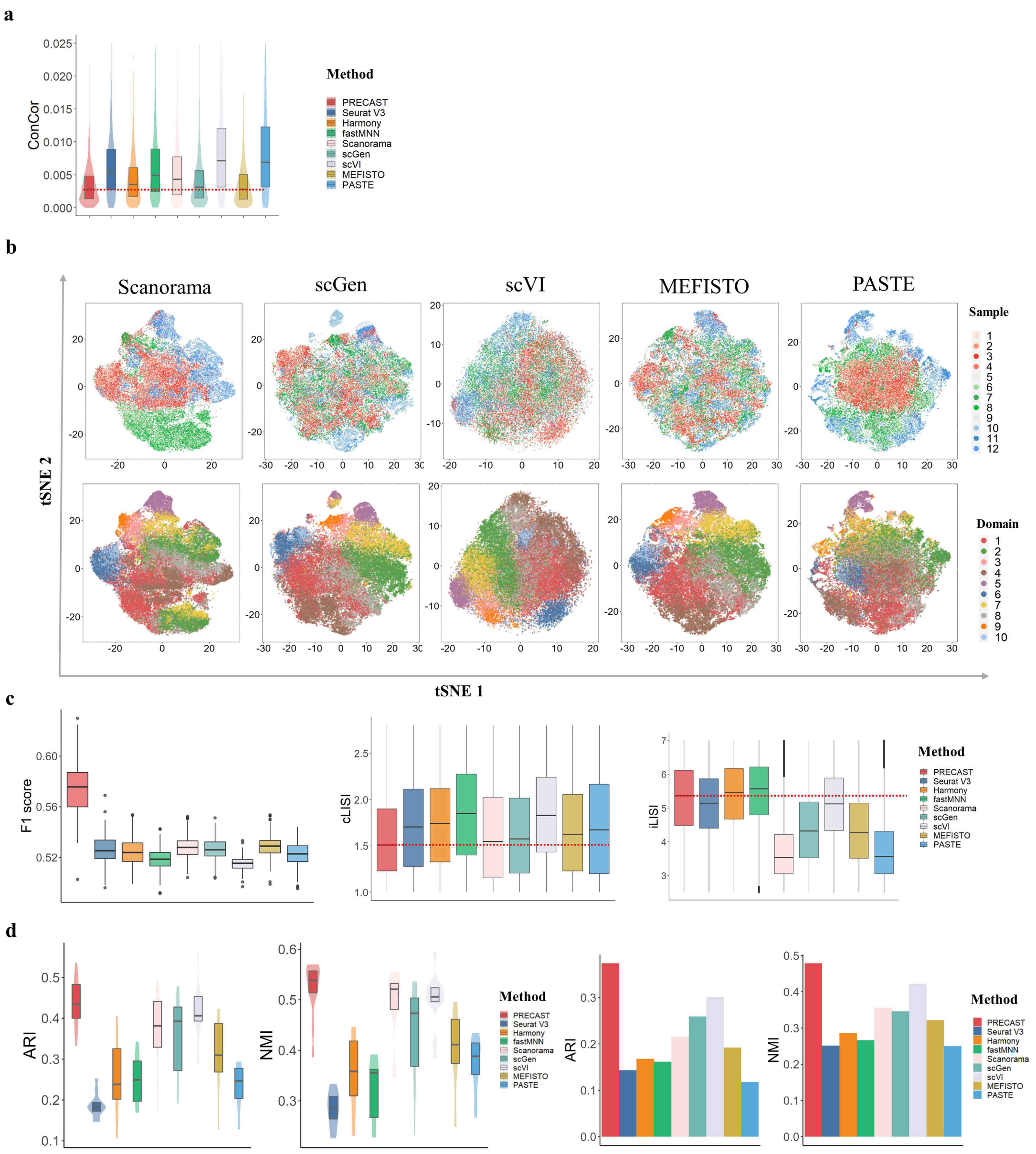


Figure S7: Dimension reduction with batch correction and clustering analysis for the 12 dorsolateral prefrontal cortex Visium sections ($n = 47,680$ locations over 12 tissue sections). a. Boxplot/violin plot of conditional correlations from PRECAST and eight other methods. b. tSNE plots of the data batch/spatial domain labels from PRECAST for five other methods. c. Boxplot of F1 score (F1 score of the average silhouette coefficients), cLISI and iLISI for PRECAST and eight other methods. d. Boxplot/violin plot of ARIs/NMIs for each sample from PRECAST, and SC-MEB clustering based on the low-dimensional embeddings of other compared methods, and bar plot of ARIs/NMIs for 12 combined samples from PRECAST and other methods. In the boxplot, the center line, box lines and whiskers represent the median, upper, and lower quartiles, and 1.5 times interquartile range, respectively.

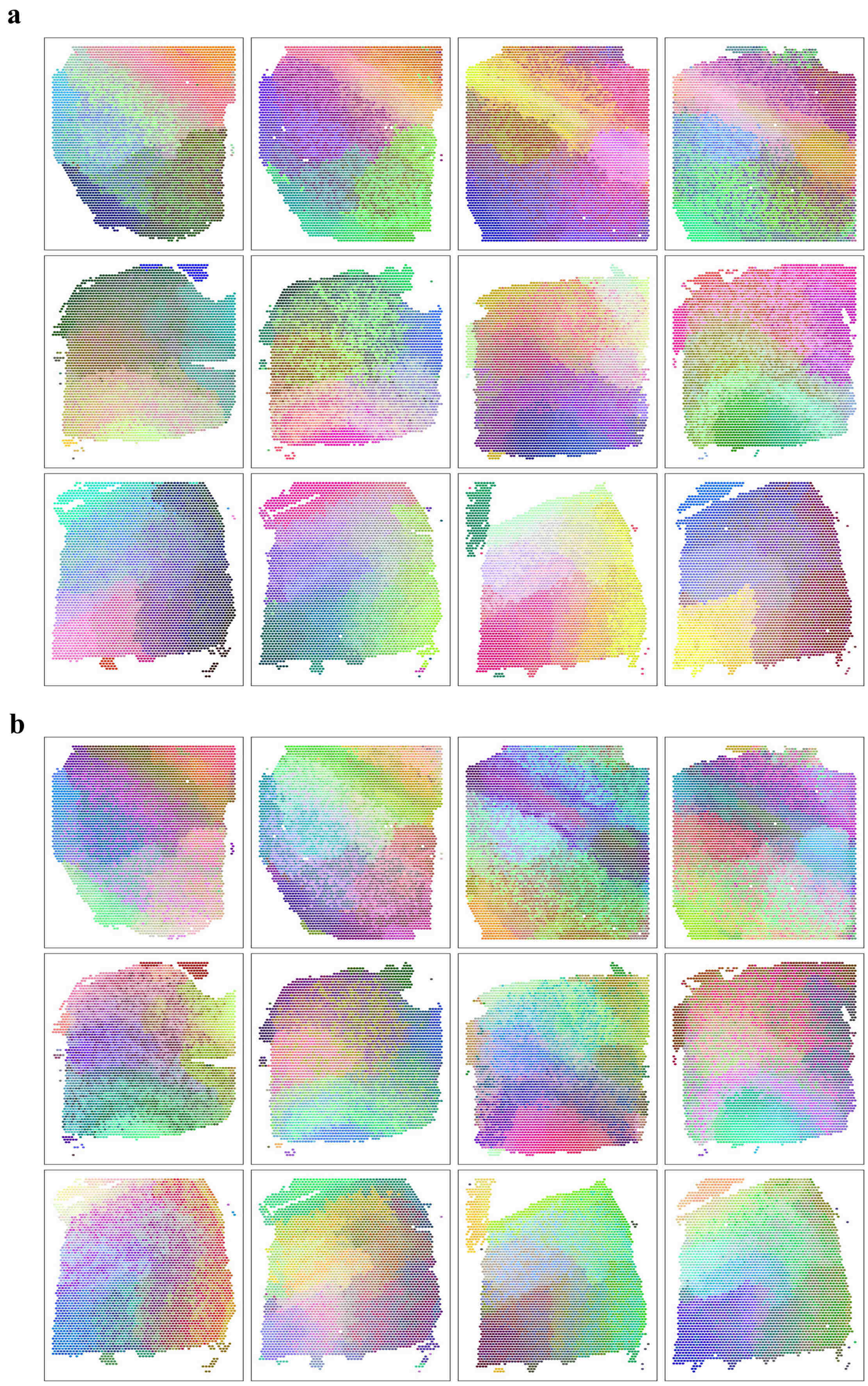


Figure S8: Microenvironment spatial dependence analysis for the 12 dorsolateral prefrontal cortex Visium sections. a. UMAP RGB plot of the inferred embeddings from the intrinsic CAR of PRECAST for 12 sections. First row, sections with sample ID 151507-151510 of donor 1; second row, sections with sample ID 151669-151672 of donor 2; third row, sections with sample ID 151673-151676 of donor 3. b. tSNE RGB plot of the inferred embeddings from the intrinsic CAR of PRECAST for the 12 sections with the same layout.

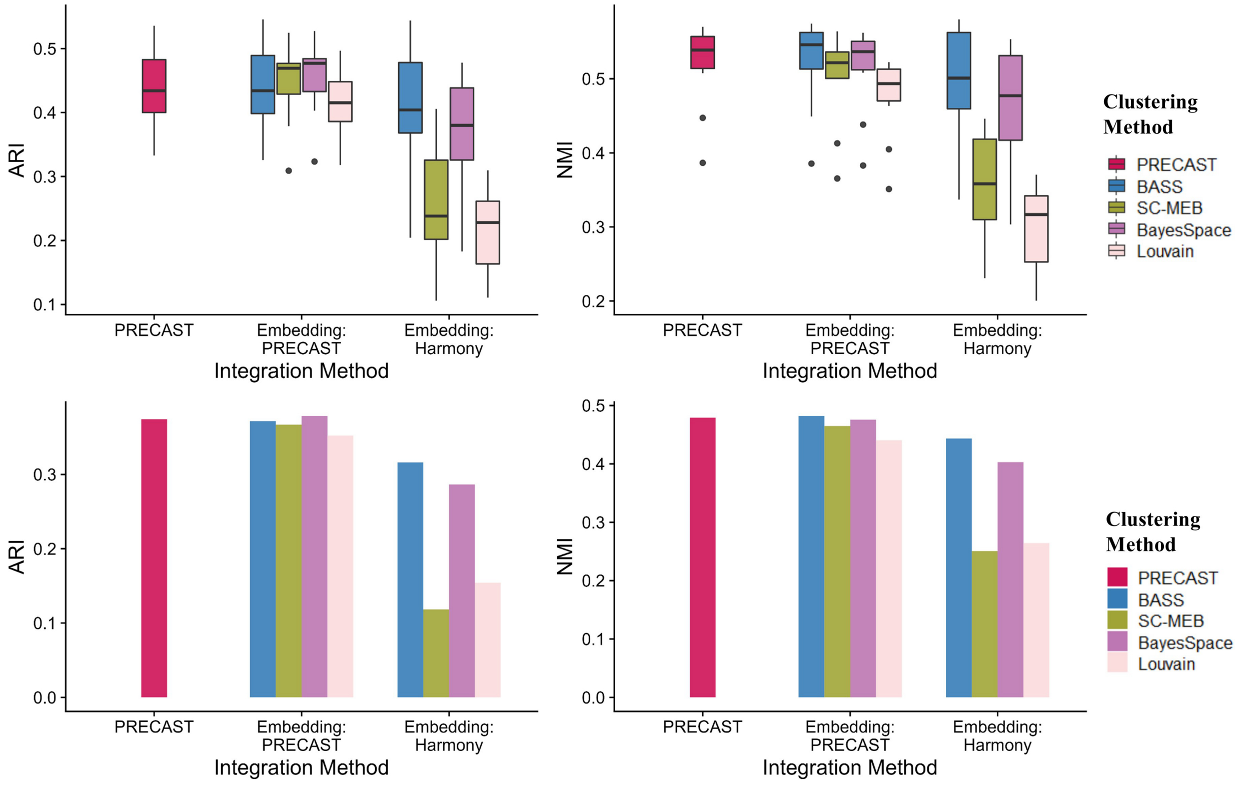


Figure S9: Clustering analysis based on different embeddings for the 12 dorsolateral prefrontal cortex Visium sections ($n = 47,680$ locations over 12 tissue sections). Upper panel: Box plot of ARIs/NMIs for each sample from PRECAST, BASS, SC-MEB, BayesSpace and Louvain clustering, based on the low-dimensional embeddings of PRECAST and Harmony. In the boxplot, the center line, box lines and whiskers represent the median, upper, and lower quartiles, and 1.5 times interquartile range, respectively. Bottom panel: Bar plot of ARI/NMI for combined samples from BASS, SC-MEB, BayesSpace and Louvain clustering, based on the low-dimensional embeddings of PRECAST and Harmony.

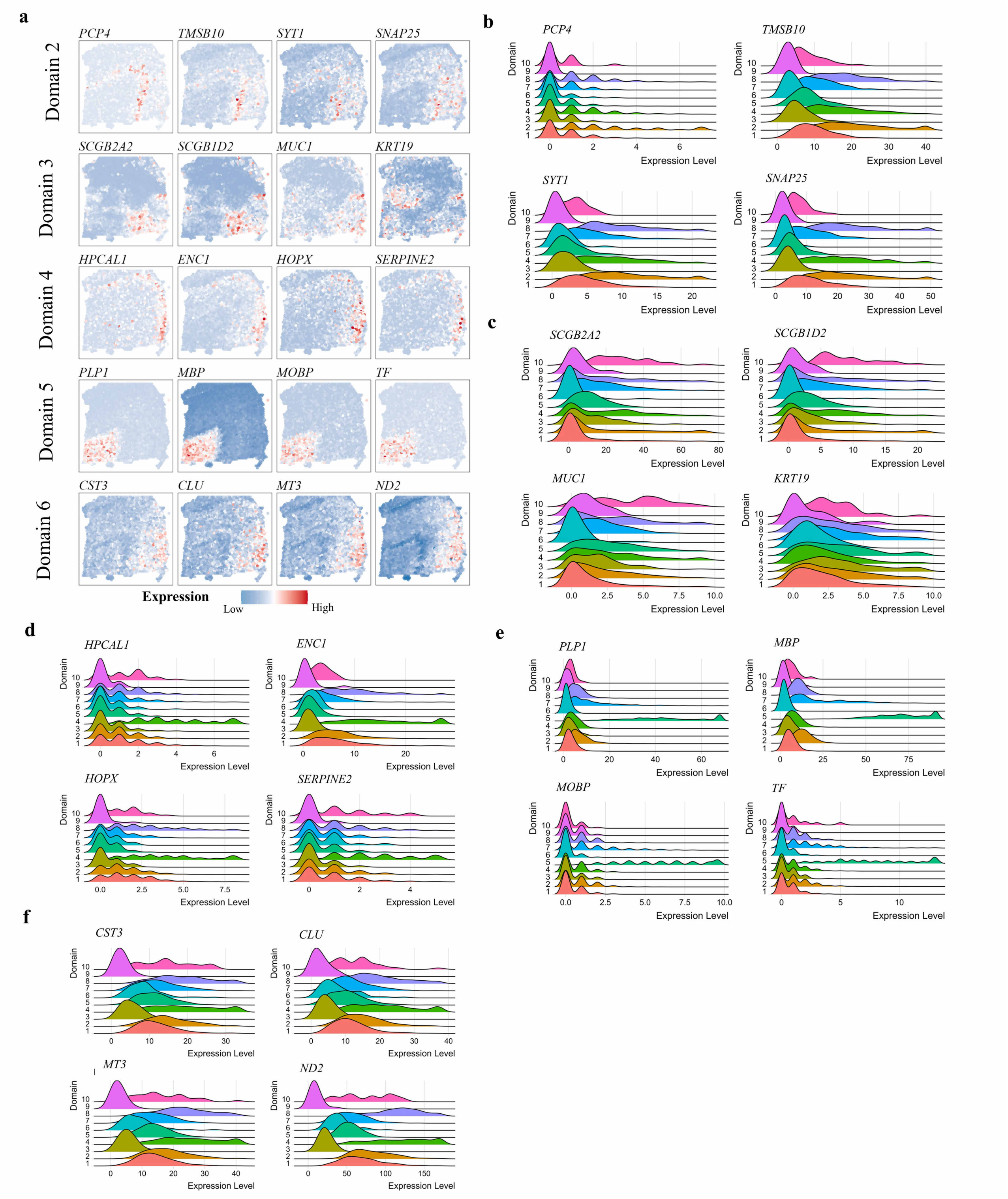


Figure S10: Combined DE analysis of the 12 dorsolateral prefrontal cortex Visium sections. a. Spatial heatmap of scaled expression of top four DE genes for Domains 2-6 of Sample 10 with ID 151674. (b)-(f): Ridge plots of raw gene expression of top four DEGs in each of Domains 2-6 in Sample 10.

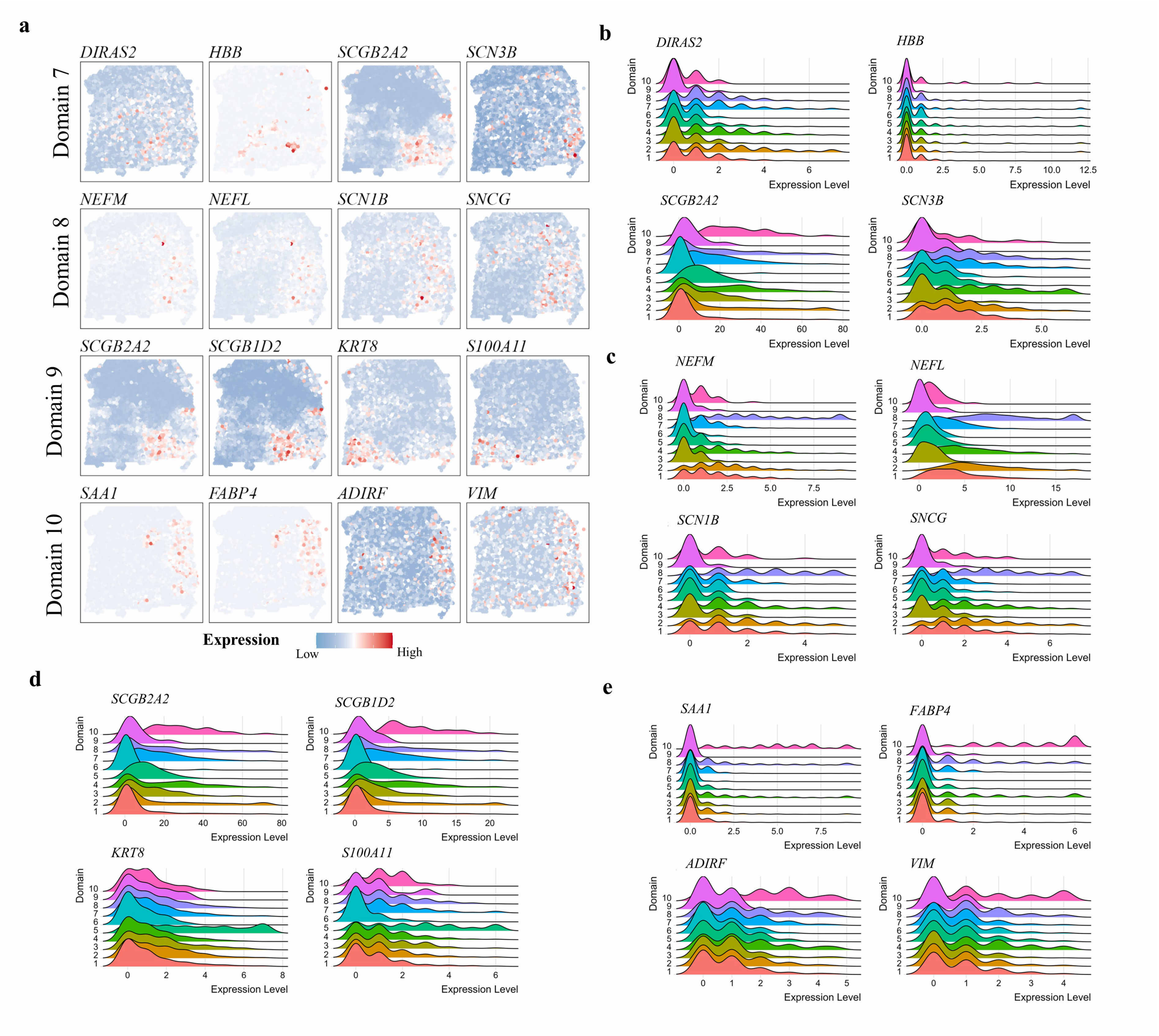


Figure S11: Combined DE analysis of the 12 dorsolateral prefrontal cortex Visium sections. a. Spatial heatmap of scaled expression of top four DE genes for Domains 7-10 of Sample 10 with ID 151674. (b)-(e): Ridge plots of raw expression of top four DE genes in each of Domains 7-10 in Sample 10.

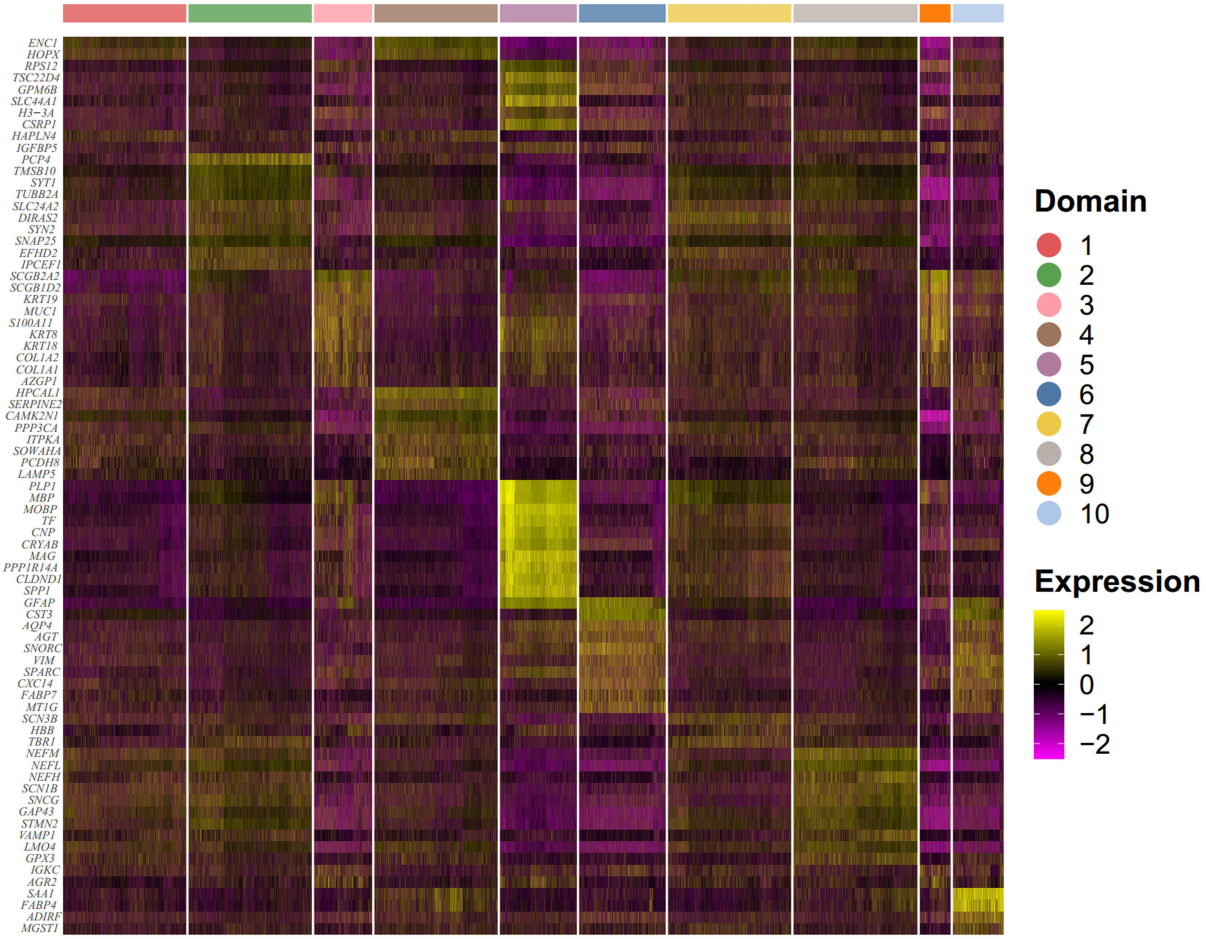


Figure S12: Combined DE analysis of the 12 dorsolateral prefrontal cortex Visium sections. Heatmap of top 10 differentially expressed genes for each domain identified by PRECAST.

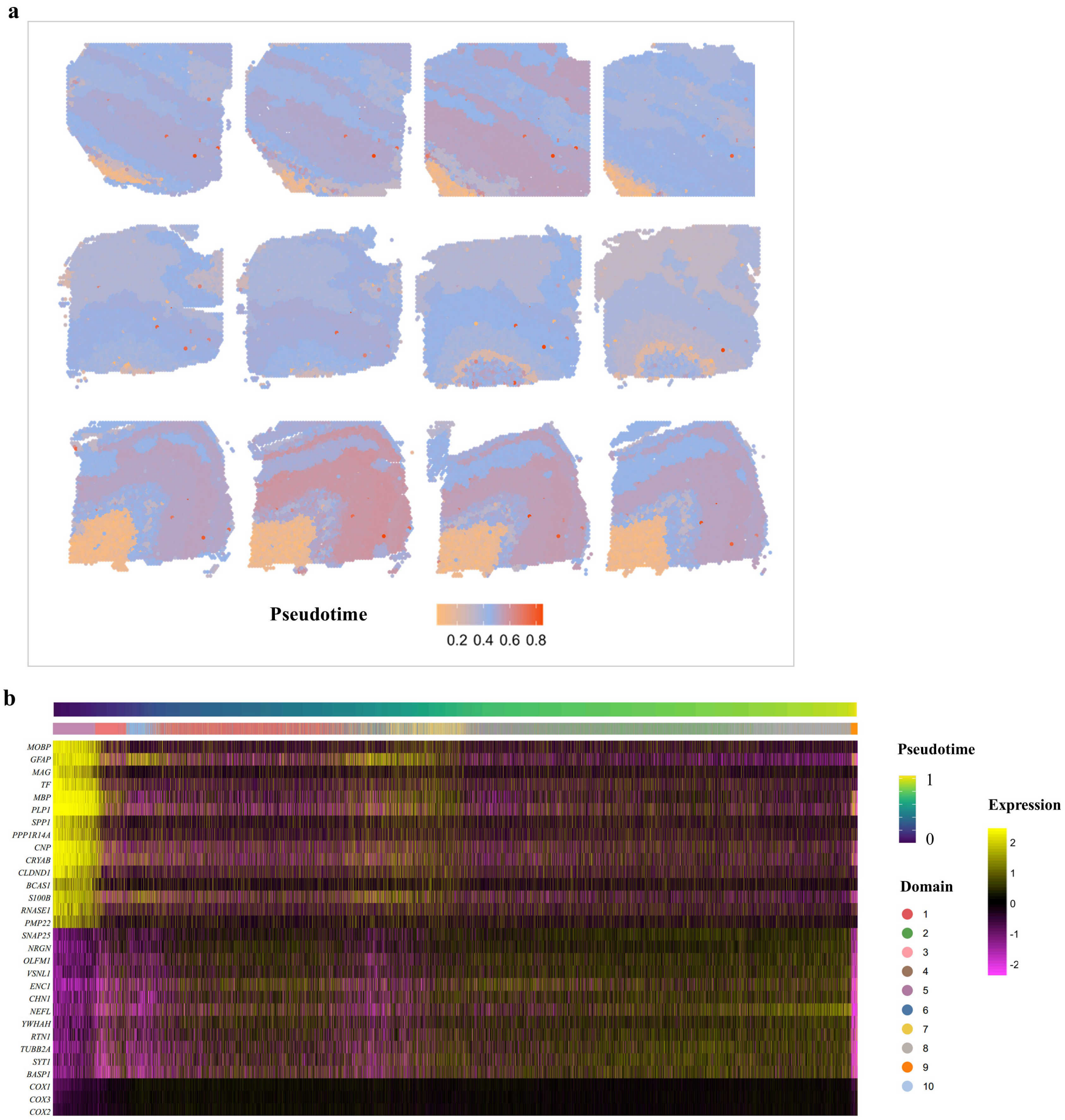


Figure S13: Combined trajectory analysis for 12 samples from the dorsolateral prefrontal cortex Visium sections. a. Spatial heatmap of the inferred scaled pseudotime by Slingshot. b. Heatmap of PRECAST batch-corrected expression levels of top 30 genes with significant changes with respect to the scaled Slingshot pseudotime. Each column represents a spot that is mapped to this path and ordered by its pseudotime value. Each row denotes a top significant gene.

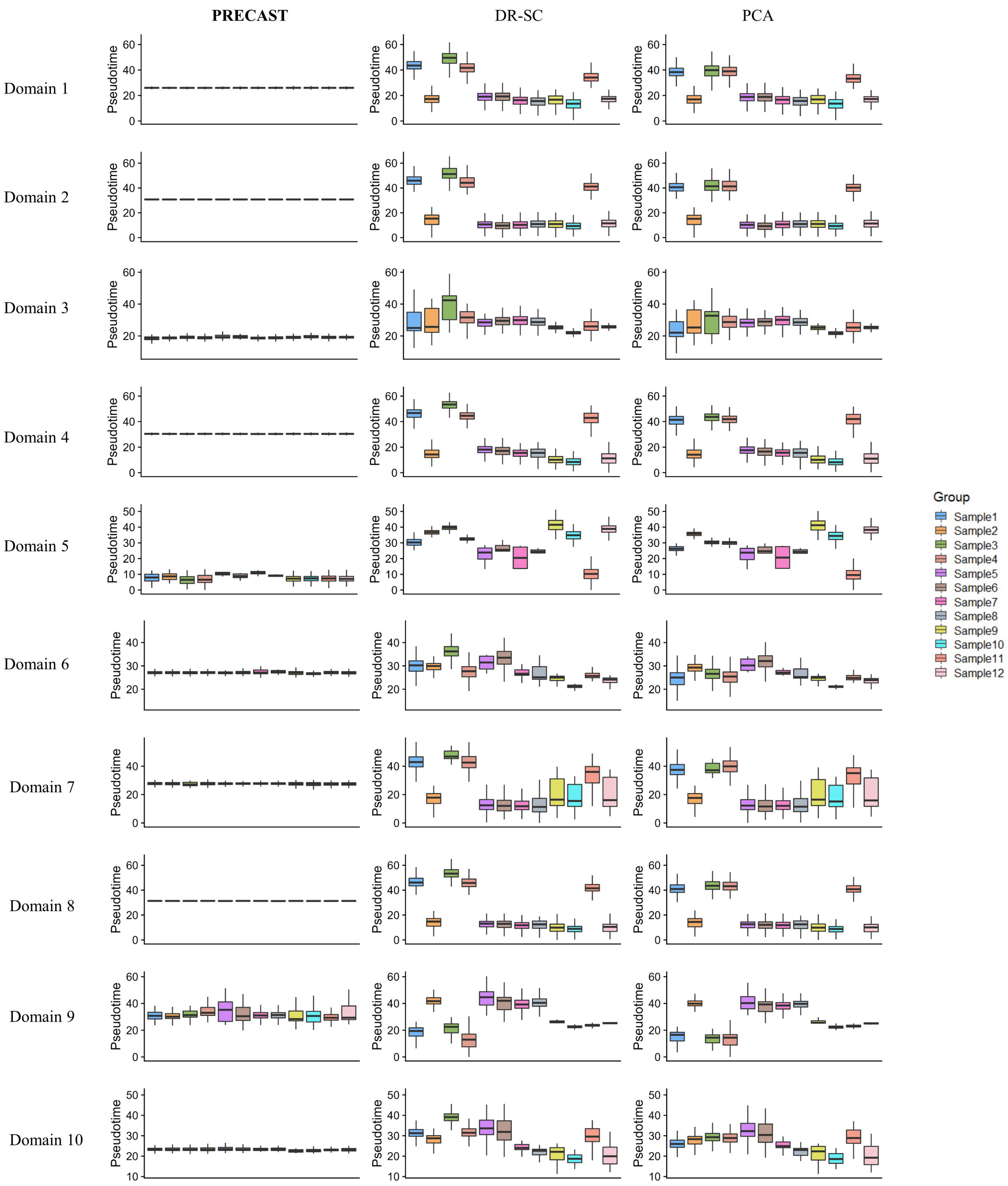


Figure S14: Comparison of the inferred pseudotime with embeddings from PRECAST, DR-SC and PCA for the 12 dorsolateral prefrontal cortex Visium sections ($n = 47,680$ locations over 12 tissue sections). PCA and DR-SC were performed on each slide to obtain the embeddings while PRECAST was performed on all slides to obtained aligned embeddings. In the boxplot, the center line, box lines and whiskers represent the median, upper, and lower quartiles, and 1.5 times interquartile range, respectively.

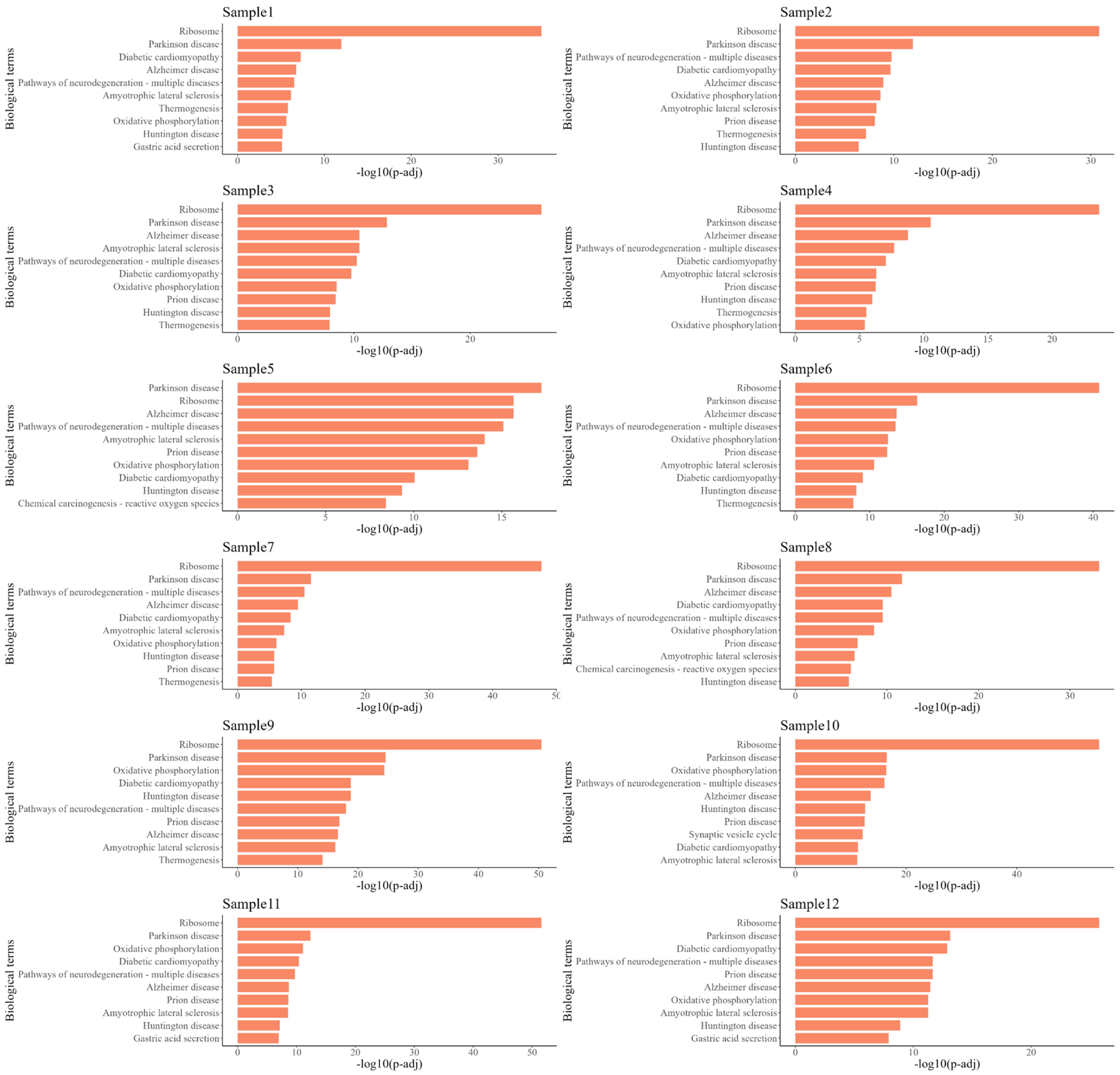

Figure S15: Enrichment analysis of conditional SVGs based on low-dimensional embeddings extracted by PRECAST from the 12 dorsolateral prefrontal cortex Visium sections. Bar plot of -log10(adjusted $p$-value) for top 10 KEGG pathways in enrichment analysis of SVGs of each sample. The $p$-values are based on one-sided Fisher's exact tests with multiple testing corrections using the Benjamini-Hochberg FDR method.

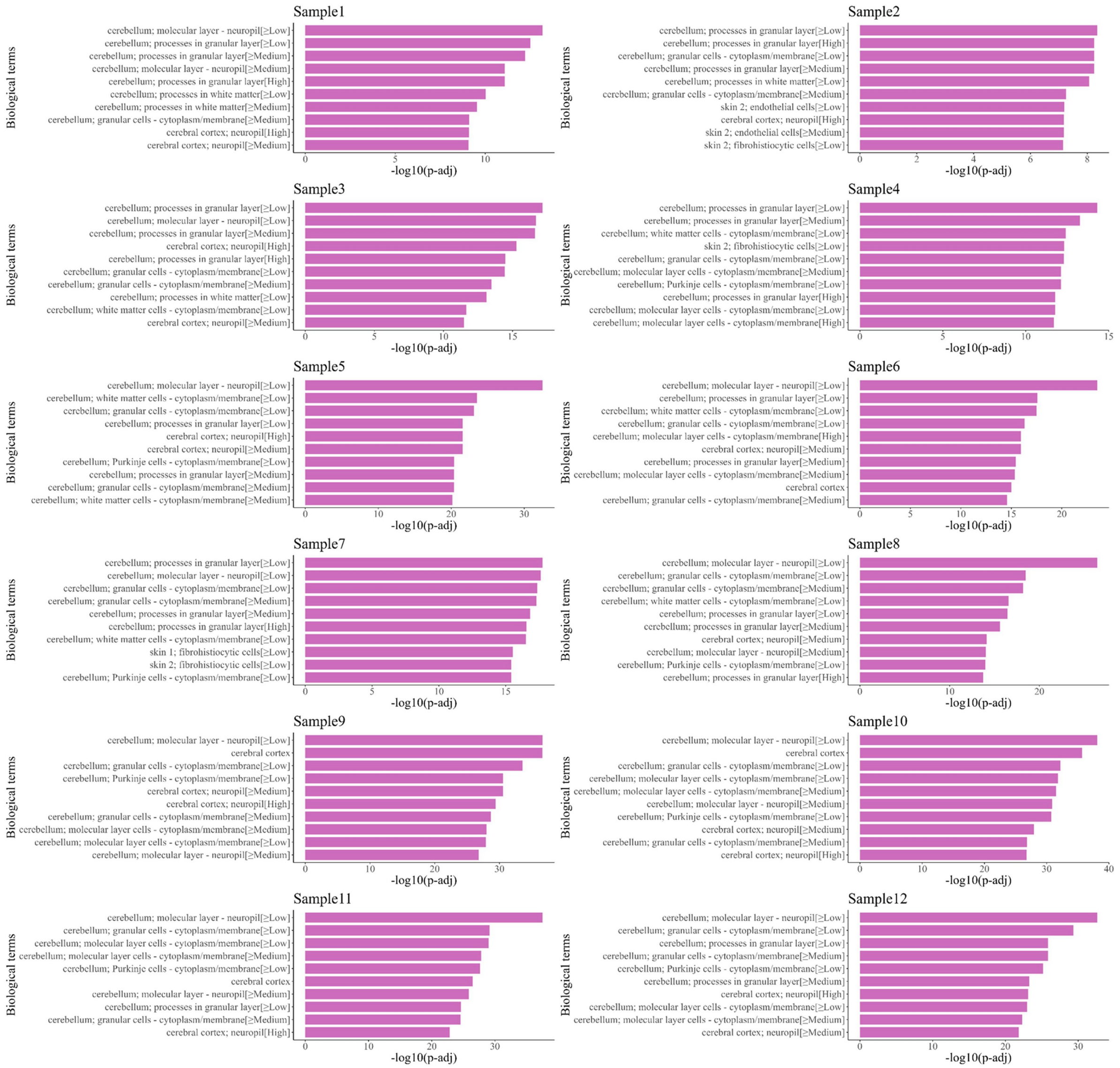

Figure S16: Enrichment analysis of conditional SVGs based on low-dimensional embeddings extracted by PRECAST from the 12 dorsolateral prefrontal cortex Visium sections. Bar plot of -log10(adjusted $p$-value) for top 10 HPA pathways in enrichment analysis of SVGs of each sample. The $p$-values are based on one-sided Fisher's exact tests with multiple testing corrections using the Benjamini-Hochberg FDR method.

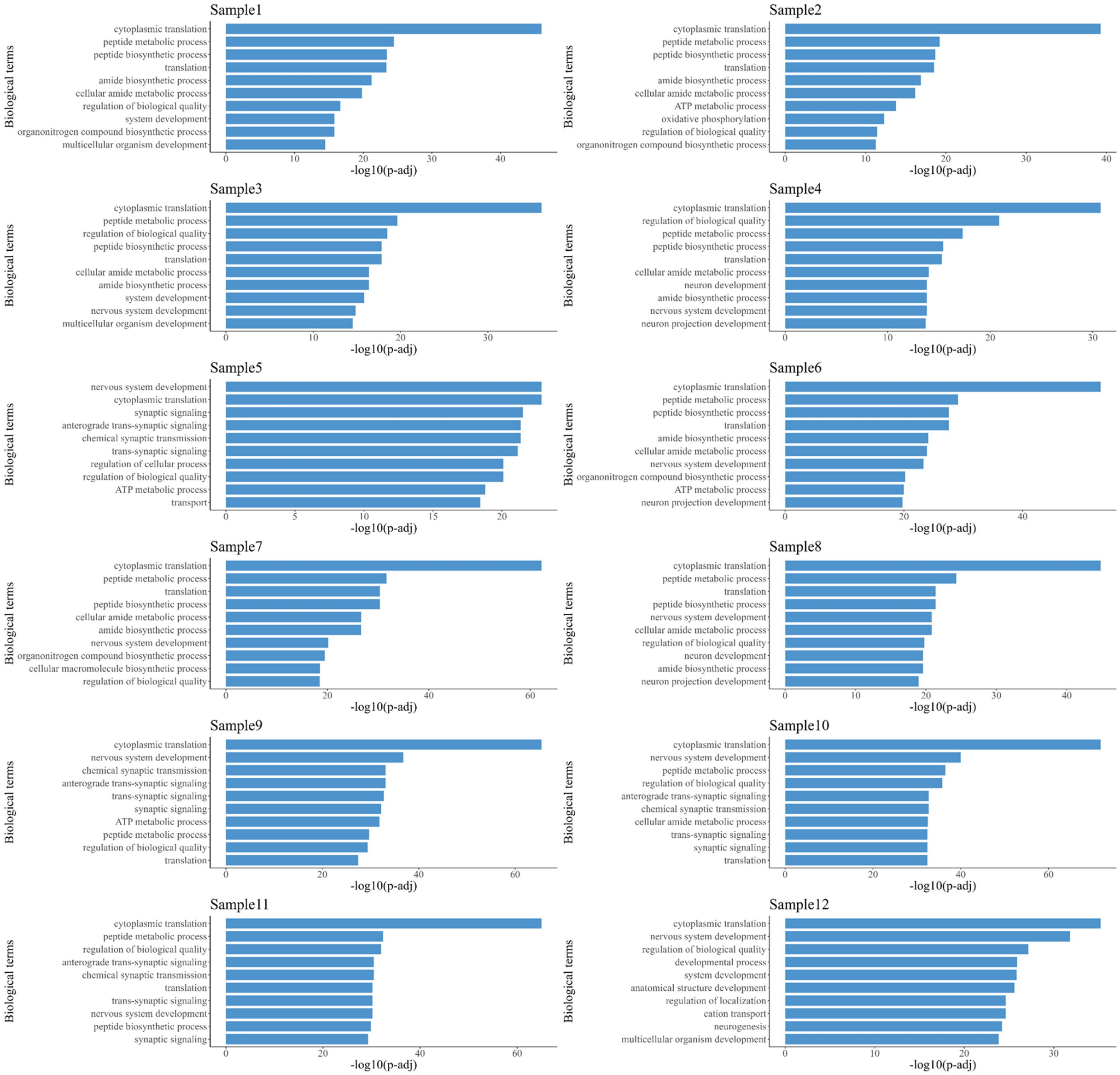


Figure S17: Enrichment analysis of conditional SVGs based on low-dimensional embeddings extracted by PRECAST from the 12 dorsolateral prefrontal cortex Visium sections. Bar plot of -log10(adjusted $p$-value) for top 10 GO biological process (BP) pathways in enrichment analysis of SVGs of each sample. The $p$-values are based on one-sided Fisher's exact tests with multiple testing corrections using the Benjamini-Hochberg FDR method.

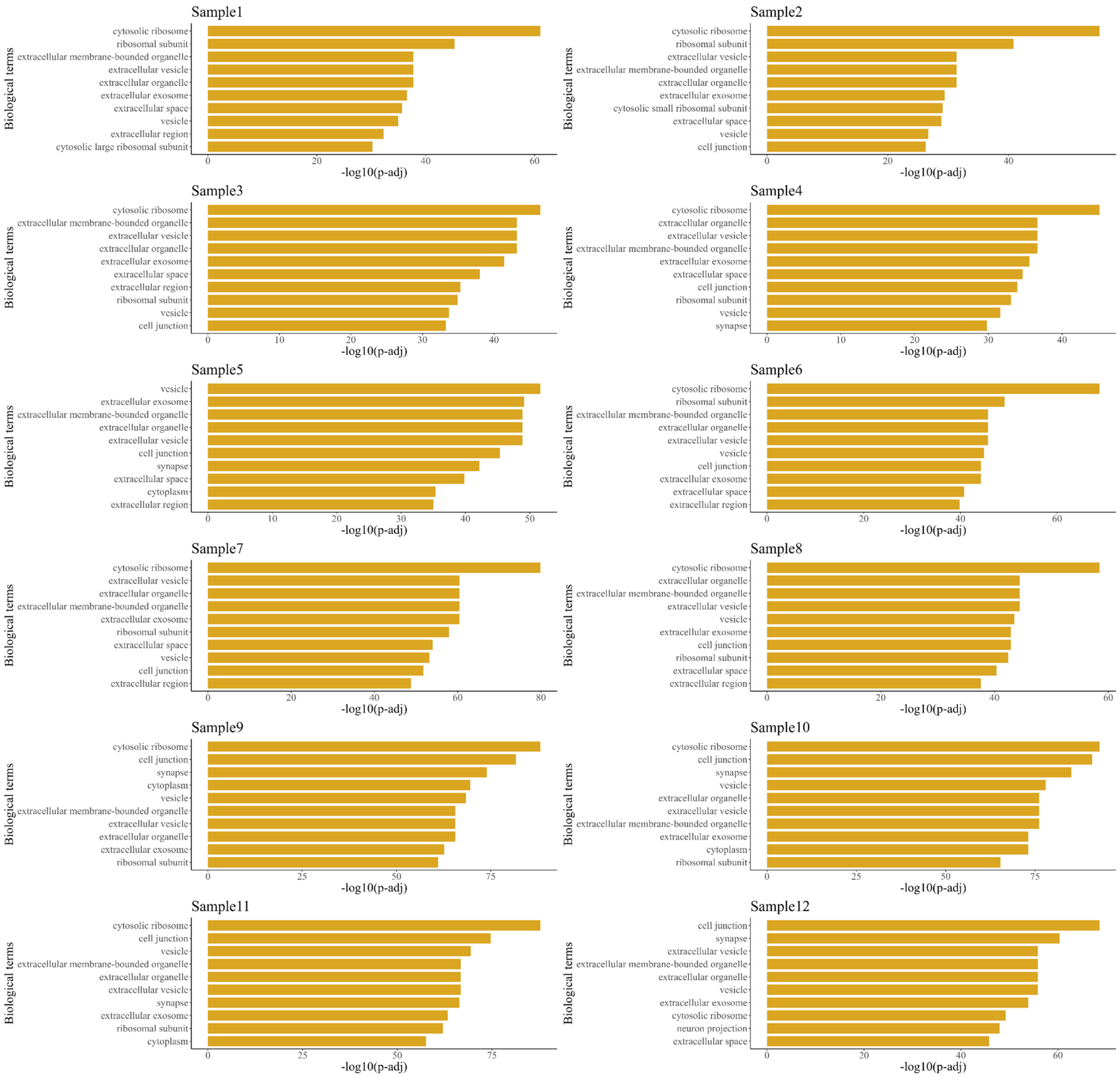


Figure S18: Enrichment analysis of conditional SVGs based on low-dimensional embeddings extracted by PRECAST from the 12 dorsolateral prefrontal cortex Visium sections. Bar plot of -log10(adjusted $p$-value) for top 10 GO cellular component (CC) pathways in enrichment analysis of SVGs of each sample. The $p$-values are based on one-sided Fisher's exact tests with multiple testing corrections using the Benjamini-Hochberg FDR method.

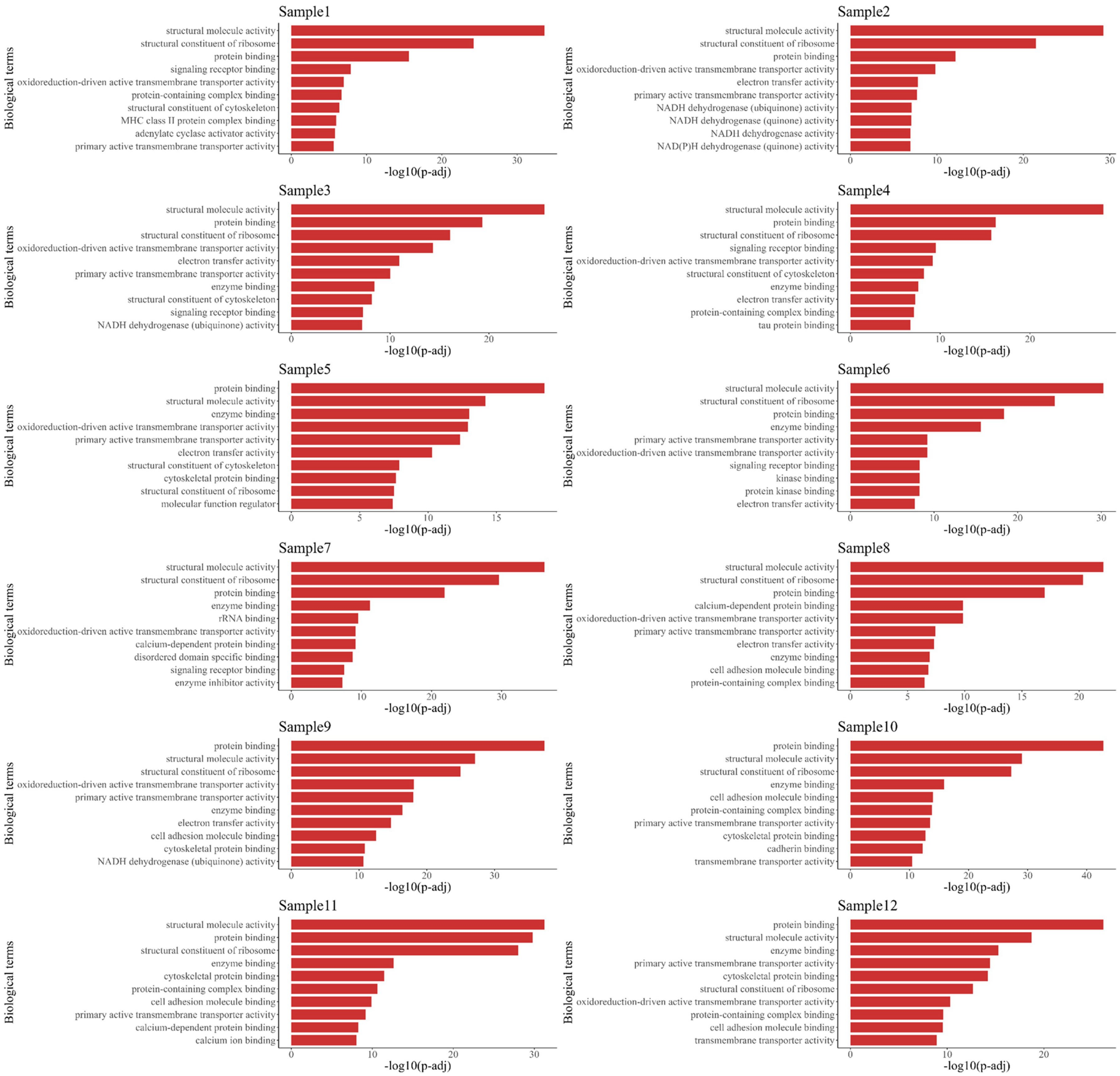


Figure S19: Enrichment analysis of conditional SVGs based on low-dimensional embeddings extracted by PRECAST from the 12 dorsolateral prefrontal cortex Visium sections. Bar plot of -log10(adjusted $p$-value) for top 10 GO molecular function (MF) pathways in enrichment analysis of SVGs of each sample. The $p$-values are based on one-sided Fisher's exact tests with multiple testing corrections using the Benjamini-Hochberg FDR method.

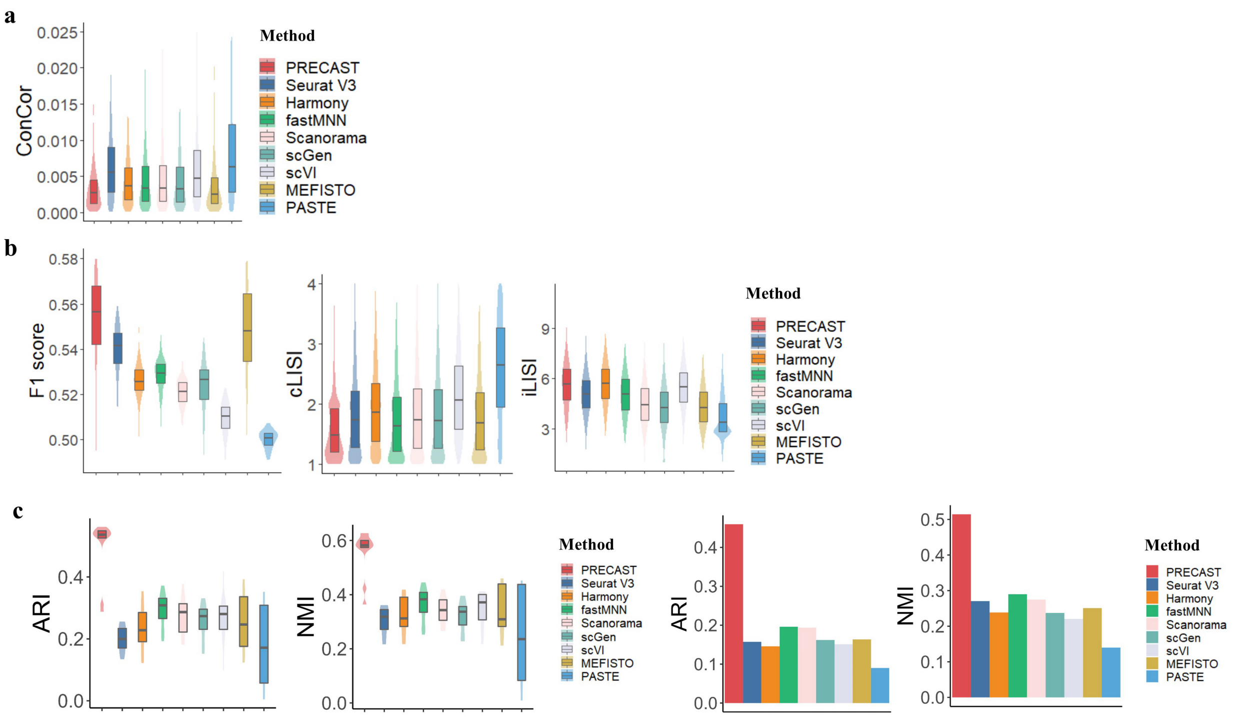


Figure S20: Dimension reduction with batch correction and clustering analysis using top 2000 HVGs for the 12 dorsolateral prefrontal cortex Visium sections ($n = 47,680$ locations over 12 tissue sections). a. Boxplot/violin plot of conditional correlations from PRECAST and eight other methods. b. Boxplot of F1 score (F1 score of the average silhouette coefficients), cLISI and iLISI for PRECAST and eight other methods. c. Boxplot/violin plot of ARIs/NMIs for each sample from PRECAST and SC-MEB clustering based on the low-dimensional embeddings of other compared methods, and bar plot of ARIs/NMIs for 12 combined samples from PRECAST and other methods. In the boxplot, the center line and box lines denote the median, upper, and lower quartiles, respectively.

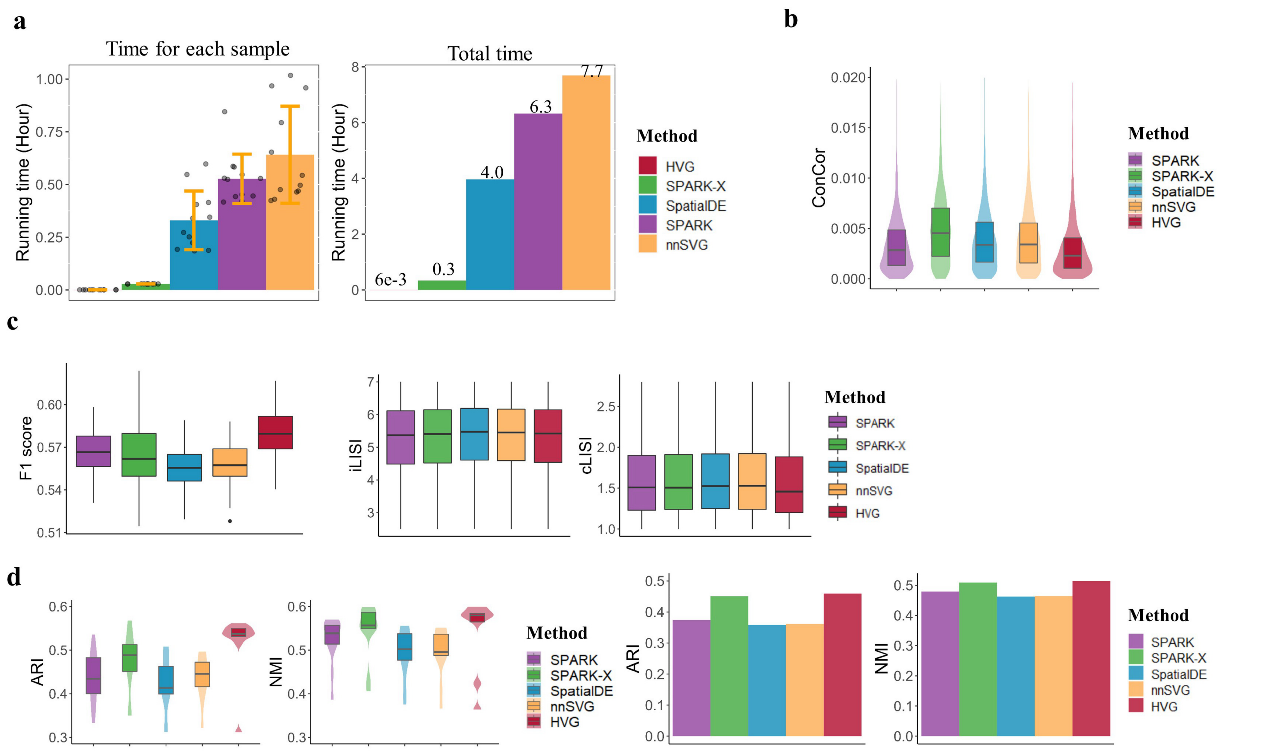


Figure S21: Performance comparison of results from PRECAST using SPARK, SPARK-X, SpatialDE, nnSVG and HVGs gene selection methods, for the 12 dorsolateral prefrontal cortex Visium sections ($n = 47,680$ locations over 12 tissue sections). a. Barplot of running times for each sample (left panel) and all samples. In the bar plot, the error bands represent the mean value ± standard deviation. b. Boxplot/violin plot of conditional correlations. c. Boxplot of F1 score of the average silhouette coefficients, cLISI and iLISI. d. Boxplot/violin plot of ARIs/NMIs for each sample, and bar plot of ARIs/NMIs for the 12 combined samples. In the boxplot, the center line, box lines and whiskers represent the median, upper, and lower quartiles, and 1.5 times interquartile range, respectively.

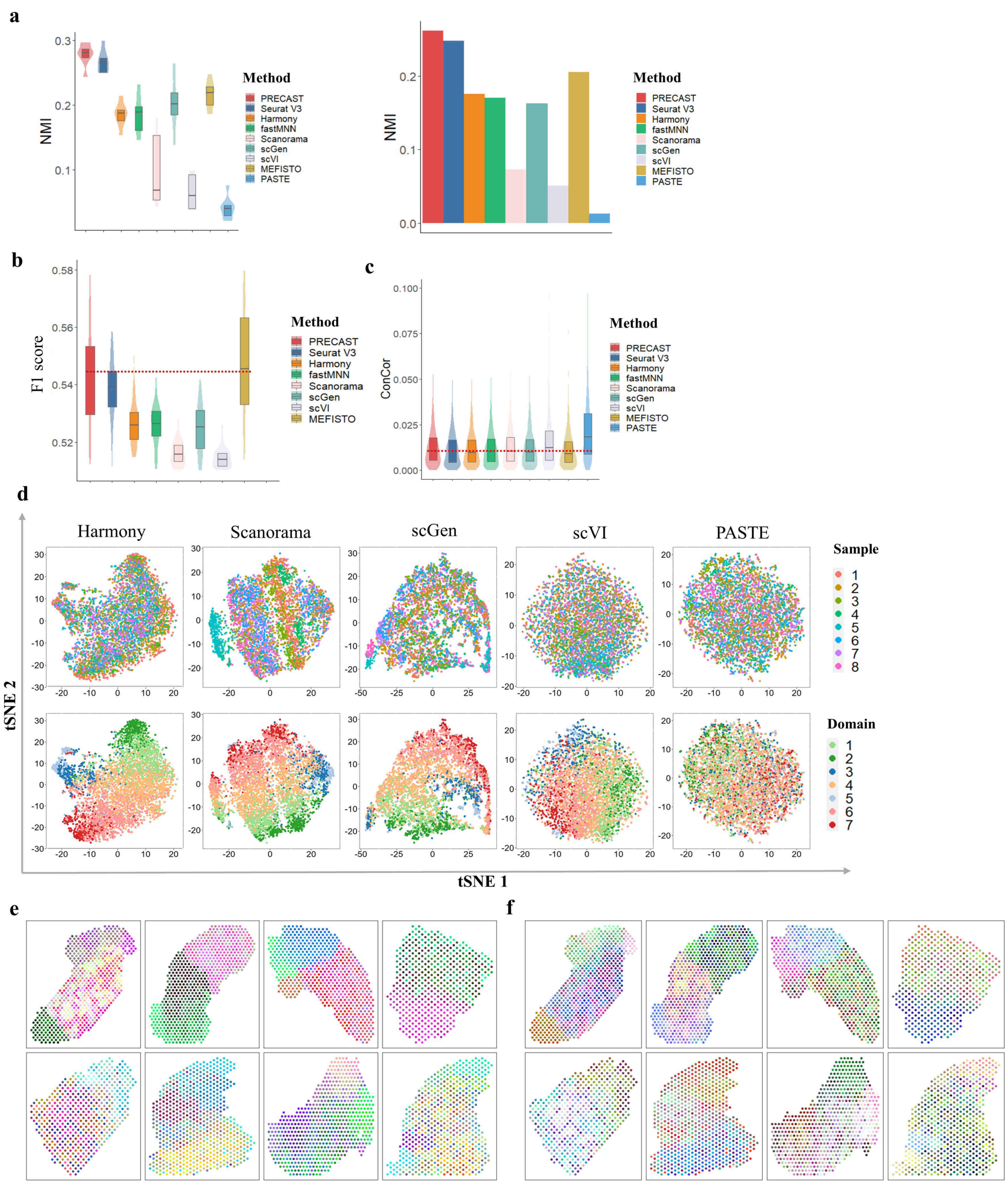


Figure S22: Comparison analysis and spatial microenvironment analysis of mouse liver ST data ($n = 4,865$ locations over 8 tissue sections). a. Boxplot/violin plot of NMIs for each sample from PRECAST, and SC-MEB clustering based on the low-dimensional embeddings of other compared methods, and bar plot of NMIs for eight combined samples from PRECAST and other methods. b. Boxplot of F1 score (F1 score of the average silhouette coefficients) for PRECAST and eight other methods. c. Boxplot/violin plot of conditional correlations from PRECAST and eight other methods. d. tSNE plots of the data batch/spatial domain labels from PRECAST for five other methods. e & f. UMAP/tSNE RGB plot of the inferred embeddings from the intrinsic CAR of PRECAST for the eight samples. In the boxplot, the center line and box lines denote the median, upper, and lower quartiles, respectively.

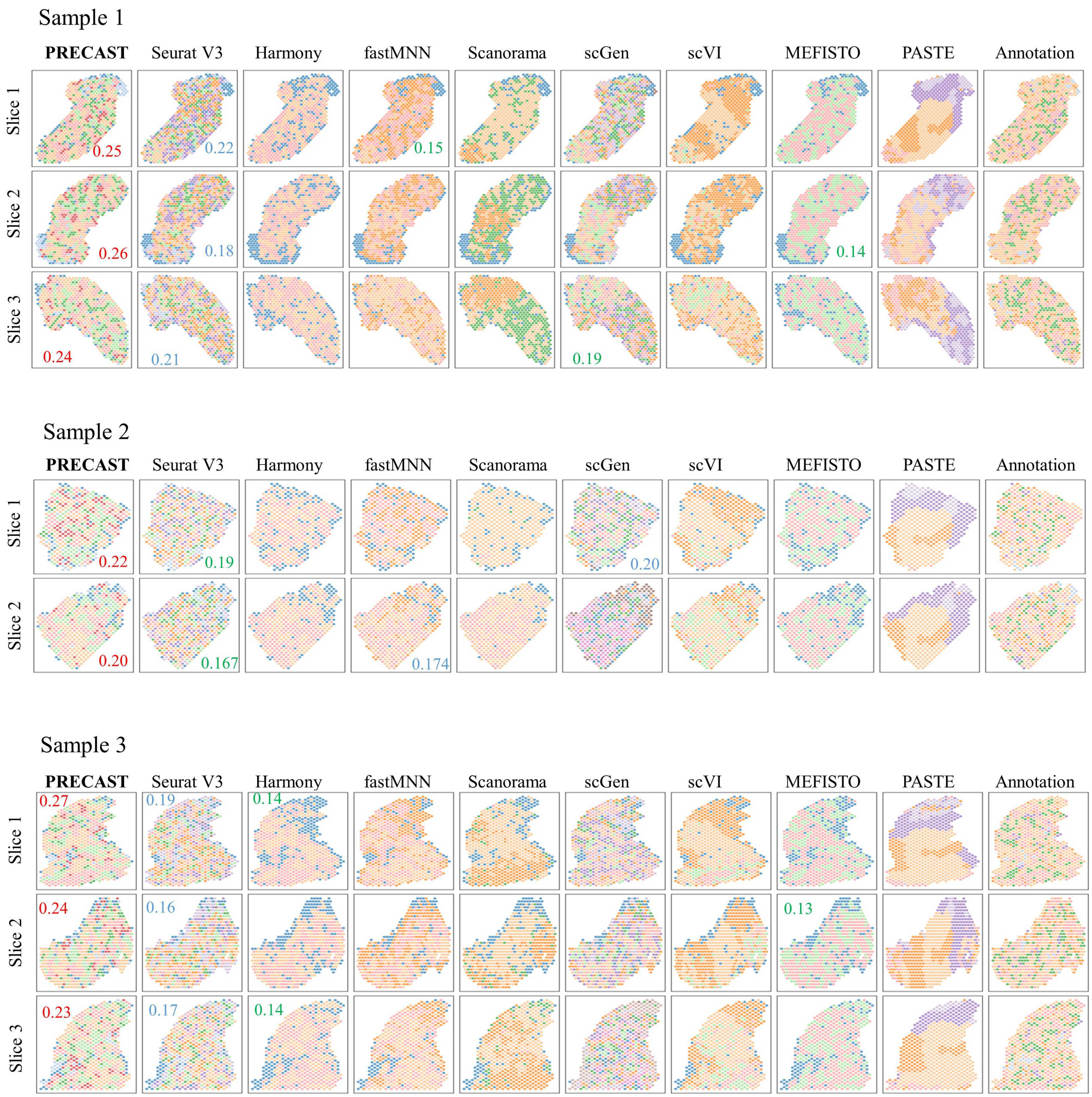


Figure S23: Clustering analysis for the eight mouse liver sections. Domain clustering plots for PRECAST and eight other methods as well as manual annotations, where SC-MEB is used in the other methods for clustering based on their low-dimensional embeddings. The ARIs resulted from top three methods are presented with red, blue and green colors, respectively.

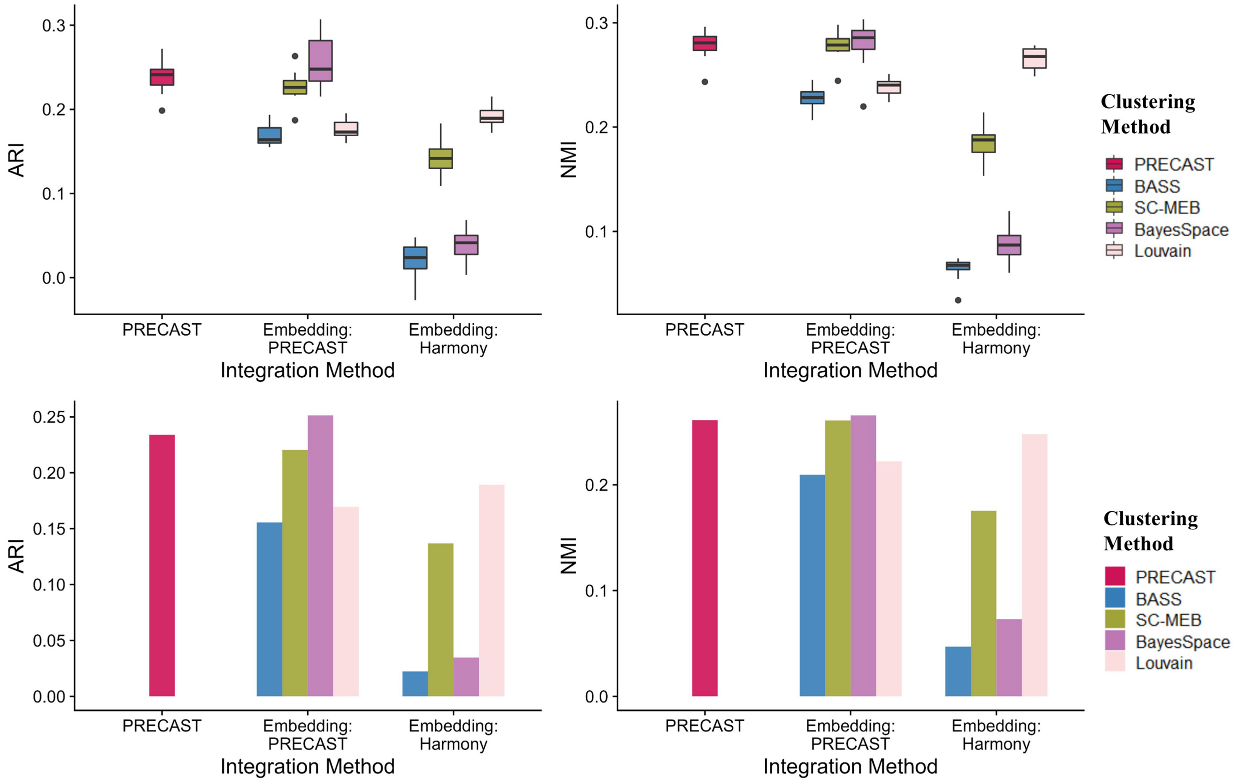


Figure S24: Clustering analysis for mouse liver ST data ($n = 4,865$ locations over 8 tissue sections) based on different embeddings. Upper panel: Box plot of ARIs/NMIs for each sample from PRECAST, BASS, SC-MEB, BayesSpace and Louvain clustering, based on the low-dimensional embeddings of PRECAST and Harmony. In the boxplot, the center line, box lines and whiskers represent the median, upper, and lower quartiles, and 1.5 times interquartile range, respectively. Bottom panel: Bar plot of ARI/NMI for combined samples from BASS, SC-MEB, BayesSpace and Louvain clustering, based on the low-dimensional embeddings of PRECAST and Harmony.

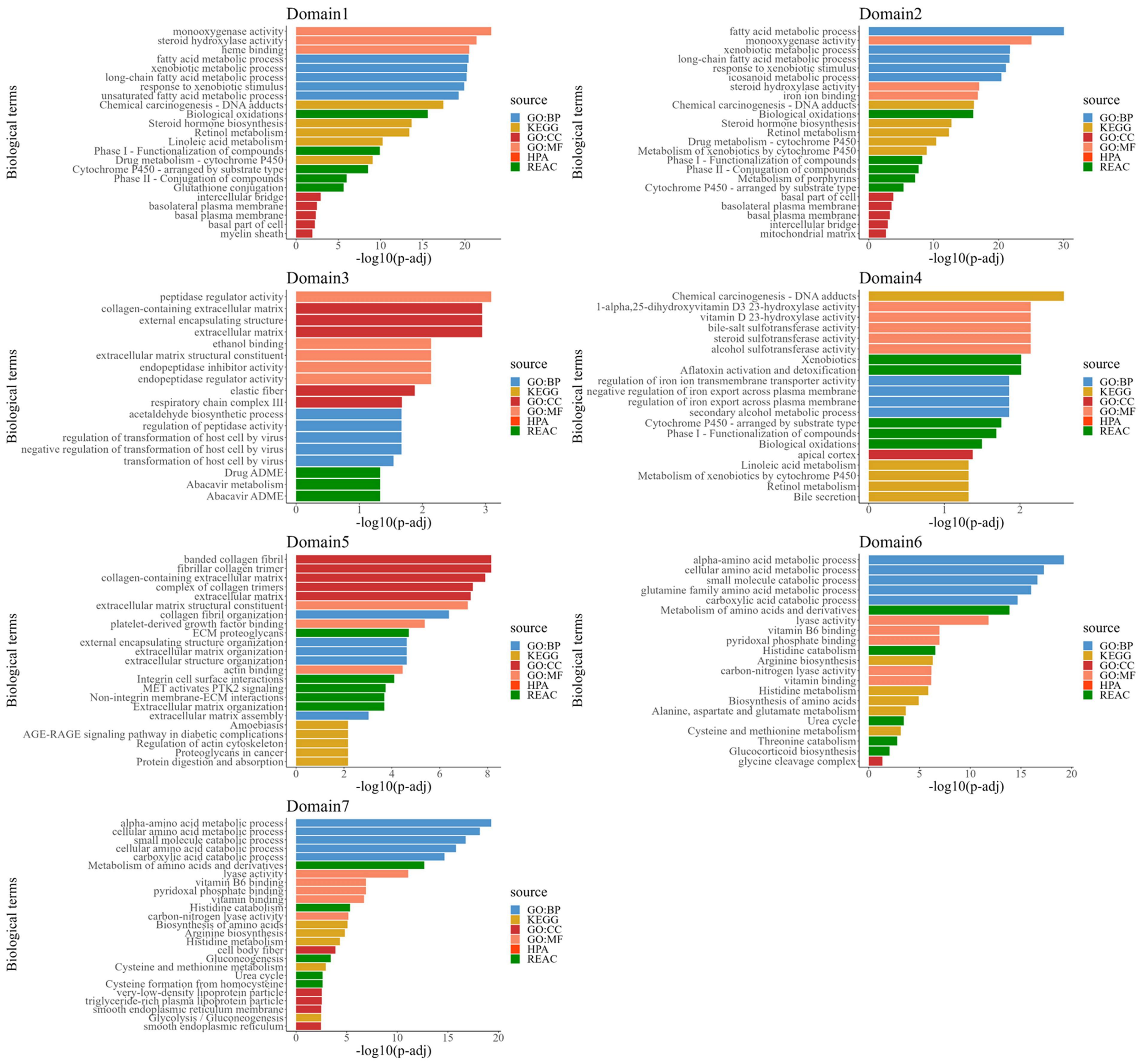


Figure S25: Enrichment analysis of DE genes in data for each domain detected by PRECAST in the eight mouse liver sections. Top five pathways for each category. The $p$-values are based on one-sided Fisher's exact tests with multiple testing corrections using the Benjamini-Hochberg FDR method.

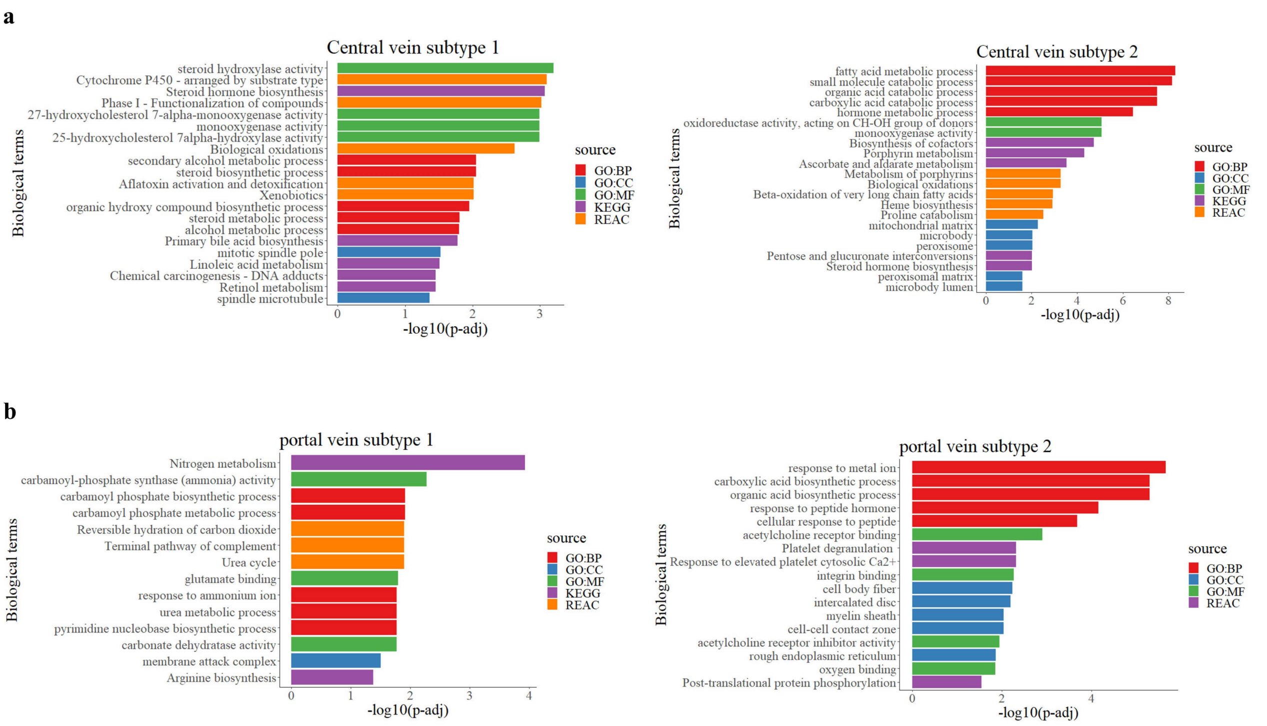


Figure S26: Enrichment analysis of unique DE genes of subtypes detected by PRECAST in the eight mouse liver sections. a. Top five pathways for each category of DE genes corresponding to two subtypes of central veins. b. Top five pathways for each category of DE genes corresponding to two subtypes of portal veins. The $p$-values are based on one-sided Fisher's exact tests with multiple testing corrections using the Benjamini-Hochberg FDR method.

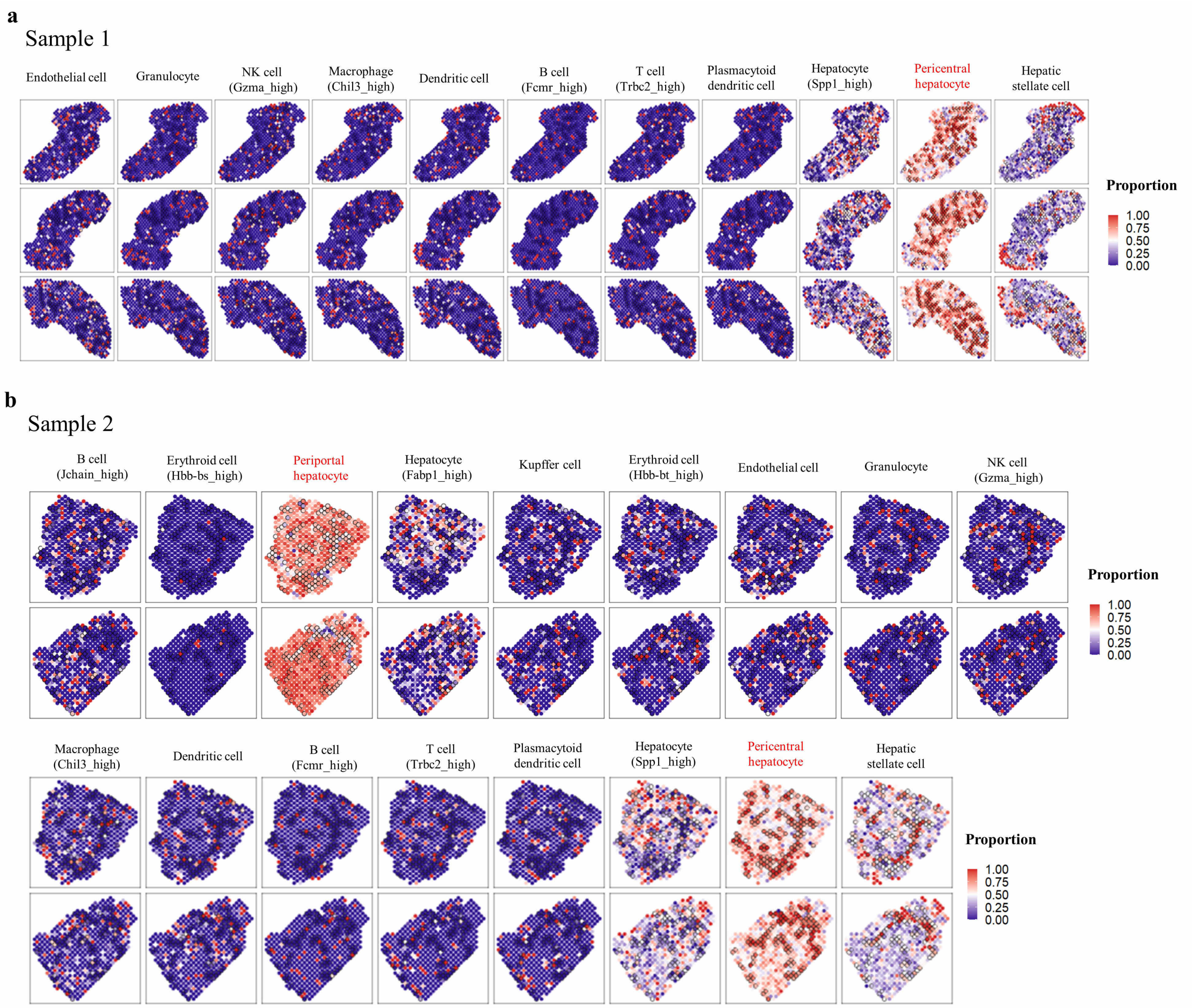


Figure S27: Cell-type deconvolution analysis for the eight mouse liver sections. a. Spatial heatmap of proportions of the remained 11 cell types annotated in MCA on liver tissues for the three slices of Sample 1. b. Spatial heatmap of proportions of 17 cell types annotated in MCA on liver tissues for the two slices of Sample 2.

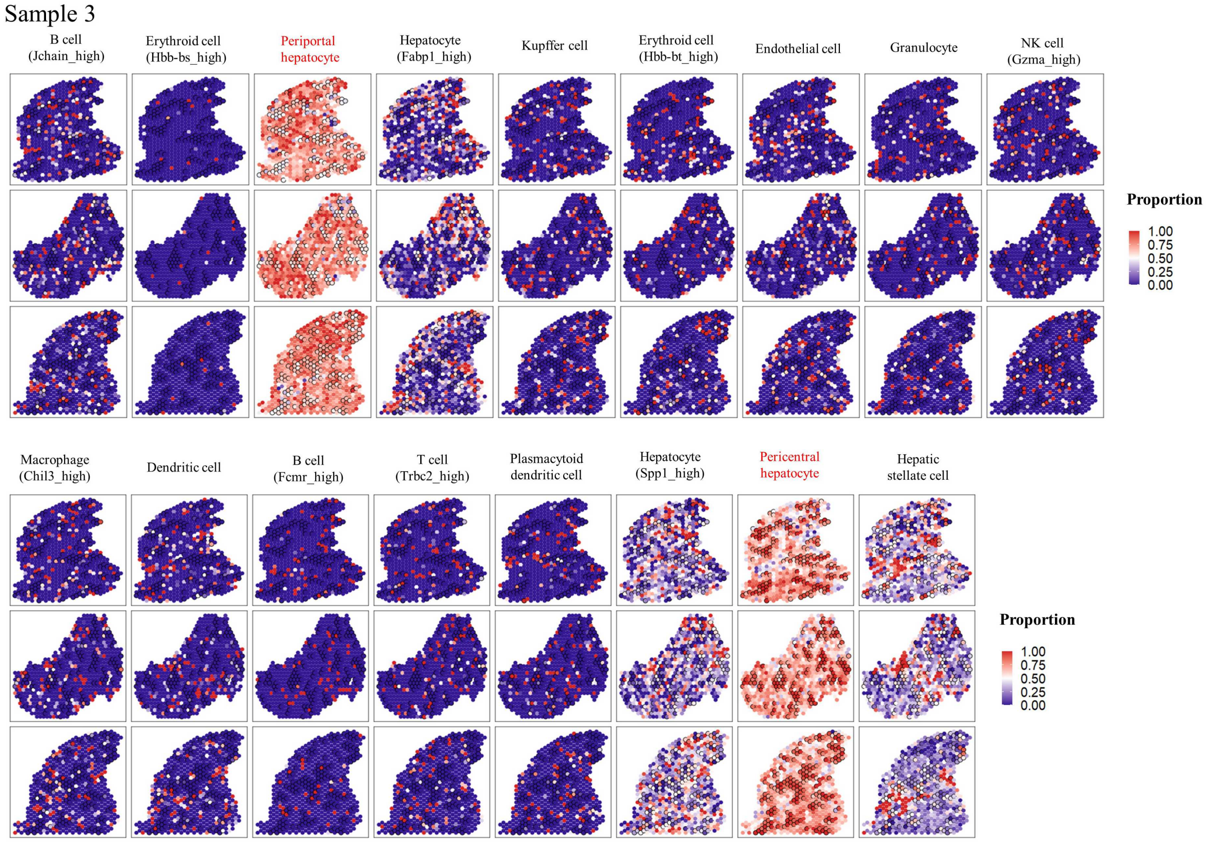


Figure S28: Cell-type deconvolution analysis for the eight mouse liver sections. Spatial heatmap of proportions of 17 cell types annotated in MCA on liver tissues for the three slices of Sample 3.

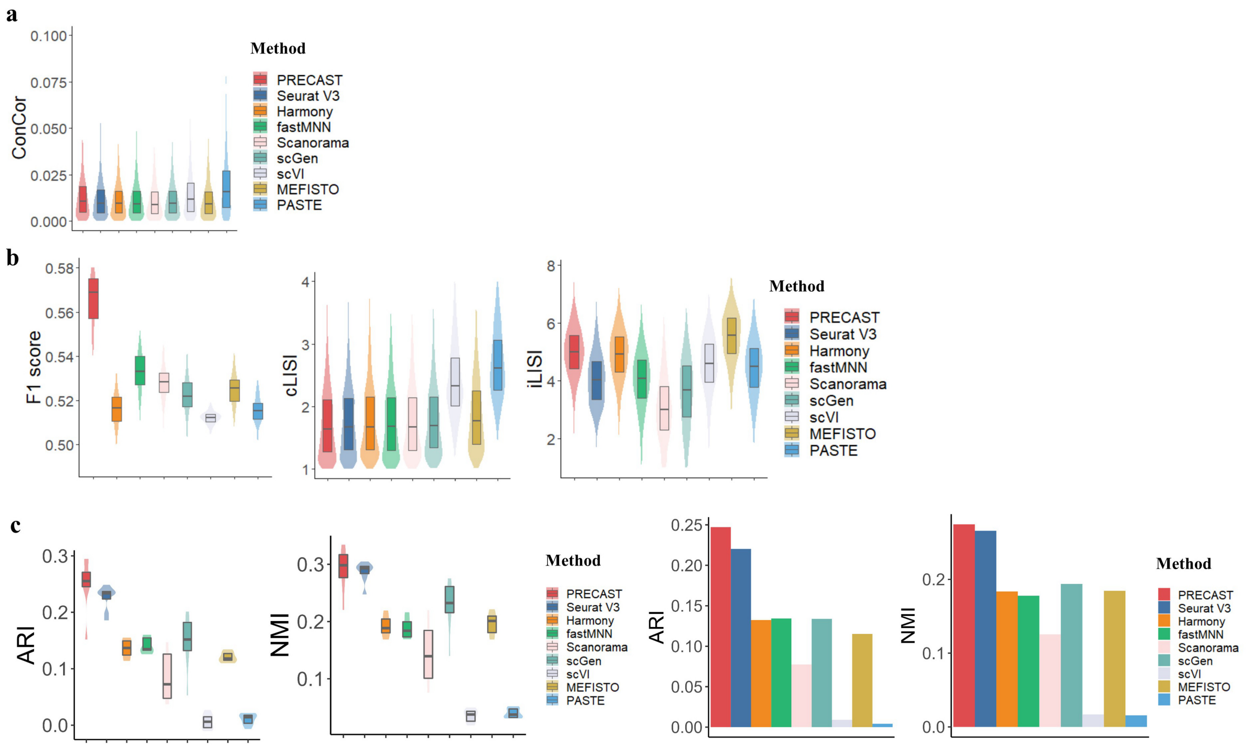


Figure S29: Dimension reduction with batch correction and clustering analysis using the top 2,000 HVGs for mouse liver ST data ($n = 4,865$ locations over 8 tissue sections). a. Boxplot/violin plot of conditional correlations from PRECAST and eight other methods. A lower conditional correlation score is better. b. Boxplots of F1 score (higher is better), cLISI (lower is better) and iLISI (higher is better) for PRECAST and eight other methods. c. Boxplot/violin plot of ARIs/NMIs (higher is better) for each sample from PRECAST and SC-MEB clustering based on the low-dimensional embeddings of other compared methods, and bar plot of ARIs/NMIs for 12 combined samples for PRECAST and other methods. In the boxplot, the center line and box lines denote the median, upper, and lower quartiles, respectively.

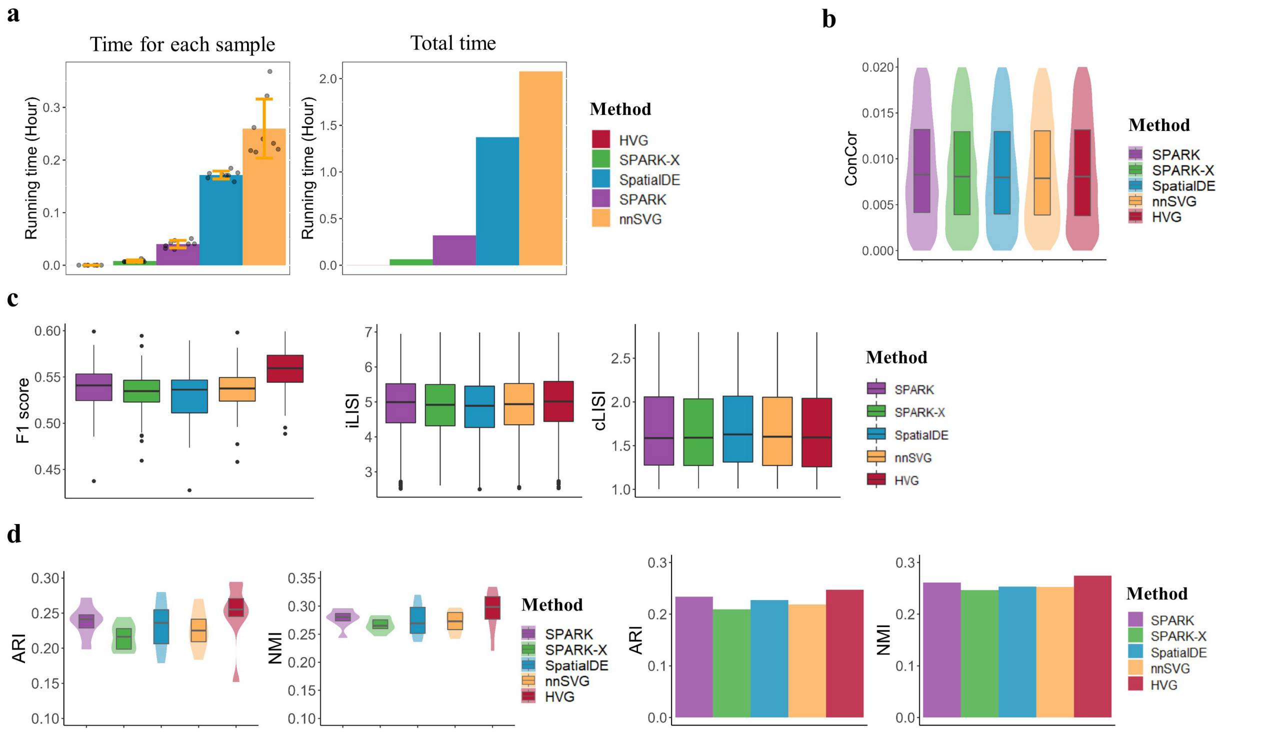


Figure S30: Comparison of PRECAST performance using five gene selection methods: SPARK, SPARK-X, SpatialDE, nnSVG and HVGs, to analyze mouse liver ST data ($n = 4,865$ locations over 8 tissue sections). a. Barplot of running times for each sample (left panel) and all samples (right panel). In the bar plot, the error bands represent the mean value ± standard deviation. b. Boxplot/violin plot of conditional correlations. c. Boxplot of F1 score, cLISI and iLISI. d. Boxplot/violin plot of ARIs/NMIs for each sample, and bar plot of ARIs/NMIs for the eight combined samples. In the boxplot, the center line, box lines and whiskers represent the median, upper, and lower quartiles, and 1.5 times interquartile range, respectively.

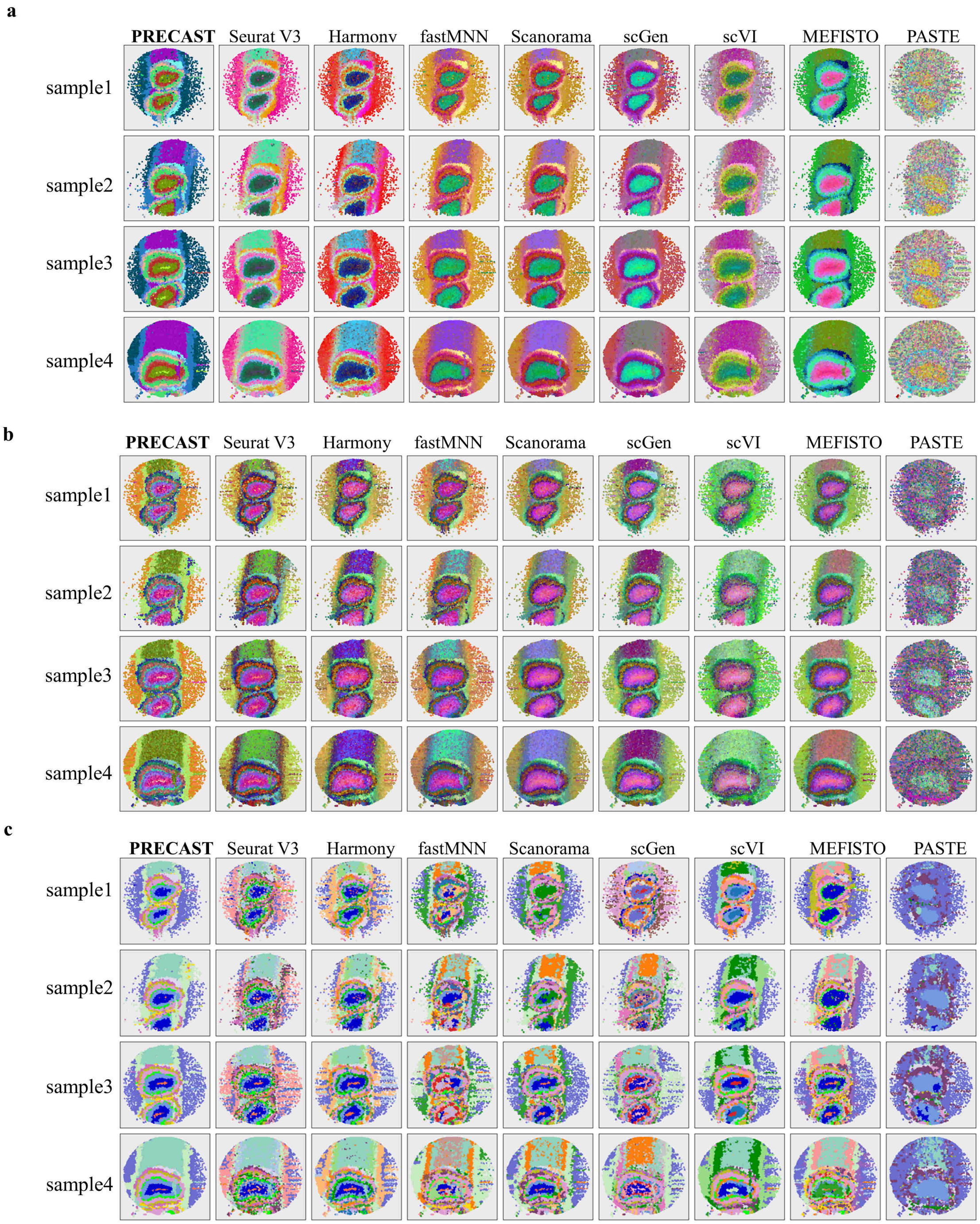


Figure S31: Dimension reduction with batch correction and clustering analysis of data for the 16 mouse olfactory bulb sections with reduced resolution. a. UMAP RGB plots for Samples 1-4 based on the low-dimensional embeddings from PRECAST and eight other methods. b. tSNE RGB plots for these methods. c. Domain clustering plots for these methods.

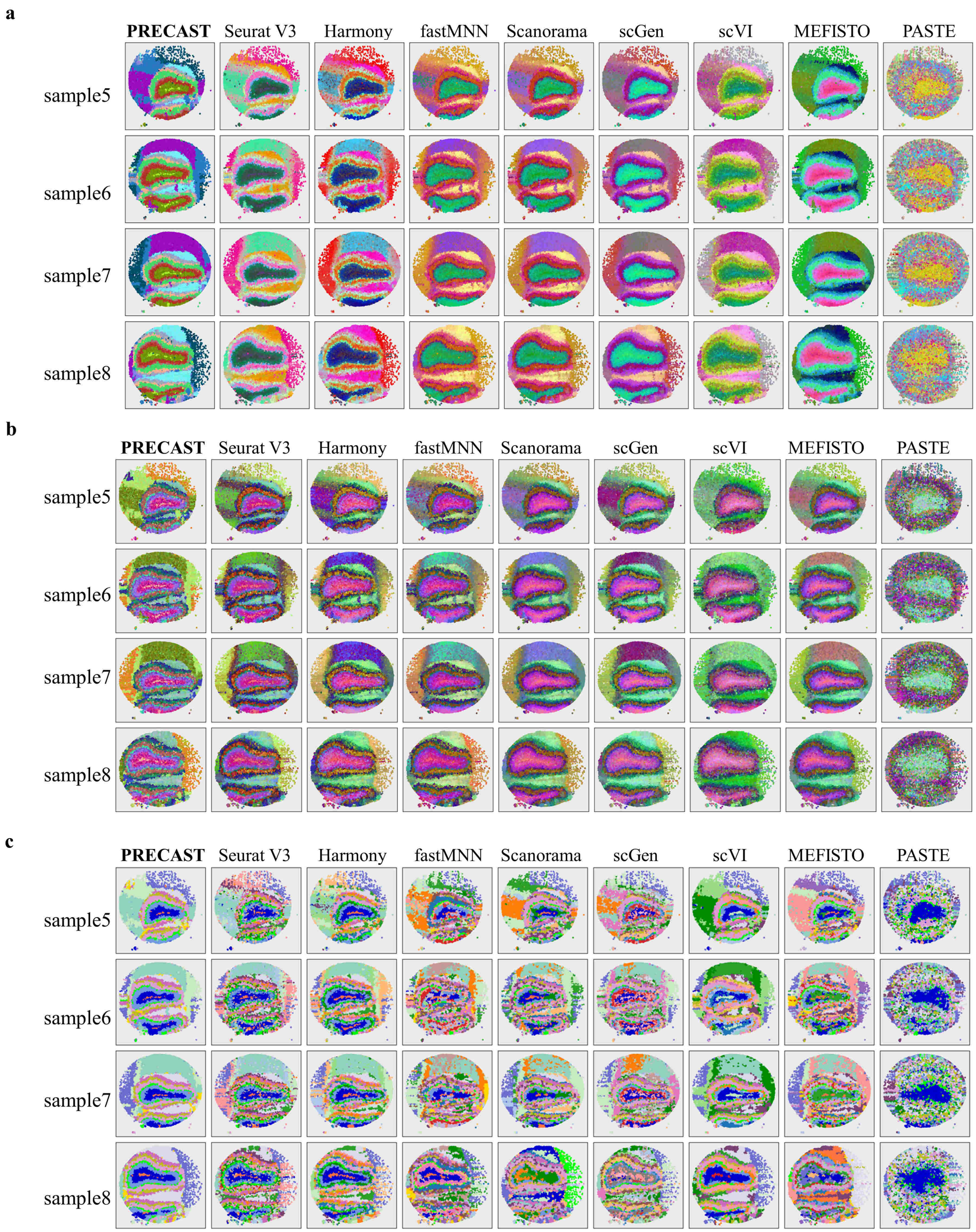


Figure S32: Dimension reduction with batch correction and clustering analysis of data for the 16 mouse olfactory bulb sections with reduced resolution. a. UMAP RGB plots for Samples 5-8 based on the low-dimensional embeddings from PRECAST and eight other methods. b. tSNE RGB plots for these methods. c. Domain clustering plots for these methods.

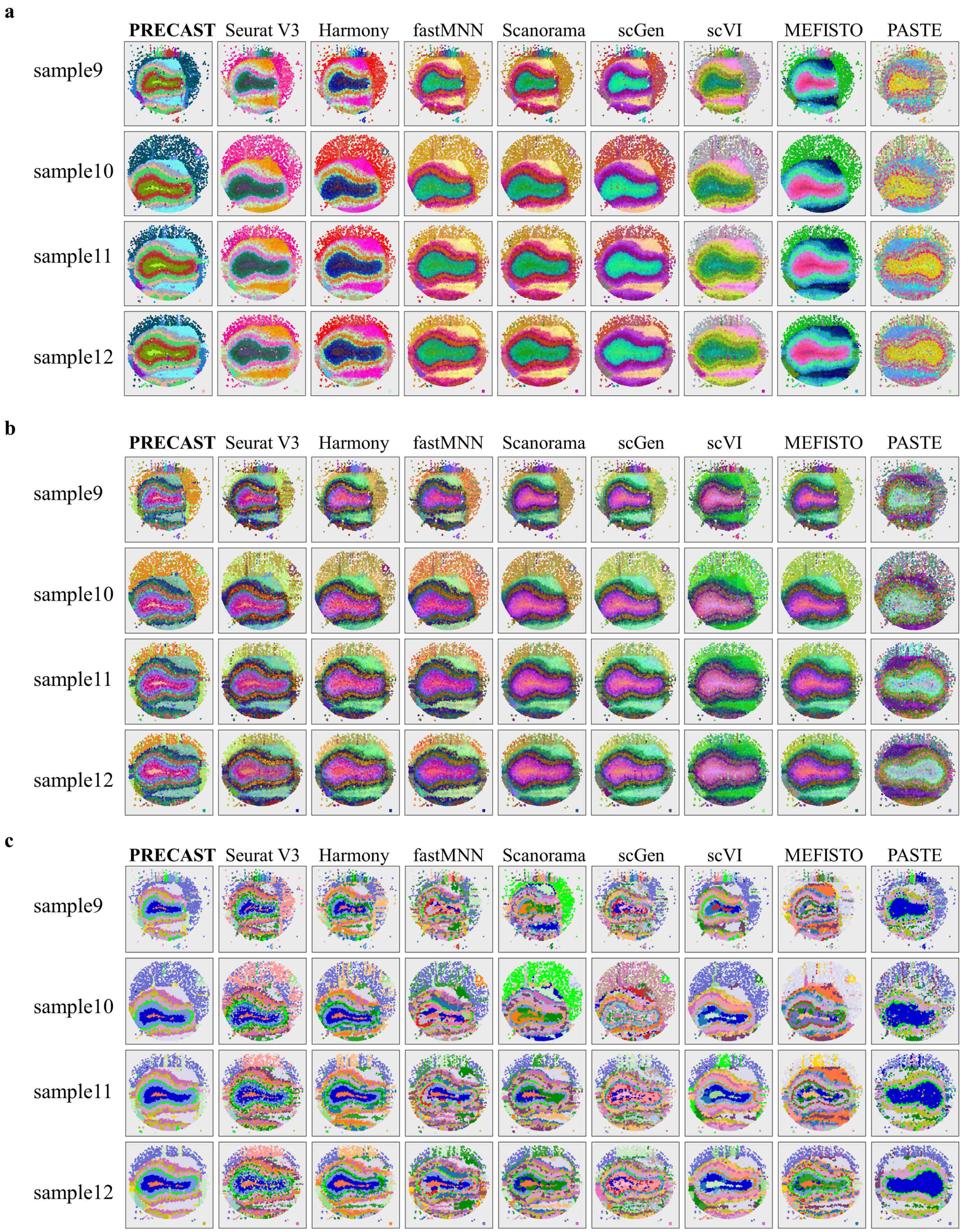


Figure S33: Dimension reduction with batch correction and clustering analysis of data for the 16 mouse olfactory bulb sections with reduced resolution. a. UMAP RGB plots for samples 9-12 based on the low-dimensional embeddings from PRECAST and eight other methods. b. tSNE RGB plots for these methods. c. Domain clustering plots for these methods.

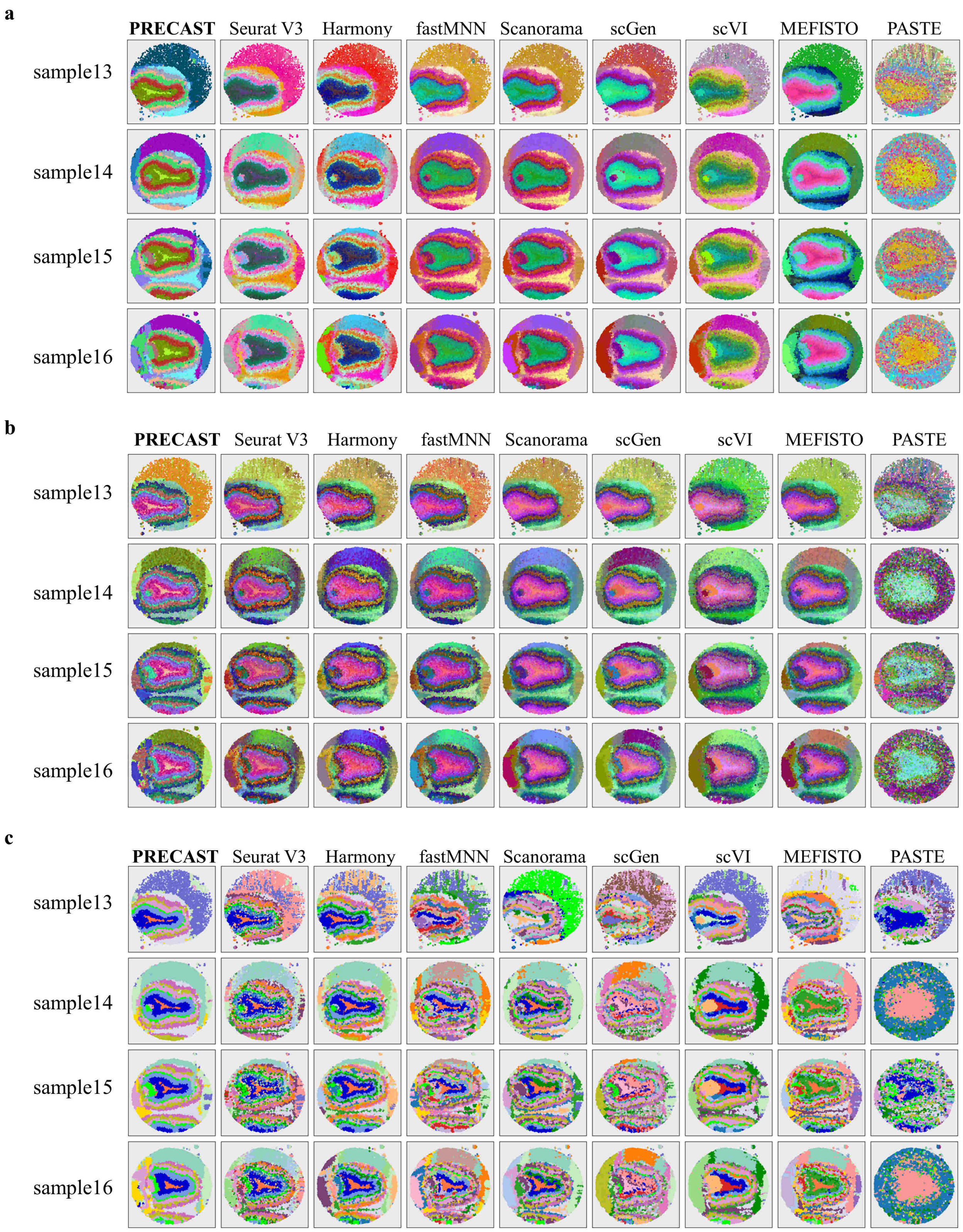


Figure S34: Dimension reduction with batch correction and clustering analysis of data for the 16 mouse olfactory bulb sections with reduced resolution. a. UMAP RGB plots for samples 13-16 based on the low-dimensional embeddings from PRECAST and eight other methods. b. tSNE RGB plots for these methods. c. Domain clustering plots for these methods.

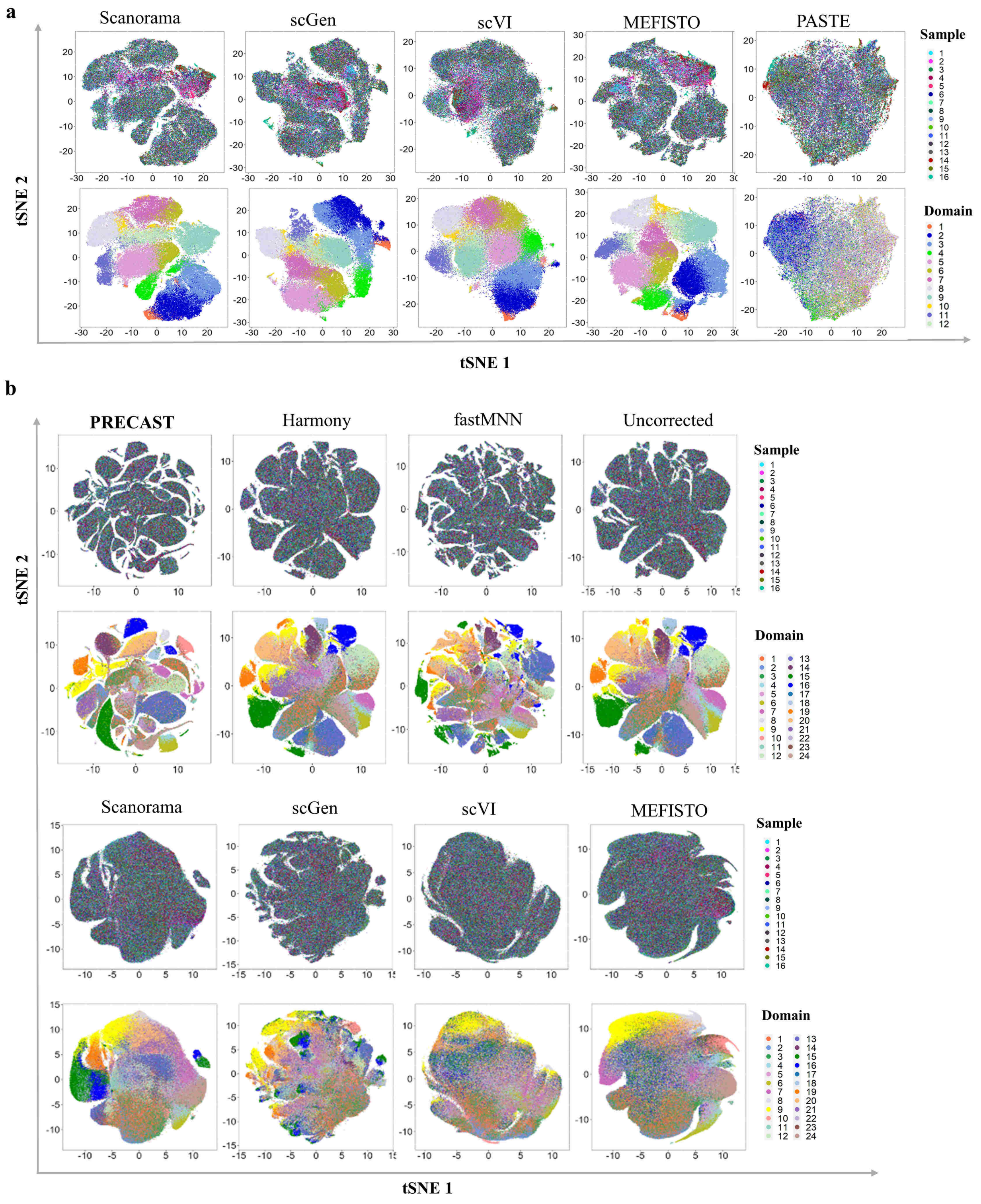


Figure S35: Batch correction analysis of data for the 16 mouse olfactory bulb sections. a. tSNE plots of five other compared integration methods (Scanorama, scGen, scVI, MEFISTO and PASTE) for resolution-reduced data; tSNE plot is colored for sample index and domain index. b. tSNE plots for PRECAST and eight other integration methods, as well as the uncorrected tSNE for the original-resolution data. It is noted that Seurat V3 and PASTE can not be implemented for this data due to the large number of spots.

**a**

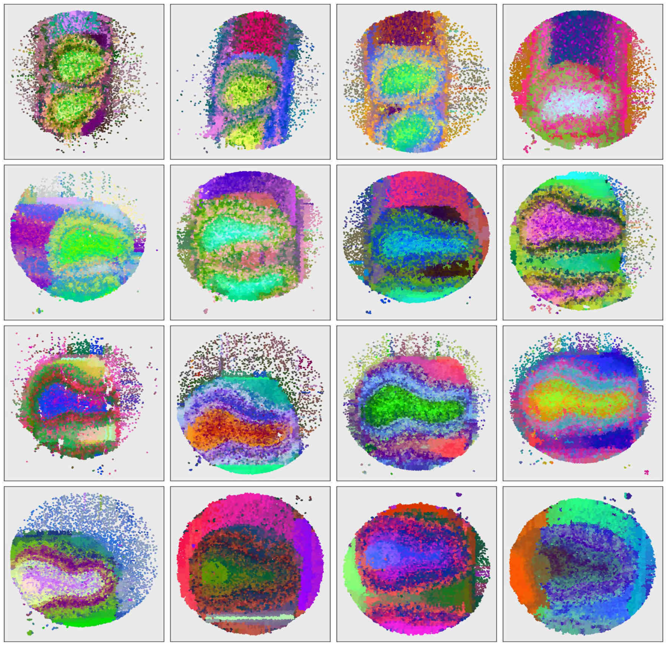

**b**

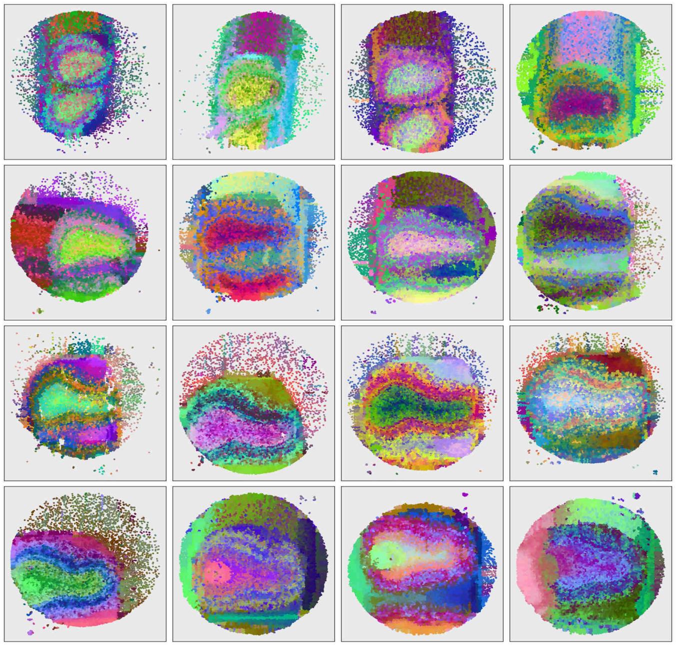

Figure S36: Microenvironment spatial dependence analysis of data for the 16 mouse olfactory bulb sections with reduced resolution. a. UMAP RGB plot using the inferred embeddings from the intrinsic CAR of PRECAST for the 16 samples. First row, Samples 1-4; second row, Samples 5-8; third row, Samples 9-12; fourth row, Samples 13-16. b. tSNE RGB plot using the inferred embeddings from the intrinsic CAR of PRECAST for the 16 samples.

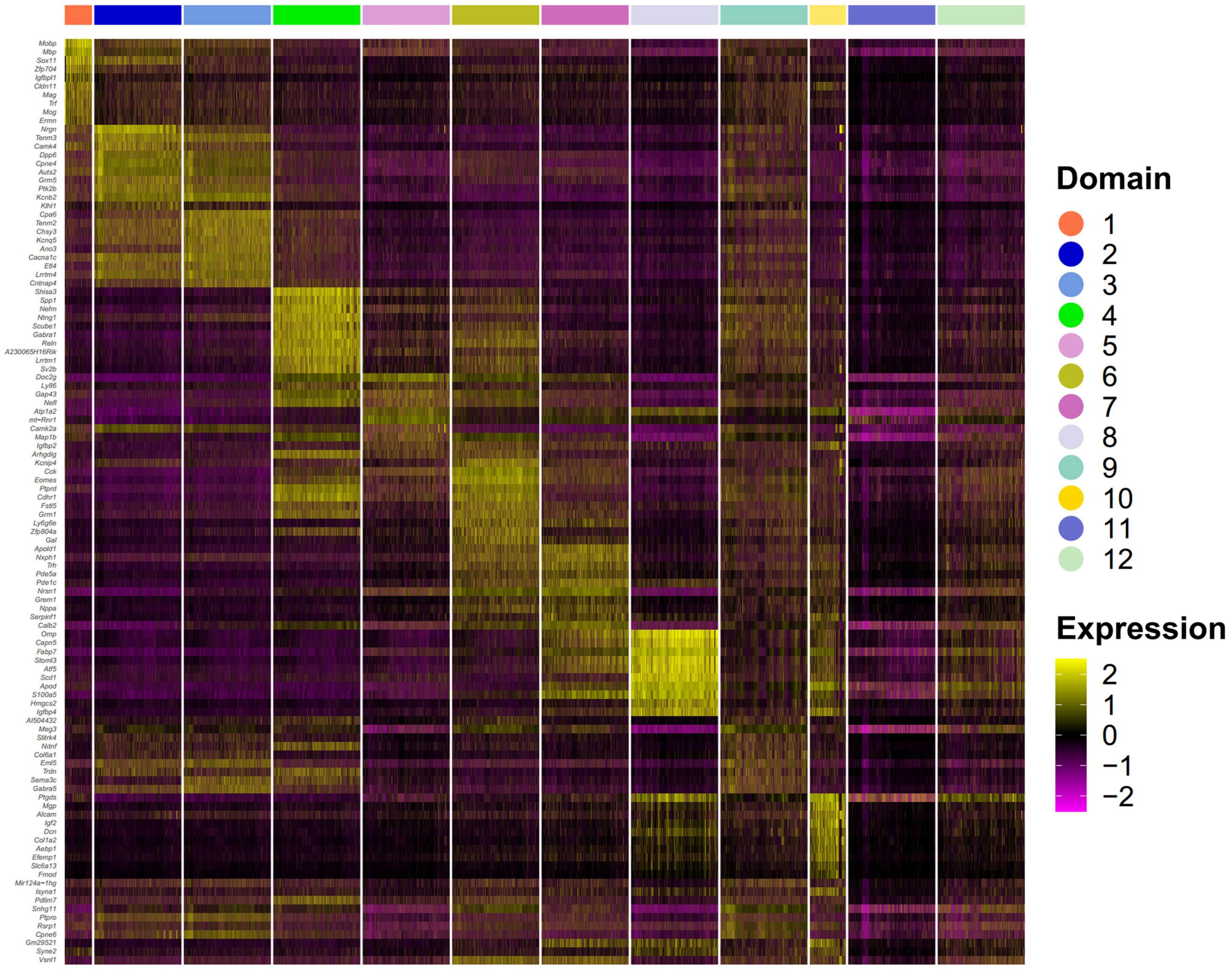


Figure S37: Combined DE analysis of data for the 16 mouse olfactory bulb sections with reduced resolution. Heatmap of top 10 DE genes for each domain identified by PRECAST.

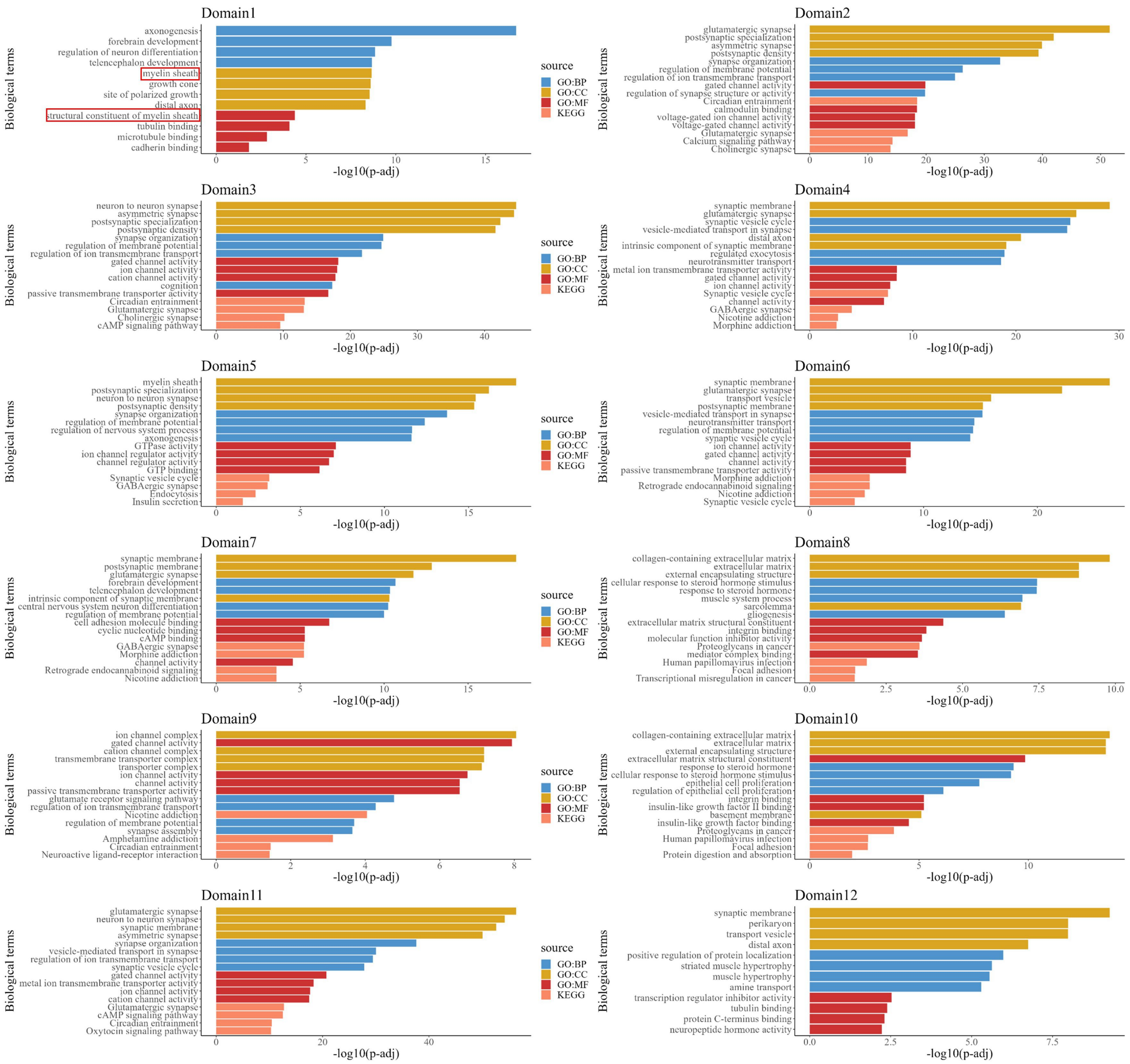


Figure S38: Enrichment analysis of data for the 16 mouse olfactory bulb sections with reduced resolution. Top 4 significant pathways (term size <500) for each category of DE genes for Domains 1-12 identified by PRECAST. The $p$-values are based on one-sided Fisher's exact tests with multiple testing corrections using the Benjamini-Hochberg FDR method.

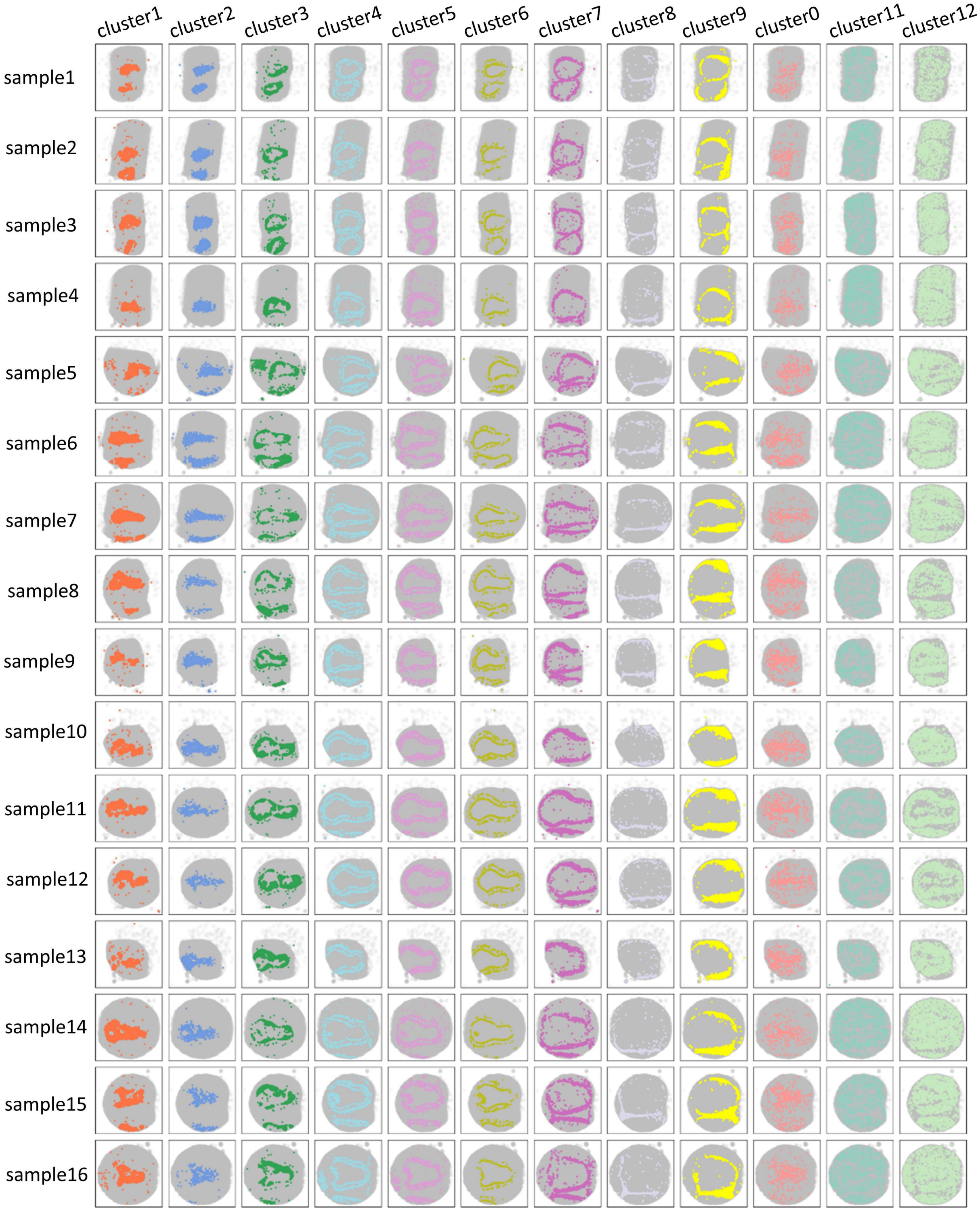


Figure S39: Spatial clustering analysis of data for the 16 mouse olfactory bulb sections with original resolution. Each sample is arranged by row, and clusters (1-12) obtained by PRECAST are sorted by column.

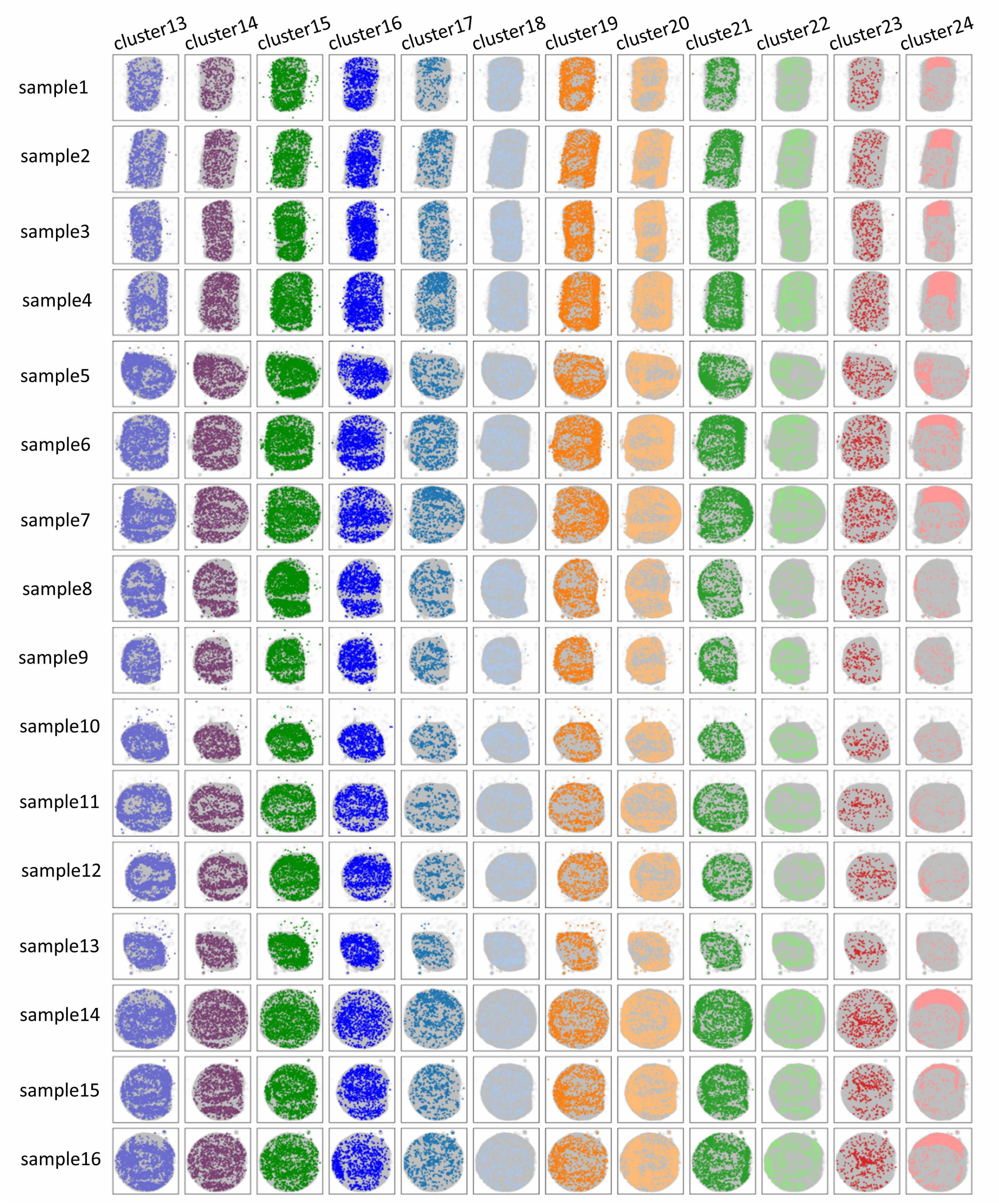


Figure S40: Spatial clustering analysis of data for the 16 mouse olfactory bulb sections with original resolution. Each sample is arranged by row, and clusters (13 -24) obtained by PRECAST are sorted by column.

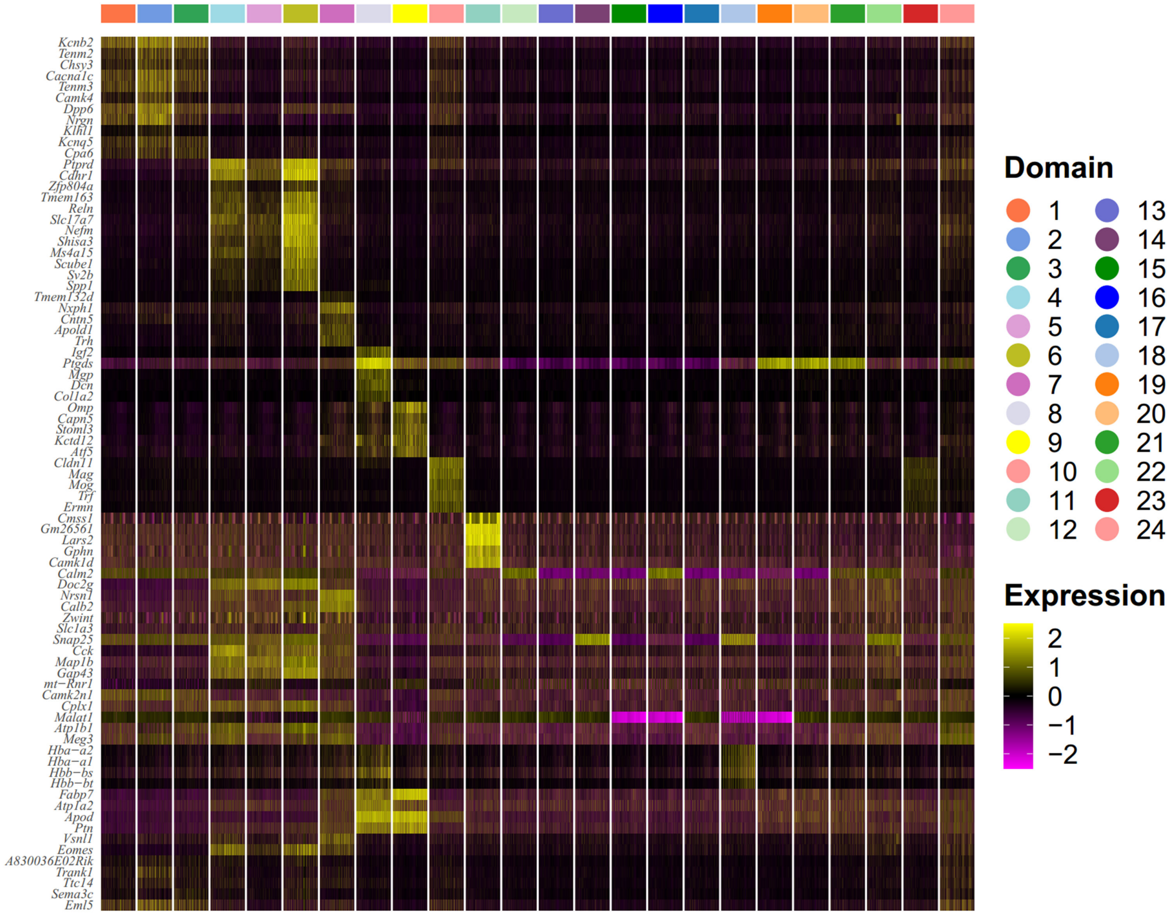


Figure S41: Combined DE analysis of data for the 16 mouse olfactory bulb sections with original resolution. Heatmap of top 5 DE genes among all clusters identified by PRECAST.

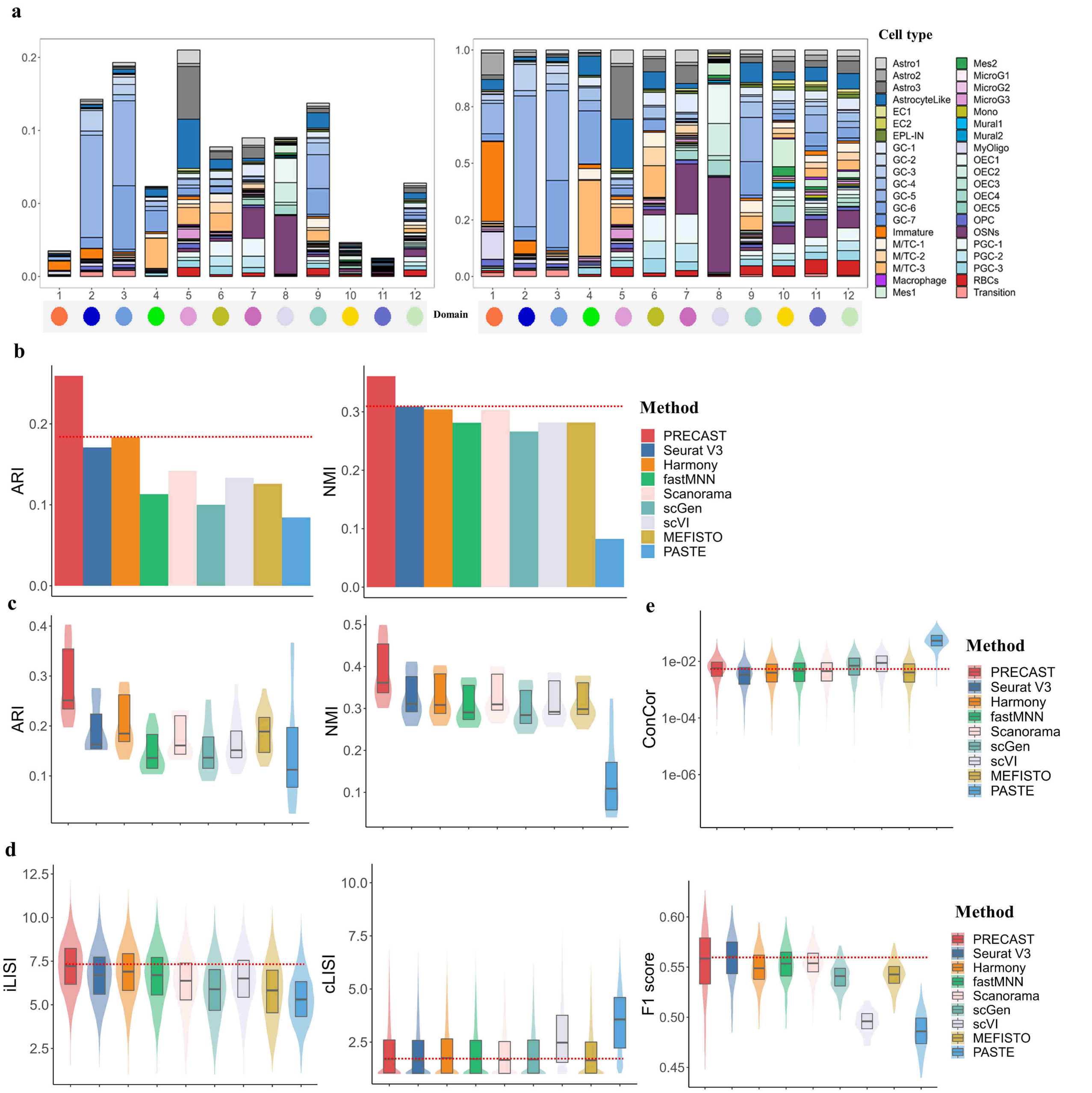


Figure S42: Cell-type deconvolution analysis of data for the 16 mouse olfactory bulb sections with reduced resolution ($n = 594,890$ locations over 16 tissue sections). a. Percentage of different cell types in each domain detected by PRECAST, with (left panel) or without (right panel) scaling to the summation of all cell types across all domains equal to 100%. b. Bar plot of ARIs/NMIs for 16 combined samples from PRECAST and SC-MEB clustering based on the low-dimensional embeddings of other compared methods. c. Boxplot/violin plot of ARIs/NMIs for each sample from PRECAST and other methods. d. Boxplot/violin of iLISI, cLISI and F1 score (F1 score of the average silhouette coefficients) for PRECAST and eight other methods. e. Boxplot/violin plot of conditional correlations from PRECAST and eight other methods. In the boxplot, the center line and box lines denote the median, upper, and lower quartiles, respectively.

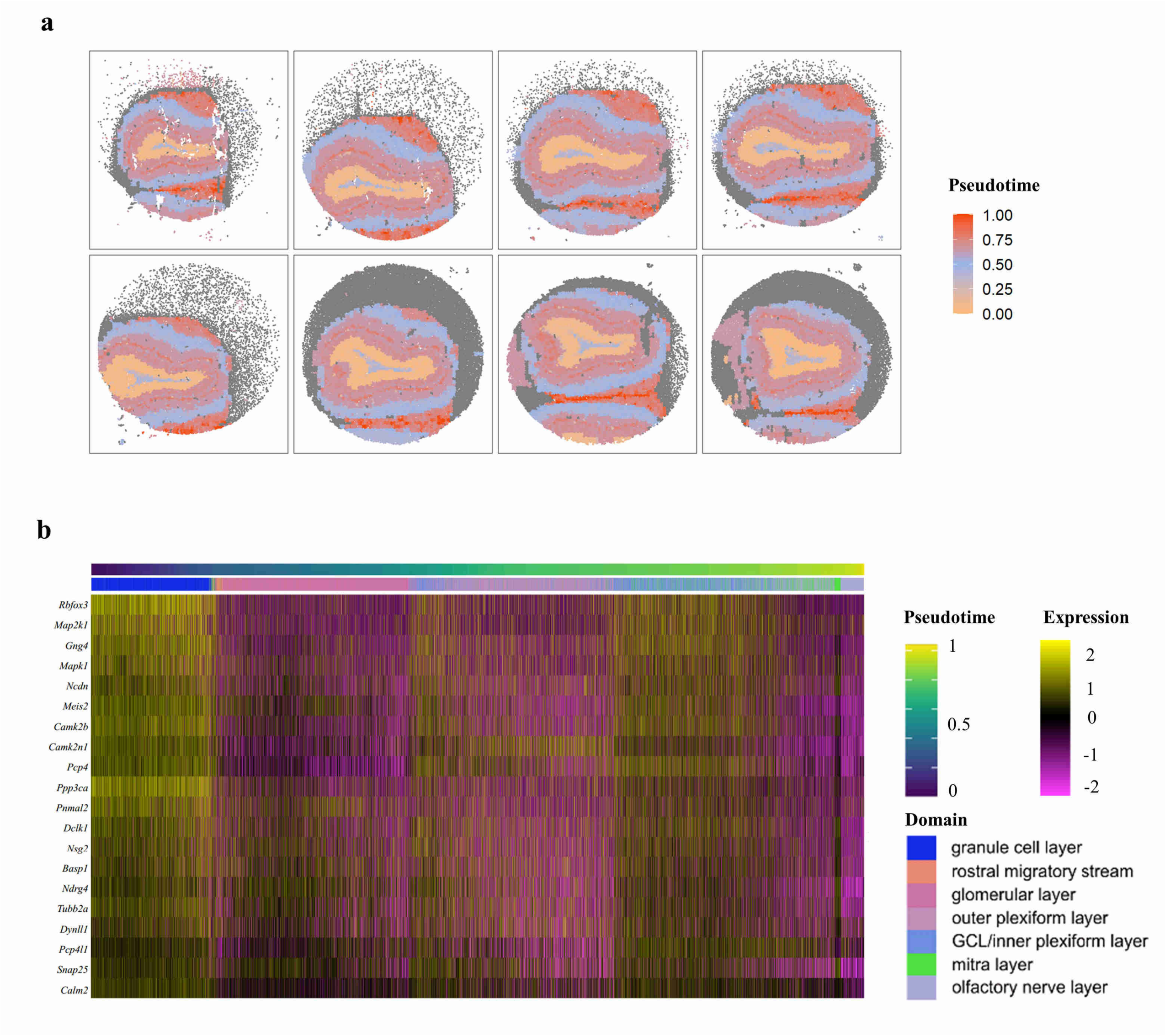


Figure S43: Combined trajectory inference of data for the 16 mouse olfactory bulb sections with reduced resolution. a. Visualization of the trajectory inferred by PRECAST in spatial heatmap of Samples 9-16; the first row shows Samples 9-12, and the second row shows Samples 13-16. b. Heatmap of top 20 differentially expressed genes along pseudotime.

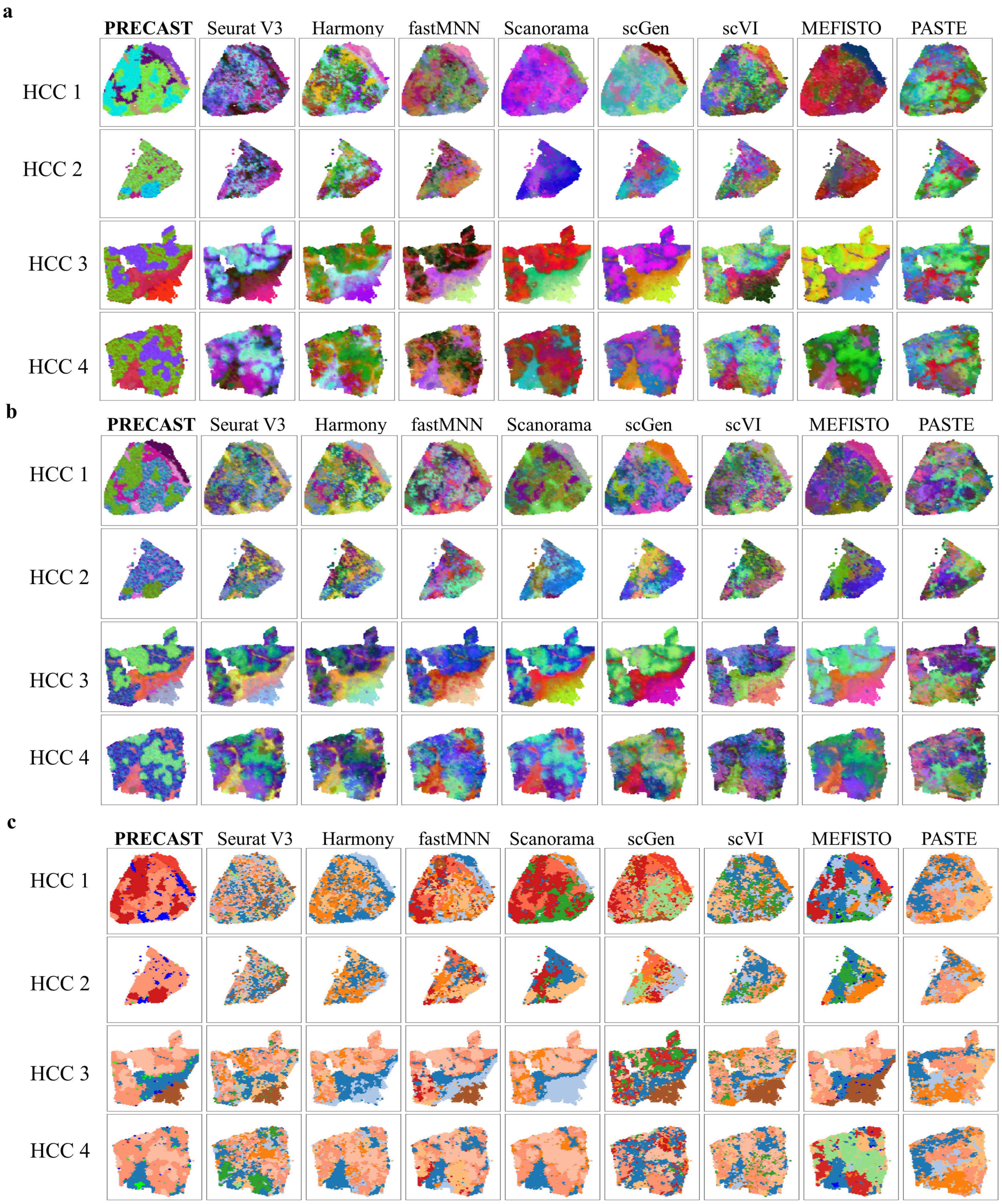


Figure S44: Dimension reduction with batch correction and clustering analysis of data for the four Human HCC sections. a. UMAP RGB plot based on low-dimensional embeddings from PRECAST and eight other methods. b. tSNE RGB plots for the methods. c. Domain clustering plots for the methods, where SC-MEB is used in the other methods for clustering based on their low-dimensional embeddings.

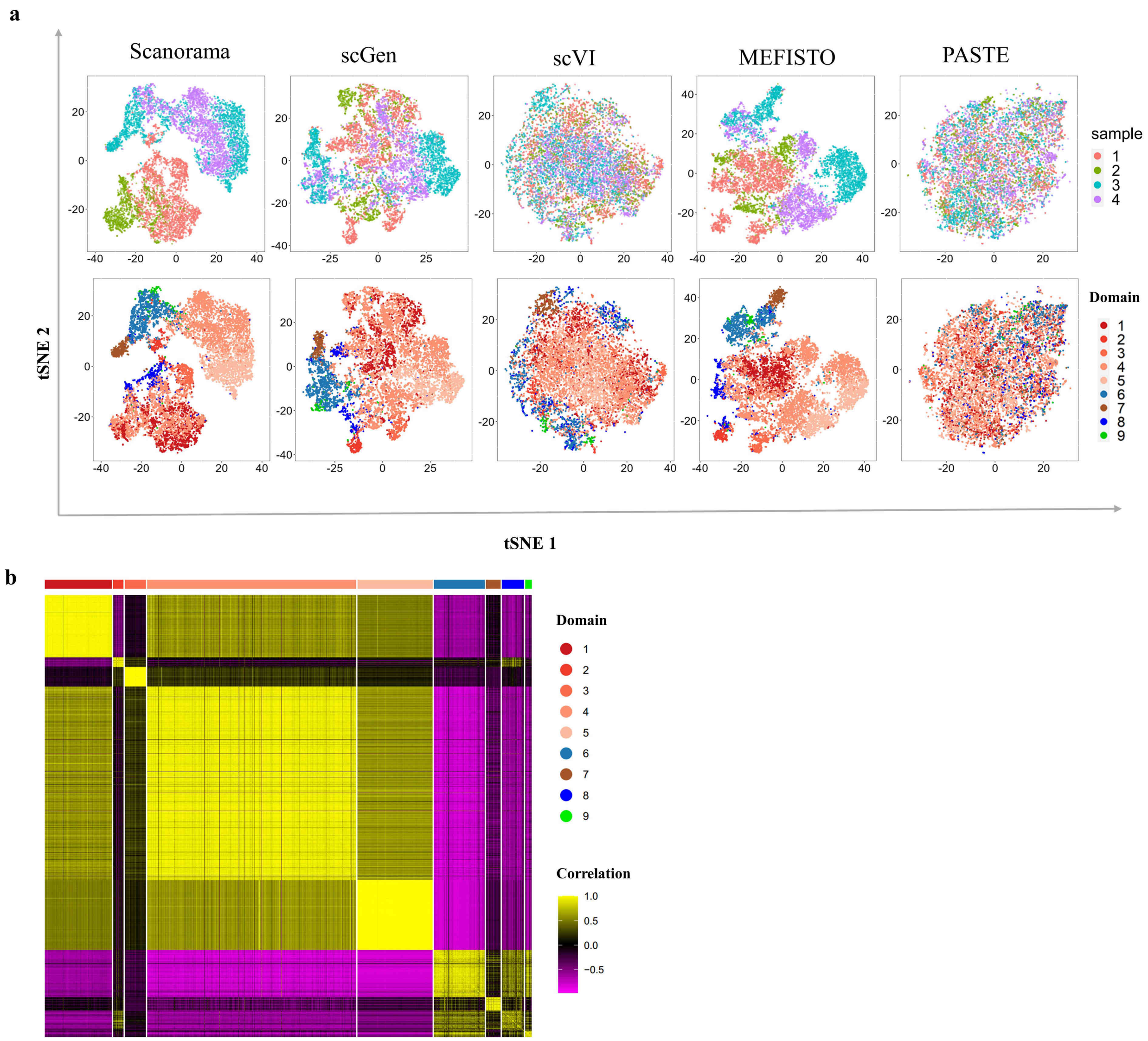


Figure S45: Batch correction analysis and correlation analysis of data for the four Human HCC sections. a. Top panel: visualization of batches based on two-dimensional tSNE embeddings from five different integration methods including Scanorama, scGen, scVI, MEFISTO and PASTE; bottom panel: visualization of the cluster labels estimated by PRECAST based on the same tSNE embeddings. b. Heatmap of correlations of batch-corrected low-dimensional embeddings estimated by PRECAST.

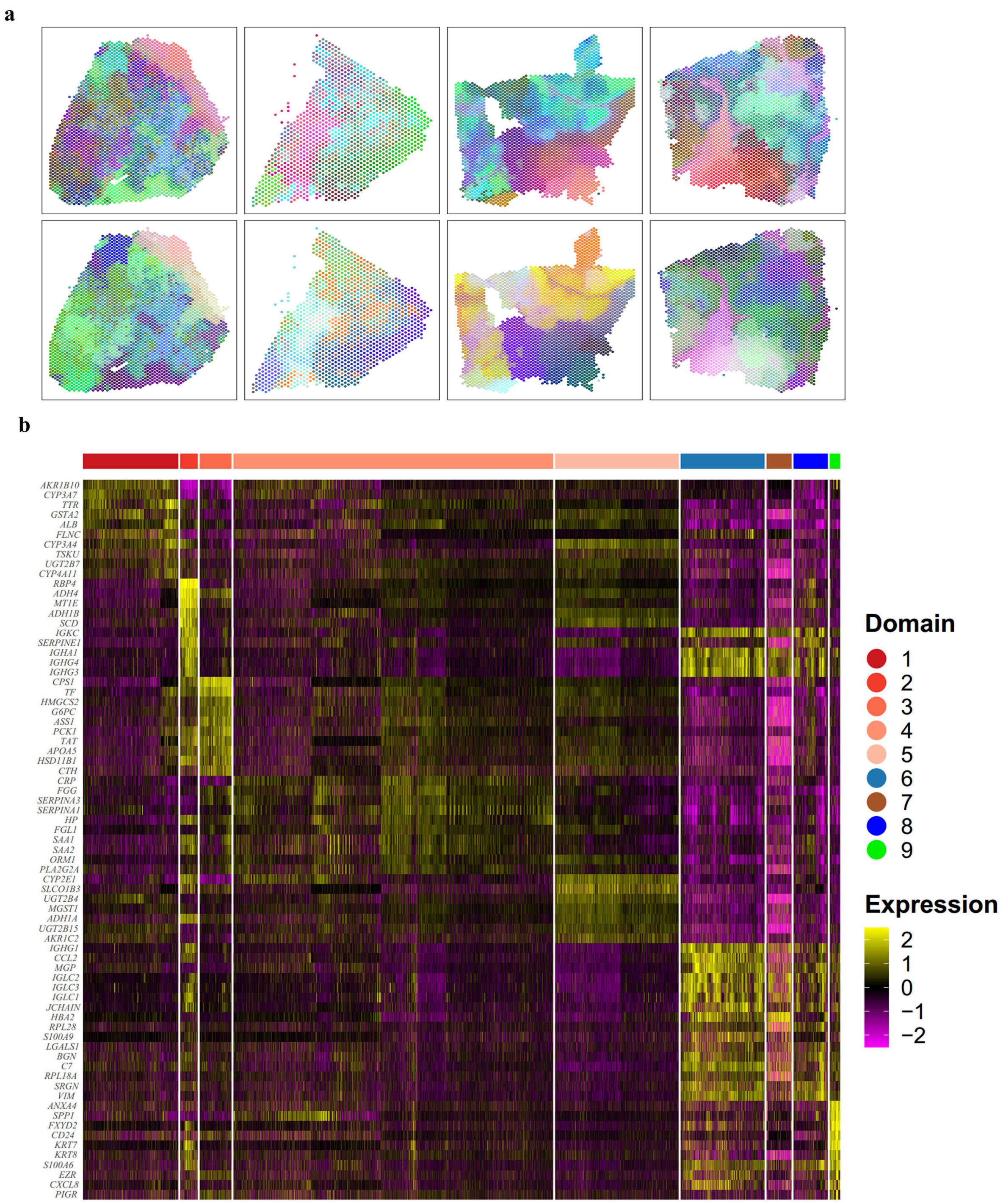


Figure S46: Microenvironment spatial dependence analysis and combined DE analysis of data for the four Human HCC sections. a. UMAP /tSNE RGB plot of the inferred embeddings from the intrinsic CAR of PRECAST for the four samples. b. Heatmap of top 10 DE genes for each domain identified by PRECAST.

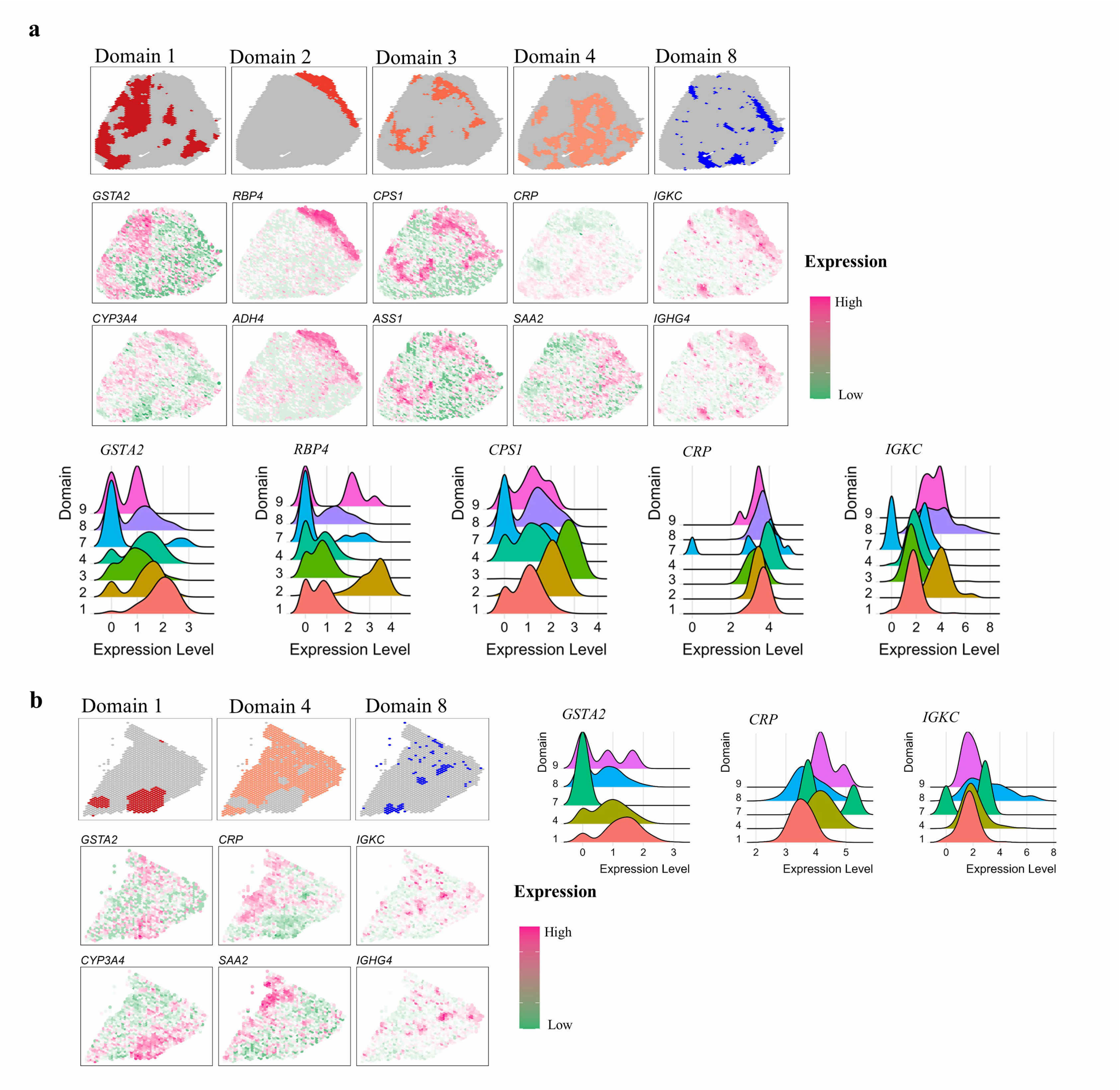


Figure S47: Combined DE analysis of data for the four Human HCC sections. a. First row: spatial heatmap of spatial domains 1–4 and 8 of HCC1 detected by PRECAST. Second/Third row: spatial heatmap of scaled expression of top two DE genes in Domains 1–4, and 8 of HCC1. Bottom row: Ridge plots of log-normalized gene expression of top DE gene in each of Domains 1–4 and 8 of HCC1. b. First row: spatial heatmap of spatial domains 1, 4 and 8 of HCC2 detected by PRECAST. Second/Third row: spatial heatmap of scaled expression of top two DE genes in Domains 1, 4 and 8 of HCC2. Bottom row: Ridge plots of log-normalized gene expression of top DE gene in each of Domains 1, 4 and 8 of HCC2.

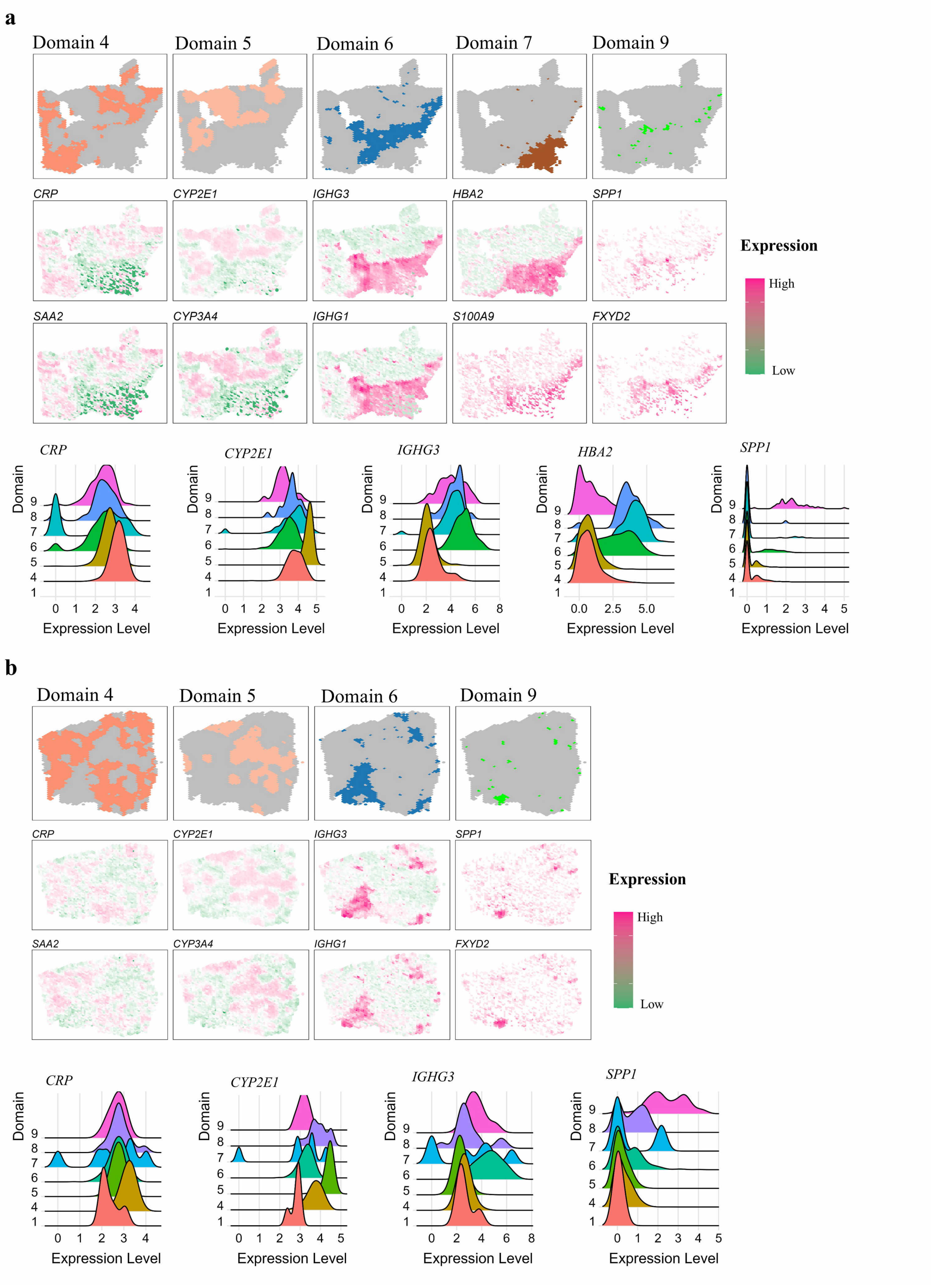


Figure S48: Combined DE analysis of data for the four Human HCC sections. a. First row: spatial heatmap of spatial domains 4–7 and 9 of HCC3 detected by PRECAST. Second/Third row: spatial heatmap of scaled expression of top two DE genes in Domains 4–7 and 9 of HCC3. Bottom row: Ridge plots of log-normalized gene expression of top DE gene in each of Domains 4–7 and 9 of HCC3. b. First row: spatial heatmap of spatial domains 4–6 and 9 of HCC4 detected by PRECAST. Second/Third row: spatial heatmap of scaled expression of top two DE genes in Domains 4–6 and 9 of HCC4. Bottom row: Ridge plots of log-normalized gene expression of top DE gene in each of Domains 4–6 and 9 of HCC4.

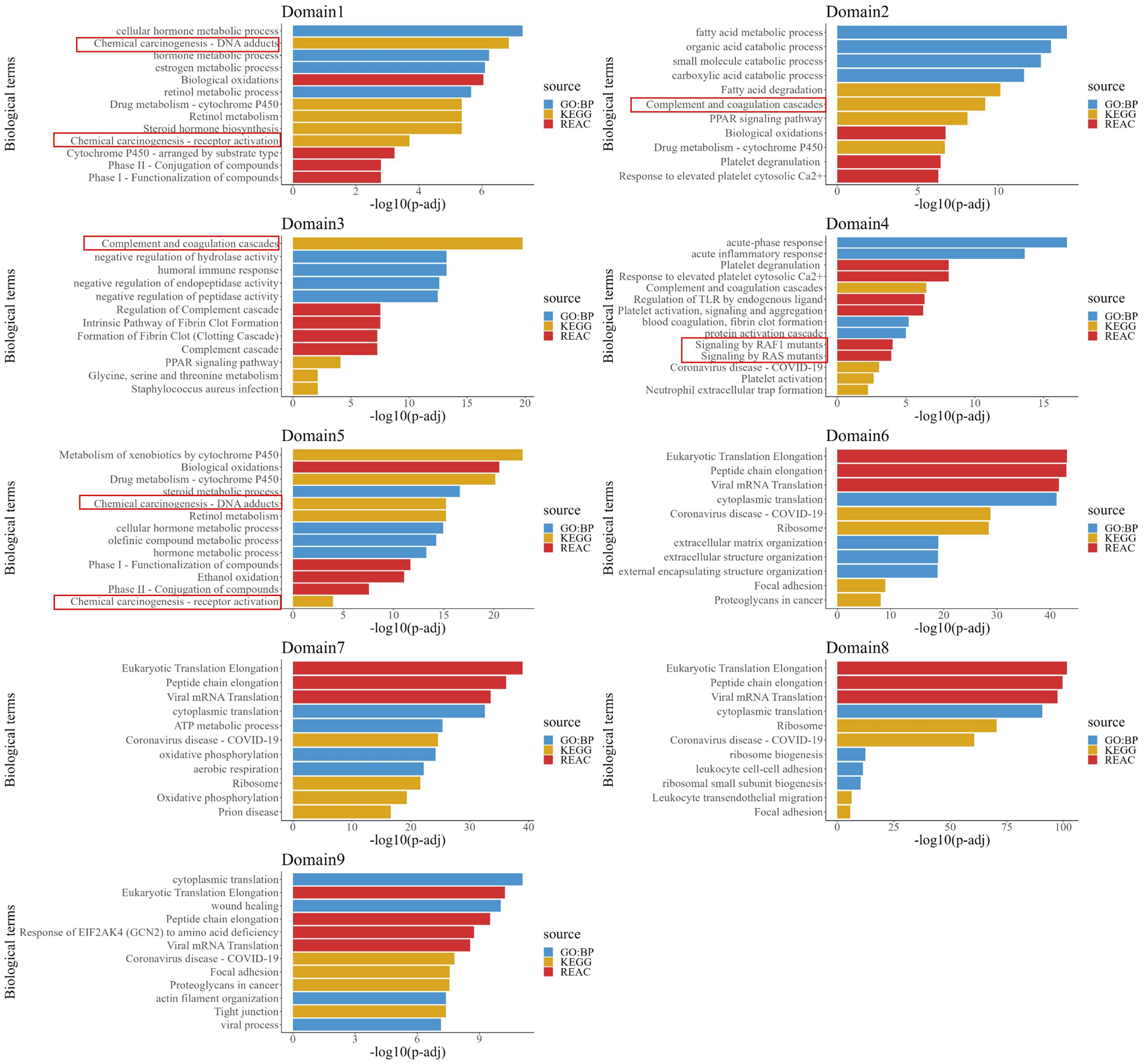


Figure S49: Enrichment analysis of DE genes in data for each domain detected by PRECAST in the four Human HCC sections. Top 4 pathways for each category, as well as some additional interesting pathways for Domains 1, 4, and 5. The $p$-values are based on one-sided Fisher's exact tests with multiple testing corrections using the Benjamini-Hochberg FDR method.

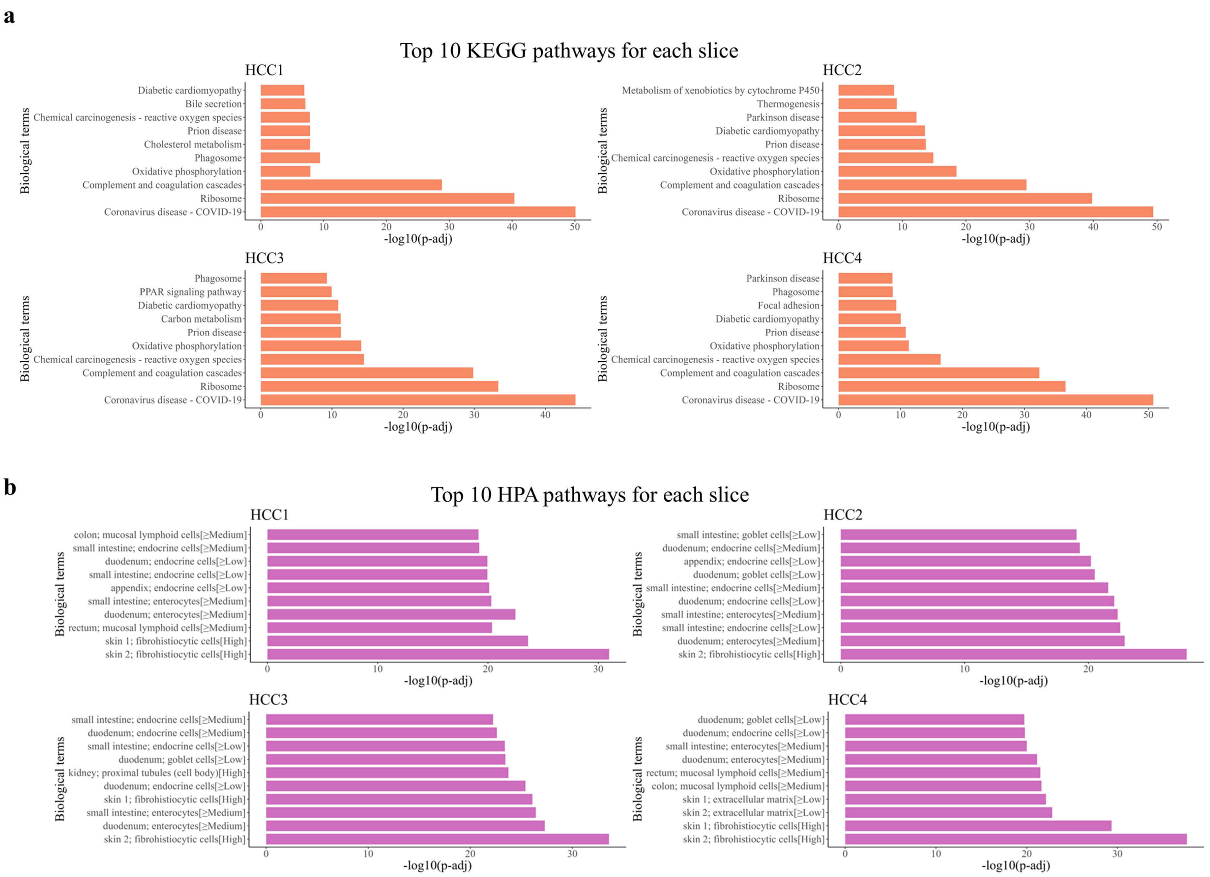


Figure S50: Enrichment analysis of spatially variable genes while controlling the domain-related low-dimensional embeddings obtained by PRECAST in the four human HCC sections. Bar plot of top 10 significant KEGG pathways (term size $< 500$). b. Bar plot of top 10 significant HPA pathways (term size $< 500$). The $p$-values are based on one-sided Fisher's exact tests with multiple testing corrections using the Benjamini-Hochberg FDR method.

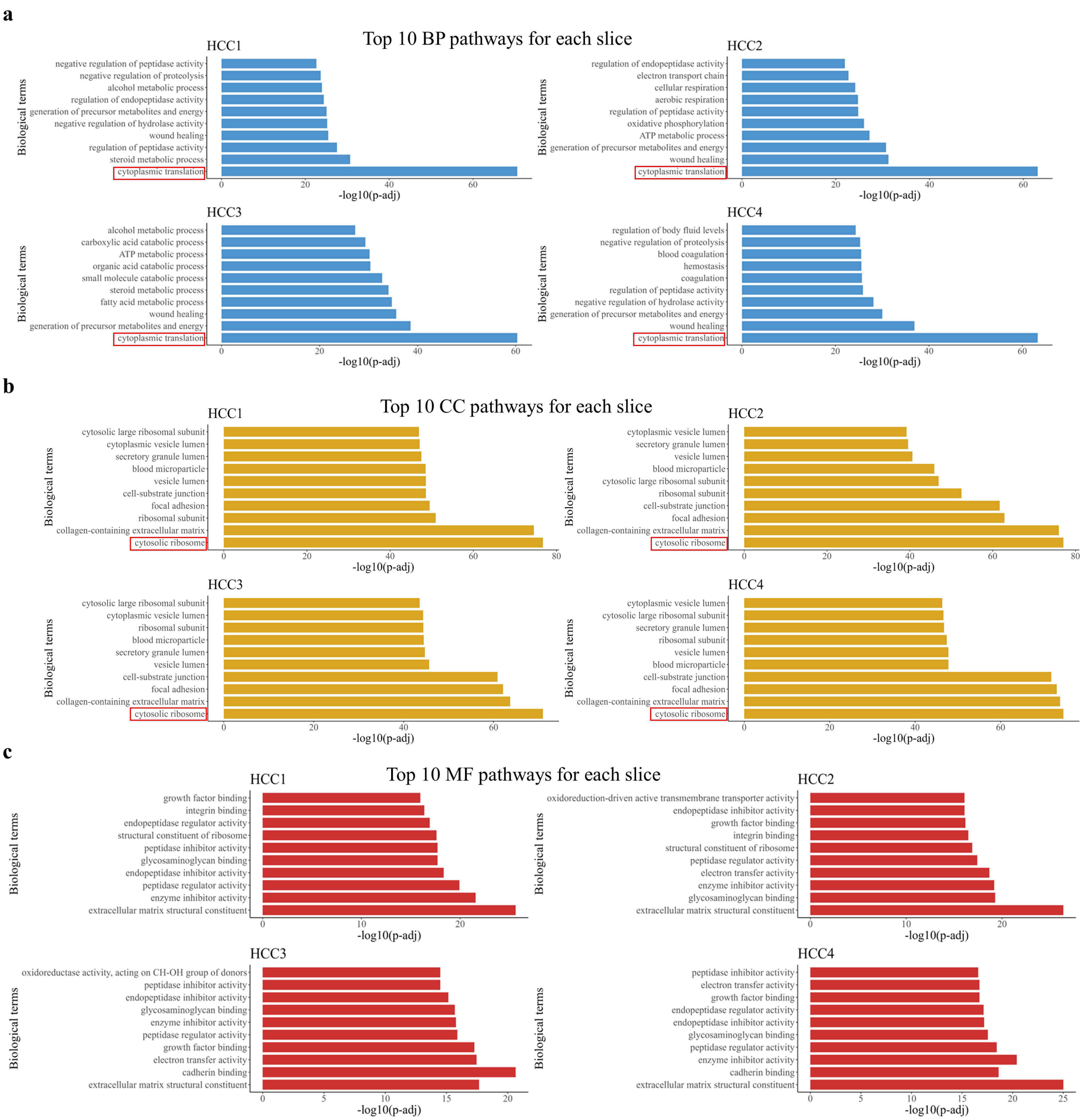


Figure S51: GO enrichment analysis of spatially variable genes while controlling the domain-related low-dimensional embeddings obtained by PRECAST in the four human HCC sections. a. Bar plot of top 10 significant BP pathways (term size $<$ 500). b. Bar plot of top 10 significant CC pathways (term size $<$ 500). c. Bar plot of top 10 significant MF pathways (term size $<$ 500). The $p$-values are based on one-sided Fisher's exact tests with multiple testing corrections using the Benjamini-Hochberg FDR method.

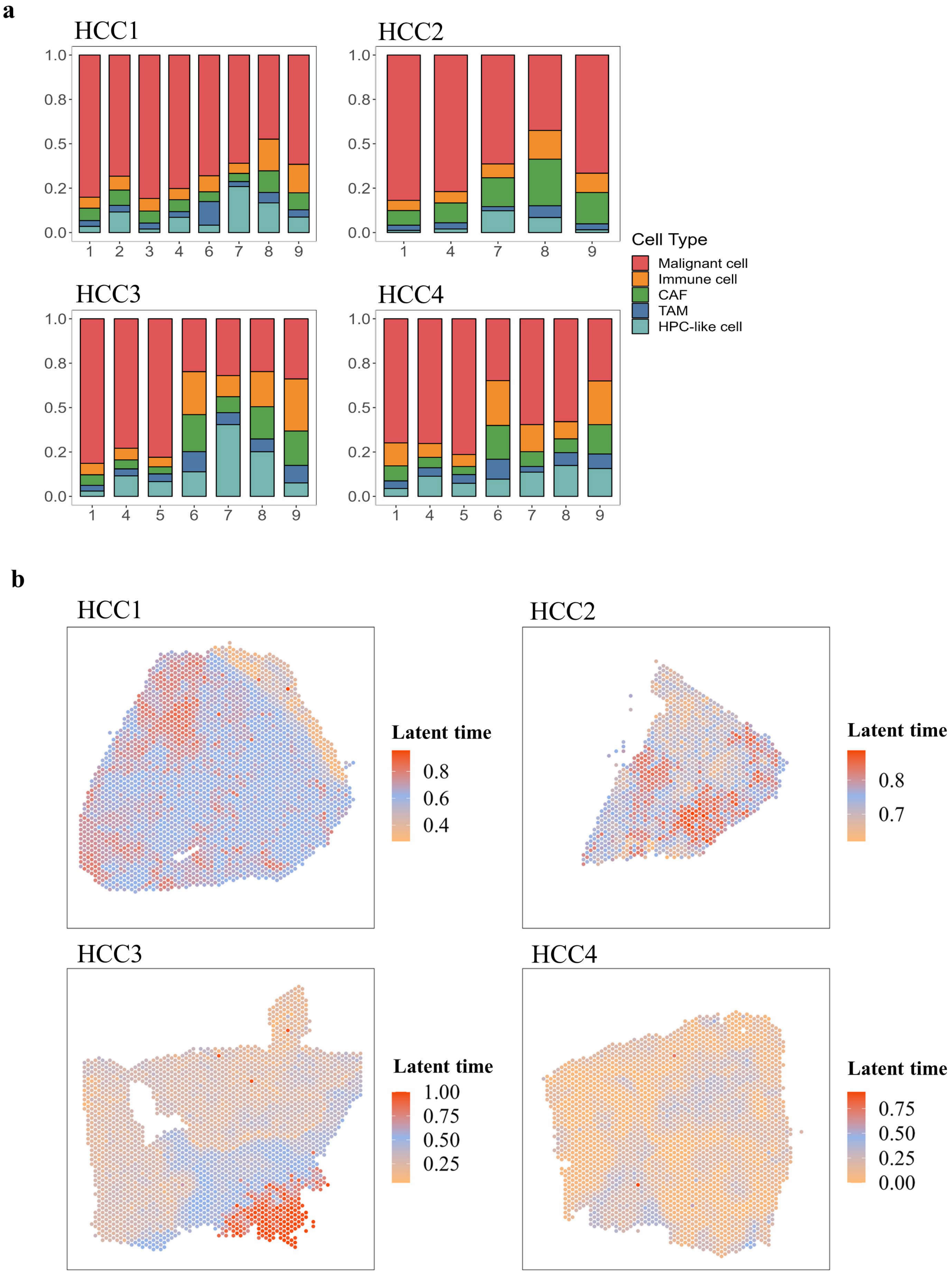


Figure S52: Cell-type deconvolution analysis and RNA velocity analysis of the four human HCC sections. a. Percentage of different cell types in each domain detected by PRECAST. b. Heatmap of RNA velocity in latent time in the spatial coordinates for the four HCC samples.
